# Local and network neural activations and their associations with sleep parameters during threat conditioning and extinction in persons with Generalized Anxiety Disorder with and without Insomnia Disorder

**DOI:** 10.1101/2025.11.11.687905

**Authors:** Jeehye Seo, Cagri Yuksel, Katelyn I. Oliver, Carolina Daffre, Huijin Song, Natasha B. Lasko, Emma R. S. McCoy, Mohammed R. Milad, Byoung-Kyong Min, Edward F. Pace-Schott

## Abstract

Deficient extinction learning and memory are hypothesized mechanisms for pathological anxiety that are associated with sleep disturbance. fMRI neural activations to threat conditioning, extinction learning, and extinction recall were measured. Activations were compared, in persons with Generalized Anxiety Disorder (GAD), between those with moderate to severe Insomnia Disorder (ID) and those with absent or sub-threshold ID. Relationships of activations with measures of sleep quality and physiology were examined. Between-group comparisons and whole-sample correlation with sleep parameters were examined in relation to large-scale brain networks using a liberal cluster-determining threshold. Localized activations were then identified using family-wise error correction. Activations to the reinforced stimulus (CS+) that increased from the beginning to end (“across”) threat conditioning were more extensive within the GAD+ID group. Increased activations to the CS+ across extinction learning were greater within the GAD-ID than the GAD+ID group, and delayed 24h in the latter. Greater sleep efficiency was associated with decreased activations across threat conditioning, but with increased activations across extinction learning. Better sleep quality promoted greater engagement of neural substrates of extinction learning. The GAD+ID group failed to engage brain areas supporting extinction learning immediately following threat conditioning, but did so when stimuli were again presented following a delay.

**Highlights:** Generalized Anxiety Disorder subjects with moderate/severe vs absent/mild insomnia compared

Neural responses to threat conditioning, extinction learning/memory analyzed and sleep recorded

Low and high thresholds identified large-scale networks and localized activations respectively

Threat and regulatory activations at extinction learning in mild insomnia delayed when more severe

Better sleep quality predicted greater activation of regulatory areas during extinction learning

## 1. Introduction

Pathological anxiety constitutes a major public health concern in the US, affecting approximately 18% of adults (Kessler et al., 2005; Kessler et al., 2010). Among such disorders, generalized anxiety disorder (GAD) is the most common and is a risk factor for psychiatric comorbidities, especially depression and substance abuse (Stein and Sareen, 2015). Anxiety-related disorders and insomnia show immense comorbidity with up to 30% of those with insomnia also having anxiety disorders, and with GAD being the most frequent comorbidity (Glidewell et al., 2015; Ohayon and Roth, 2003). Whereas, it is well documented that sleep disturbance is ubiquitous in persons with existing depressive and anxiety-related disorders (Alvaro et al., 2013; APA, 2013b; Johnson et al., 2006; Krystal et al., 2017; Mellman, 2008; Ohayon and Roth, 2003; Riemann, 2007), there is now abundant evidence that insomnia disorder (ID) also increases the odds of incident depressive and anxiety disorders (Baglioni et al., 2011a; Baglioni et al., 2016; Baglioni and Riemann, 2012; Baglioni et al., 2011b; Breslau et al., 1996; Ford and Cooper-Patrick, 2001; Ford and Kamerow, 1989; Hertenstein et al., 2018; Jansson-Frojmark and Lindblom, 2008; Neckelmann et al., 2007). For example, a recent meta-analysis showed that individuals with preexisting ID are 3.2X more likely to develop an anxiety disorder and 2.8X more likely to develop major depression (Hertenstein et al., 2018). We have suggested that preexisting sleep disturbance can disrupt emotion regulatory circuits and that mutually reinforcing sleep disruption and emotion-regulatory impairment can increase the probability that pathological anxiety will arise (Pace-Schott et al., 2015; Pace-Schott et al., 2023; Pace-Schott et al., 2017).

### 1.1 Sleep symptoms of GAD

GAD is the anxiety disorder most commonly comorbid with ID (Breslau et al., 1996; Monti and Monti, 2000; Ohayon et al., 1998; Tsypes et al., 2013). More than half of patients with GAD report insomnia symptoms (Belanger et al., 2004), and difficulty initiating and/or maintaining sleep is a DSM-5 diagnostic criterion for GAD (APA, 2013a). (Monti and Monti, 2000) reviewed polysomnographic (PSG) studies comparing GAD and healthy controls (HC) and reported, in GAD, longer sleep onset latency (SOL) and/or more wake time after sleep onset (WASO), reduced total sleep time (TST) and/or sleep efficiency (SE), as well as increased Stage 1 NREM and/or reduced slow wave (SWS) and rapid eye movement (REM) sleep. Similarly, a systematic review by (Cox and Olatunji, 2016) reported that, compared to healthy controls, different studies have shown that individuals with GAD display decreased TST, increased SOL and WASO as well as various differences in sleep stage percentages suggestive of abnormal sleep architecture.

### 1.2 Functional differences between GAD and healthy controls

Both white matter and grey matter structural differences between individuals with GAD and healthy controls (HC) have been reported (Ma et al., 2019; Ou et al., 2024; Yang et al., 2020). Similarly, differences in resting state functional connectivity (rsFC) between GAD and HC have shown specific alterations of limbic circuitry involved in the modulation of anxiety and fear (Li et al., 2020; Ma et al., 2019; Oathes et al., 2015; Peterson et al., 2014; Qiao et al., 2017), alterations recently also seen using high-density EEG (Wang et al., 2025). For example, prefrontal cortex (PFC) rsFC was reduced to areas of the salience network (SN), including the anterior insula (AIC) and dorsal anterior cingulate (dACC) (Etkin et al., 2009). Other studies link GAD with reduced rsFC between the amygdala and frontal regions such as the dorsolateral PFC (DLPFC), suggesting reduced top-down control of negative emotion (Liu et al., 2015; Roy et al., 2013; Wang et al., 2021). Differences between GAD and HC have also been observed in regions associated with the default and salience networks (Li et al., 2023b; Wang et al., 2016; Xiong et al., 2020).

In contrast, in functional imaging studies using task-based emotional learning and reactivity protocols, individuals with GAD versus HC have shown diverse and often inconsistent results even among those studies using similar protocols (Fonzo and Etkin, 2017; Hilbert et al., 2014). Fonzo and Etkin (2017) suggest that this variability is due to the domain-variable nature of each individual’s unique, private inner thoughts that generate anxiety in this disorder. Nonetheless, some general findings have emerged. A recent review found that PFC hypoactivation with abnormal PFC-amygdala connectivity was the most typical finding in task-based fMRI studies of GAD (Mochcovitch et al., 2014). A more recent meta-analysis (Kolesar et al., 2019) also identified functional differences between GAD and HC in DLPFC, anterior cingulate cortex (ACC), amygdala, and hippocampus suggesting disturbance of emotion processing in GAD. Kolesar et al. (2019) suggest that seeking task-based differences at the level of large-scale networks identified using rsFC may be an avenue to characterize differences between GAD and HC while casting a wide enough net to accommodate a high degree of inter-individual variation in the neural substrates of GAD symptoms in responses to task-based paradigms.

### 1.3 Functional differences between ID and healthy controls

Individuals with Insomnia Disorder (ID) also show rsFC abnormalities relative to HC similar to those reported with GAD. For example, in an rsFC study comparing ID with HC (Huang et al., 2012), the left amygdala showed reduced rsFC with the left insula, bilateral striatum, and thalamus but increased connectivity with premotor (PMC) and sensorimotor cortex. The authors suggested that decreased amygdala connectivity was evidence for impaired emotional processing in ID and that increased cortical connectivity was compensatory. Another study showed hyper-connectivity in ID between the insula and rostral ACC (rACC) suggesting increased connectivity within the salience network (SN) (Wang et al., 2017), a finding consistent with the hypothesized hyperarousal underlying ID (Bonnet and Arand, 2010; Kay and Buysse, 2017; Riemann et al., 2010). In healthy individuals, sleep deprivation has also been associated with decreased amygdala-PFC functional connectivity both in rsFC paradigms (Lei et al., 2015; Motomura et al., 2017; Shao et al., 2014) and in response to emotional images (Motomura et al., 2013; Yoo et al., 2007). A comparison of individuals with GAD, ID, and HC using rsFC with seeds in the fear/salience network (bilateral amygdala, dACC) and the extinction/default network (vmPFC), showed connectivity between the left amygdala and bilateral rostral ACC (rACC) in ID that was intermediate between their connectivity in HC and GAD (Pace-Schott et al., 2017). The rACC is an emotion regulatory region (Dodhia et al., 2014; Pizzagalli, 2011) thus suggesting that weakened emotional regulatory circuits may predispose those with ID to develop GAD.

### 1.4 Threat conditioning and extinction in GAD

Although numerous emotional-task based neuroimaging studies of GAD have been published (for review see Fonzo and Etkin, 2017; Hilbert et al., 2014; Kolesar et al., 2019; Mochcovitch et al., 2014), only two of these studies included threat conditioning (Cha et al., 2014; Cooper et al., 2018). Cha et al. (2014) reported that overgeneralization of fear in GAD versus HC was accompanied by a less accurate correspondence between activations in the mesocorticolimbic circuit and the degree to which generalization stimuli resembled the originally reinforced stimulus, suggesting abnormal threat processing in GAD. To the best of our knowledge, the current study is the first to examine the sleep correlates of threat conditioning and extinction in GAD.

### 1.5 Threat conditioning and extinction in ID

Research on the neural bases and sleep correlates of threat conditioning and extinction in ID is limited, with one study showing that, compared to good sleepers, recruitment of extinction-learning related structures is delayed in those with ID (Seo et al., 2018), and another study reporting that REM is associated with poorer extinction memory in ID but not in HC (Bottary et al., 2020). In addition, a theoretical paper (Perogamvros et al., 2020), a protocol proposal (Sun et al., 2024) and a comorbid ID and PTSD treatment study (Hunt et al., 2023) also address extinction in ID. However, to the best of our knowledge, no studies have examined the influence of ID on the neural bases of extinction learning and memory in GAD.

### 1.6 Hypotheses

In the current study, we used a validated 2-day, 3-phase experimental protocol (Milad et al., 2013; Milad et al., 2007; Seo et al., 2018) to compare neural activations during threat conditioning, extinction learning and extinction recall between individuals with GAD and comorbid moderate to severe insomnia (GAD+ID) and those without or with sub-threshold insomnia (GAD-ID). In addition, among both groups combined, we examined associations of these activations with pre-selected sleep variables derived from PSG, actigraphy, sleep diaries and retrospective questionnaires. We hypothesized that, across all 3 phases of this experiment: (1) GAD+ID compared to GAD-ID would exhibit greater activation of threat-related neural structures (i.e., those of the SN) and lesser activation of regulatory regions (e.g., those of the fronto-parietal control network, FPCN). (2) Activations of regulatory regions would positively correlate with sleep quality (subjective and objective SE), REM percent and SWS percent, whereas these same sleep variables would negatively correlate with activations of SN regions. (3) As suggested by literature on GAD and ID wakefulness (Dodds et al., 2017; Wang et al., 2023), lower parasympathetic tone in REM and SWS would predict greater activation of threat-related SN, and diminished activation of regulatory regions. We have previously associated parasympathetic measures negatively with hyperarousal (Daffre et al., 2022), and positively with extinction memory (Yuksel et al., 2024a) as well as positively and negatively with neural activations using the current protocol (Seo et al., 2022).

### 1.7 Localized and network neural activations to threat conditioning and extinction

Based on animal studies and initial human neuroimaging, two opposed neural networks have been hypothesized to support the acquisition of conditioned fear and the extinction of such fear (Graham and Milad, 2011; Milad and Quirk, 2012). The amygdala and dACC activate during acquisition and expression of conditioned threat whereas the ventromedial prefrontal cortex (vmPFC) and hippocampus activate during learning and recall of extinction (Milad et al., 2007). However, recent meta-analyses have shown that, in humans, much broader regions of the forebrain and brainstem are recruited in experimental models of both threat conditioning and extinction (Fullana et al., 2018; Fullana et al., 2016; Wen et al., 2024).

In fMRI analyses, the use of highly conservative corrections, such as nonparametric permutation, to restrict family-wise error (FWE) to p<0.05 has been strongly advocated following an influential study by Eklund et al. (2016) showing that the probability of false positives had been greatly underestimated in prior fMRI studies. However, by thus maximizing specificity and minimizing Type 1 error, sensitivity and statistical power are lost and the probability of Type II error is greatly increased (Noble et al., 2020).

Interpretations of neuroimaging findings have increasingly invoked the large-scale networks initially revealed in rsFC studies but clearly relevant to task-based analyses (Bressler and Menon, 2010; Kong et al., 2024; Pessoa and McMenamin, 2017; Uddin, 2023; Uddin et al., 2023; Uddin et al., 2019). Moreover, Multivariate Pattern Analysis (MVPA) has revealed a high degree of overlap and variability in the activation of such networks in response to specific behavioral tasks (Pessoa, 2018; Wen et al., 2024; Westlin et al., 2023). Such variability presents a plausible explanation for the lack of replicability among specific regional activations to seemingly equivalent behavioral tasks (Westlin et al., 2023). In a recent review, Westlin et al. (2023) argue that instead of specific regions or groups of regions consistently becoming activated by specific neurobehavioral functions, and thus that the varying results seen in the literature must reflect task differences, a particular function can elicit multiple distributed and varying activities depending upon contextual factors many of which cannot be controlled.

### 1.8 Network and localized neural activations in the current study

Given the alternative, but non-exclusive perspectives described above (1.6, 1.7), we have pursued a two-tiered approach whereby activations to specific fMRI contrasts in a threat-conditioning and extinction protocol are first reported and interpreted in relation to large-scale networks (based on (Uddin et al., 2019; Yeo et al., 2011) using a liberal detection threshold, after which regions surviving stringent FWE correction are reported to provide additional specificity. These activations will first be compared between GAD+ID and GAD-ID. Correlations of these activations with the sleep variables described above (1.4) will then be examined in the entire sample. Among networks, we will specifically focus on the SN, FPCN and default (DMN) networks, which have shown rsFC differences between GAD and HC as well as correlations with anxiety symptoms (Li et al., 2023a; Li et al., 2023b; Li et al., 2016; Yuan et al., 2023).

## 2. Methods

### 2.1 Participants

Participants with GAD symptoms were recruited from the greater Boston general public via social media and electronic bulletin boards. Interested individuals first completed a telephone screening that included the Generalized Anxiety Disorder 7-Item (GAD-7) Questionnaire (Spitzer et al., 2006). Those scoring >10 on the GAD-7 who self-reported GAD symptoms and did not report exclusion criteria were then scheduled for an interview. The psychiatric interview confirmed GAD diagnosis and absence of exclusion criteria using the Structured Clinical Interview for DSM-5, Research Version (SCID-5 RV) (First et al., 2015). The Pittsburgh Structured Clinical Interview for Sleep Disorders (SCIDSLD), a widely used (Insana et al., 2013; Stocker et al., 2017) but unpublished sleep interview, was used to diagnose DSM-5 ID, identify sleep-related exclusion criteria (*vis*., an intrinsic sleep disorder other than ID), and obtain a comprehensive picture of sleep habits. For detailed exclusion criteria, please see Supplemental Materials. Inclusion criteria included diagnoses of GAD using the SCID-5 RV and ID using the SCIDSLD, a score of ≥ 10 on the Generalized Anxiety Disorder 7-Item (GAD-7) Questionnaire (Spitzer et al., 2006) and a score <u>≥</u> 62 on the Penn State Worry Questionnaire (PSWQ) (Behar et al., 2003). If all criteria were met, the participant was assigned to one of two groups: GAD with (GAD+ID) or without (GAD-ID) Insomnia Disorder. Among the 35 individuals who met GAD criteria, GAD+ID was defined as those individuals with an Insomnia Severity Index (ISI) (Bastien et al., 2001) ≥ 13 (N 21, mean 17.8, SD 3.6, range 13-25) and GAD-ID as those with an ISI score ≤ 12 (N 14, mean 6.4, SD 3.4, range 1-12) (please see Supplemental Material for details). The age 40 upper limit was selected to ensure that subjects retained a measurable amount of SWS, a sleep stage that is closely linked with sleep-dependent memory consolidation (Diekelmann and Born, 2010; Rasch and Born, 2013), and which has declined considerably by middle age (Ohayon et al., 2004). Of the 35 participants meeting criteria, 5 were excluded due to excessive movement during fMRI scans, poor quality ambulatory sleep (actigraphy and/or PSG) data, or both. Of the remaining 30, 28 (93%) were female and both males were in GAD-ID. All study procedures were approved by the Partners Healthcare Institutional Review Board. Participants provided written informed consent and were paid for their participation.

### 2.2 Procedures

Interviews were followed by a 2-week sleep assessment period using sleep diaries and actigraphy to estimate subjective and objective habitual sleep parameters. During this period, an acclimation/diagnostic night with ambulatory polysomnography (PSG) was completed in order to acclimate the participant to PSG and detect obstructive sleep apnea (OSA) or periodic limb movement disorder (PLMD) of a severity sufficient to warrant referral for treatment. No participants were excluded based on this criterion following review of the acclimation night PSG by a highly experienced clinical Registered Polysomnographic Technician (RSPGT). Ambulatory PSG used the Compumedics Somte-PSG system and a standard montage (Bottary et al., 2020; Seo et al., 2018). The second PSG occurred on the night before the first evening MRI scan (Baseline night) and a third recording occurred during the night following Threat Conditioning and Extinction Learning, the night during which initial consolidation of threat and extinction learning was expected to take place (Consolidation night). Participants were instrumented for PSG in the lab and then returned to their homes to sleep in their usual environment. Please see Supplemental Material regarding the rationale for holding experimental sessions in the evening. Please also see Figure S1 in Supplemental Material for a graphic representation of the protocol.

On the evening following the Baseline PSG, participants began a validated 2-day threat-conditioning and extinction protocol (Milad et al., 2013; Milad et al., 2007; Seo et al., 2018) carried out, with simultaneous measurement of skin conductance level (SCL), in a 3-T Siemens Prisma scanner. Threat Conditioning and Extinction Learning phases occurred on the first evening followed 24 hours later by the Extinction Recall phase. During Threat Conditioning, partial reinforcement (62.5%) with a mild electric shock produced conditioned skin conductance responses (SCR) to 2 differently colored lamps (CS+), but not a third color (CS-). During Extinction Learning, one CS+ (CS+E) but not the other (CS+U) was extinguished by un-reinforced presentations. At Extinction Recall, all 3 stimulus types were presented. (Please see Milad et al., 2013; Milad et al., 2007; Seo et al., 2018 and Supplemental Materials for additional details.) Blood-oxygen-level-dependent (BOLD) functional magnetic resonance imaging (fMRI) signal was recorded throughout all phases of the protocol. Concurrently, skin conductance level (SCL) was recorded throughout the protocol and shock expectancy ratings were made by participants at the end of each phase.

### 2.3 Sleep (diary, actigraphy, PSG) and clinical measures (Table 2)

#### 2.3.1 Sleep diary: Evening-Morning Sleep Questionnaire (EMSQ)(Pace-Schott et al., 1994)

The evening portion of the EMSQ sleep diary queried the time at which the participant began to attempt sleep. Morning portions queried the time of waking for the day, subjective sleep onset latency (SOL), and number and duration of nocturnal awakenings (WASO). Subjective sleep efficiency (SE) was computed from these data as total sleep time (TST) as a proportion of time in bed (TIB) ([TIB – (SOL + WASO]/TIB).

#### 2.3.2 Actigraphy

The Actiwatch 2 (Philips Respironics, Bend, OR) was a motion-sensitive wrist monitor that counted arm movements in 1-min epochs. Participants pressed an event marker when beginning to attempt sleep and when waking for the day. Within this period, epochs were scored as sleep or wake using the Actiwatch default algorithm. Objective SE was computed from these data. (Please see Supplemental Materials for additional details.)

#### 2.3.3 Ambulatory PSG

The Somte-PSG ambulatory recorder (Compumedics USA, Inc., Charlotte, NC) was worn on the chest in a cloth pack. The montage included the following channels: 6 EEG (F3, F4, C3, C4, O1, O2) referenced to contralateral mastoids (A1, A2), 2 EOG, 2 submental EMG and 2 ECG (right clavicle, left 5th intercostal space). A research-experienced RPSGT, blind to participant group, scored sleep following standard AASM criteria (Iber and al., 2007) using Compumedics Profusion 4.0 software. Heart-rate variability (HRV) was measured using Kubios HRV Premium software (Kubios Oy, Kuopio, Finland). Measures of parasympathetic activity in all REM and SWS periods exceeding 5 min were computed in the time domain as RMSSD (root-mean square differences of successive R-R intervals). PSG data were used to compute sleep-stage percentages and HRV (Please see Supplemental Materials for additional details.)

#### 2.3.4 *Retrospective sleep and clinical questionnaires* are listed in Table 2 and described in Supplemental Materials

### 2.4 fMRI methods

Whole brain images were collected using a 32-channel head coil inside a 3 T Siemens Prisma MAGNETOM scanner (Siemens Medical Systems, Iselin, NJ). Detailed fMRI data acquisition and preprocessing methods are reported in (Seo et al., 2018) and (Seo et al., 2022) and summarized in Supplemental Methods.

#### 2.4.1 First-level analyses

Nine contrasts (3 per phase) were analyzed to generate whole-brain first-level parametric maps (Table 1). For Threat Conditioning, differential BOLD activations to the CS+ versus the CS- for the second to fourth (early CS+>CS-) and the last 4 (late CS+>CS-) as well as across the Threat-Conditioning phase (early CS+>late CS+) were averaged for each of the 2 CS+’s. (During Threat Conditioning, the first presentation of each of the two CS+s, along with their ordinally corresponding CS-, were excluded from analyses because pairing with the US, and hence threat learning, had not yet occurred.) For Extinction Learning, activations to the CS+E versus the CS- were analyzed for the first 4 (early CS+E>CS-) and last 4 (late CS+E>CS-) trials of Extinction Learning as well as across the Extinction Learning phase (late CS+E> early CS+E). For Extinction Recall, only the first 4 trials of each stimulus type were modeled to capture extinction recall but not the new extinction learning occurring during this phase. The early CS+E>early CS+U contrast identified regions that were more responsive to the CS+ that was both threat-conditioned and subsequently extinguished (CS+E) compared with the CS that was previously only threat-conditioned (CS+U). The first 4 (early) CS+E>CS- and first 4 (early) CS+U>CS- contrasts were also modeled. (see Table 1 for working interpretations of contrasts). Standardized methods for recording, preprocessing, motion correction, and first-level analysis of all task-based fMRI procedures for the threat conditioning and extinction paradigm are detailed in references (Seo et al., 2018) and (Seo et al., 2022) and summarized in Supplemental Methods.

**Table 1.** Interpretation of fMRI contrasts in threat conditioning and extinction protocol

| **Phase and contrast** | **Neurocognitive construct** |
| --- | --- |
| **Fear Conditioning** | . |
| Early CS+ > Early CS- | Differential threat perception at the beginning of Threat Conditioning |
| Late CS+ > Late CS- | Differential threat perception at the end of Threat Conditioning |
| Late CS+ > Early CS+ | Acquisition of threat perception across Threat Conditioning |
| **Extinction Learning** |  |
| Early CS+E > Early CS- | Differential threat perception at the beginning of Extinction Learning |
| Late CS+E > Late CS- | Differential threat perception at the end of Extinction Learning |
| Late CS+E > Early CS+E | Acquisition of extinction, decrease in threat perception over phase |
| **Extinction Recall** |  |
| Early CS+E > Early CS+U | Distinguishing extinguished from unextinguished threat perception |
| Early CS+E > Early CS- | Remaining differential threat perception followed by its extinction 24-h earlier |
| Early CS+U > Early CS- | Differential threat perception remaining following 24-h delay |

#### 2.4.2 Second-level analyses

We compared brain activation to the 9 contrasts within and between GAD+ID and GAD- ID groups as well as associations of these activated areas with sleep parameters within the entire sample. Within- and between-group analyses and multiple regression analyses with sleep parameters were run using age and sex as covariates to account for potential confounds. Activations within each group and in the entire sample were defined using both low- and high-cluster determining thresholds (CDT) (defined in Statistical Analyses). Both high- and low-threshold activations for the 9 contrasts are shown in Table 3. Low-threshold activations were modeled to illustrate the potential involvement of one or more of 6 large-scale networks following a modified Yeo et al. 7-network template (Uddin et al., 2019; Yeo et al., 2011). The current modification of the Yeo et al. template equates the ventral attention network with the salience network and assigns Yeo et al.’s cortical limbic network areas to the default network (see Kong et al., 2024; Uddin et al., 2019). Thus, low-threshold activations were assessed in salience (SN), default (DMN), frontoparietal control (FPCN), somatomotor (SM), dorsal attention (DAN), and visual (VN) networks. Among sub-cortical activation clusters, amygdala and hippocampal activations are assigned to salience and default networks respectively, while remaining agnostic regarding network involvement of basal ganglia and thalamic clusters (reported here as “subcortical”). Here, we first report clusters activated at a low-threshold by listing anatomic regions (see glossary) in which clusters are seen in a specified network, followed by the total number of voxels in all anatomic regions within that network [e.g., “default network (DMPFC, PCC, hippocampus; 408 DMN)”] as shown also in Fig. 1. Low-threshold results are followed by any high-threshold clusters that survived SPM12 FWE correction. Low-threshold voxel counts are inclusive of both hemispheres (Fig. 1). However, their laterality is provided in Tables 3, 4 and Fig. 3, 4. High-threshold laterality is indicated in both text and tables. For the sake of clarity, only changes in activation to the CS+ across Threat Conditioning and Extinction Learning phases (i.e., early CS+ vs. late CS+ contrasts) are compared between groups and regressed against sleep parameters in the main paper. However, for these two phases, low- and high-threshold results for CS+>CS- contrasts at the beginning (early) and end (late) of the Threat Conditioning and Extinction Learning phases are provided for group comparisons in Table 3 and for multiple regressions against sleep parameters in Table 4 as well as in Supplemental Results.

**Figure 1.**
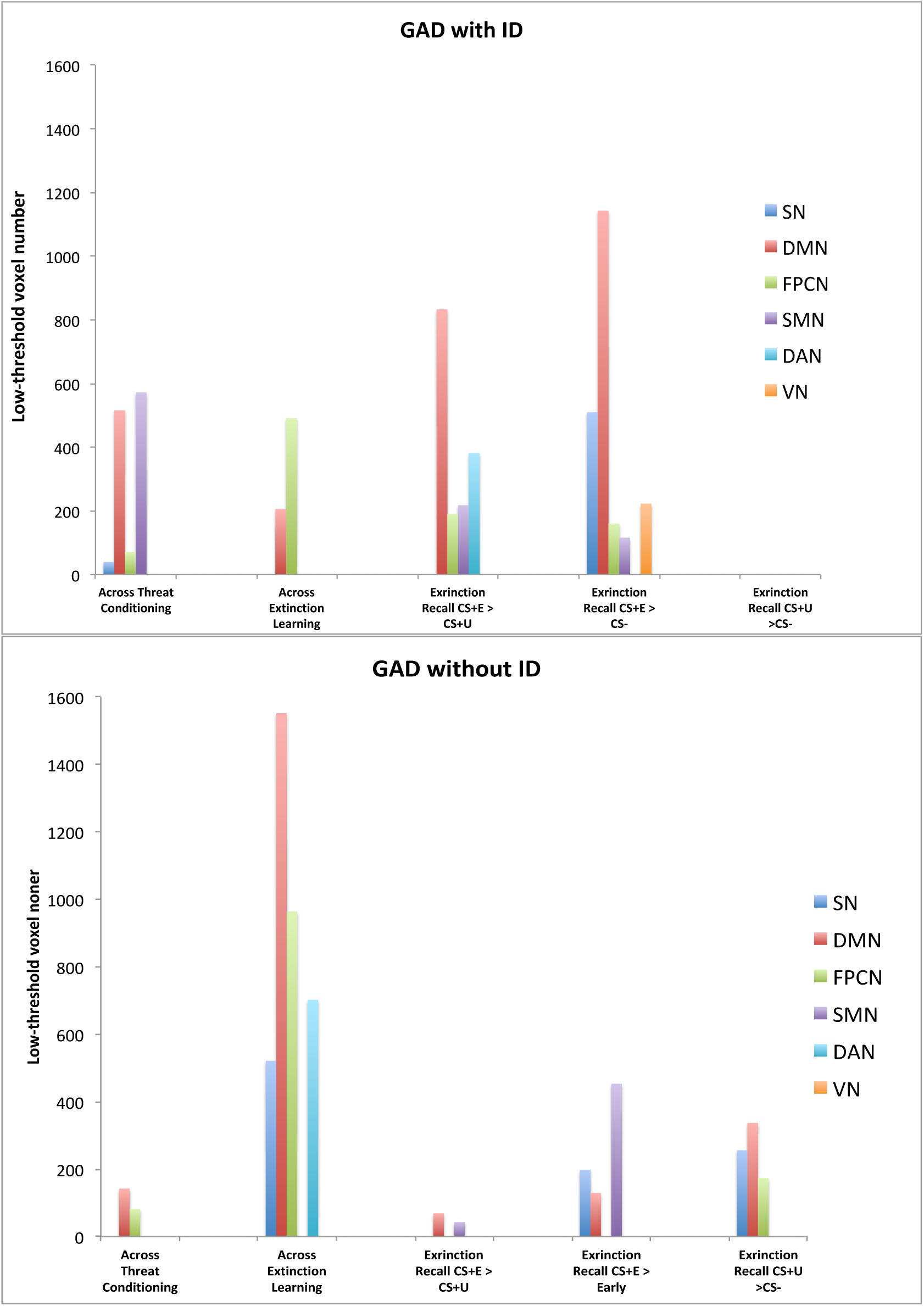
Low-threshold activations within large scale brain networks (Yeo et al., 2011) across Threat Conditioning and Extinction Learning and at early Extinction Recall in individuals with GAD. Low threshold is defined as a cluster-determining threshold (CDT) of *p* <0.005 and clusters with at least 10 contiguous voxels. GAD with ID (GAD+ID) was defined as having an Insomnia Severity Index (ISI) of 13 and above (mean ISI 17.76, range 13-25) and GAD without ID (GAD-ID) is defined as having an ISI of 12 or below (mean ISI 6.43, range 1-12). See Table 3 and Figure 2 for clusters activated at high threshold that used a CDT of *p* <0.001 and survived SPM12 family-wise error (FWE) correction at *p* <0.05.

**Figure 2.**
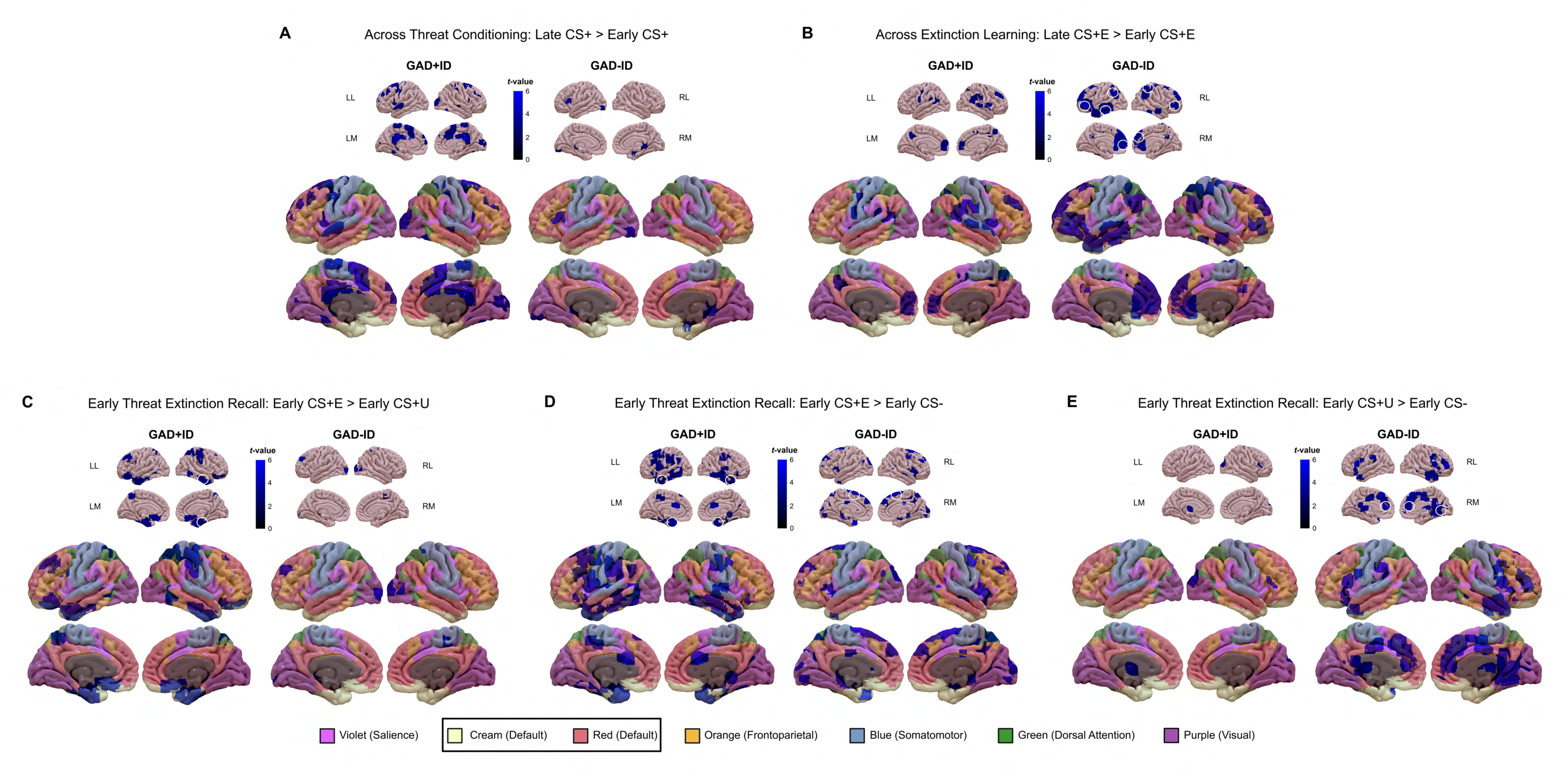
Within-group activations to contrasts across the Threat Conditioning (panel A), across the Extinction Learning phase (B), and during early Threat Extinction Recall phase (C, D, and E). In each of panels A-E, *low-threshold activations* (cluster-determining threshold [CDT] of *p* <0.005 and at least 10 contiguous) are indicated (in graded blues) on both the upper sets of 8 smaller images and on the lower sets of 8 larger images which also display the 7 large-scale networks reported by Yeo et al. (Uddin et al., 2019; Yeo et al., 2011) color coded as indicated by the bottom legend. (Note that the Yeo ventral attention network is renamed “salience” and Yeo limbic network is added to the default network). Clusters circled in white on the upper (smaller) images indicate *high-threshold activations* that survived family-wise error correction at *p* <0.05 with a cluster-determining threshold of *p* <0.001. Abbreviations: CS+, conditioned stimulus reinforced by the unconditioned (shock) stimulus; CS-, nonreinforced conditioned stimulus; CS+E, CS+ extinguished during Extinction Learning; CS+U, un-extinguished CS+; GAD, generalized anxiety disorder; ID, insomnia disorder; LL, left lateral view; RL, right lateral view; LM, left medial view; RM, right medial view.

**Figure 3.**
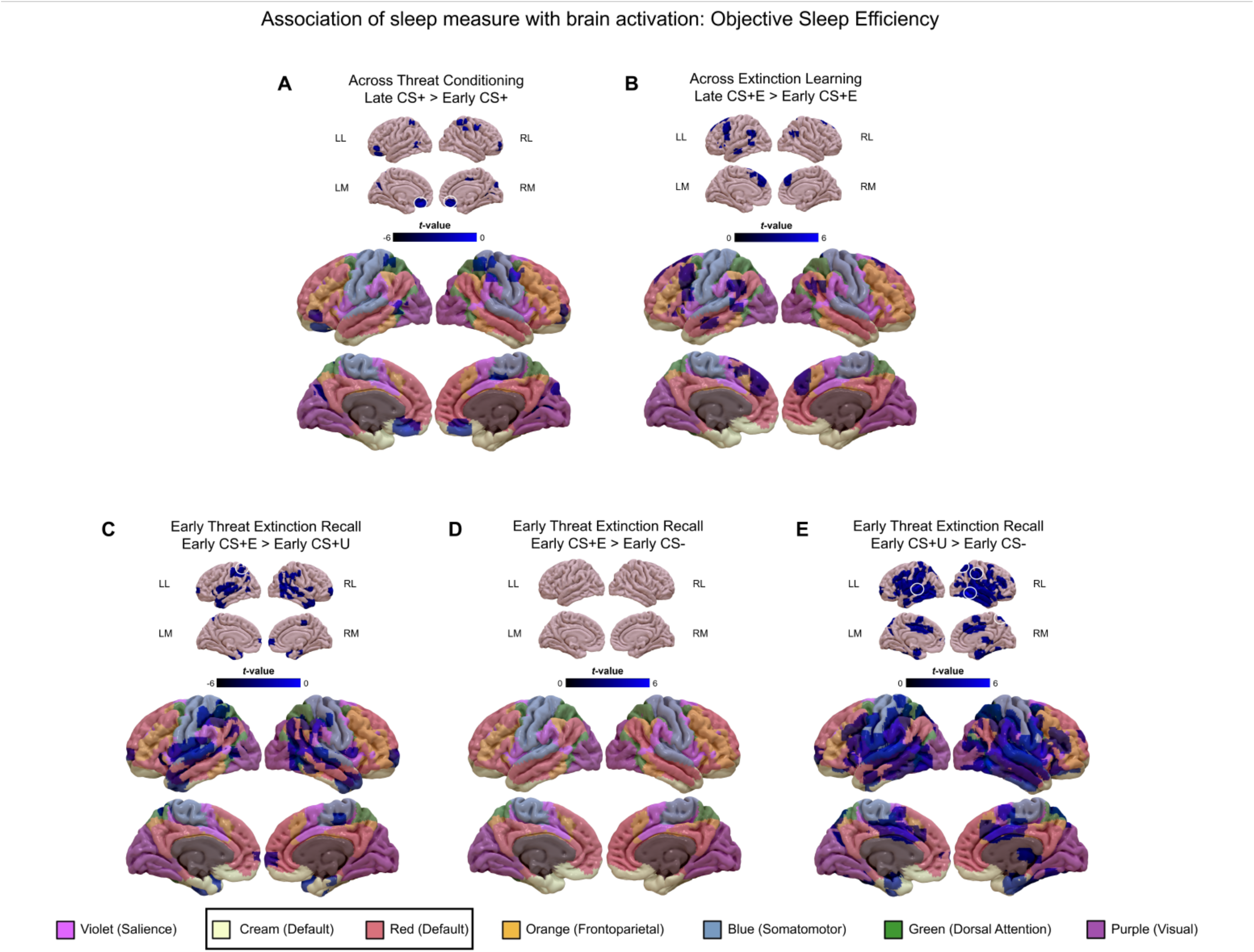
Associations of activations in the whole-sample (GAD+ID and GAD-ID) with **objective sleep efficiency** across Threat Conditioning (panel A), across Extinction Learning (panel B), and at early Threat Extinction Recall (panels C, D, and E). In each of panels A-E, *low-threshold activations* (cluster-determining threshold [CDT] of *p* <0.005 and at least 10 contiguous) are indicated in graded blues on both the upper sets of 8 smaller images and on the lower sets of 8 larger images which also display the 7 large-scale networks reported by Yeo et al. (Uddin et al., 2019; Yeo et al., 2011) color coded as indicated by the bottom legend. Clusters circled in white on the upper (smaller) images indicate *high-threshold activations* that survived family-wise error correction at *p* <0.05 with a CDT of *p* <0.001. Abbreviations: CS+, conditioned stimulus reinforced by the unconditioned (shock) stimulus; CS-, nonreinforced conditioned stimulus; CS+E, CS+ extinguished during Extinction Learning; CS+U, un-extinguished CS+; GAD, generalized anxiety disorder; ID, insomnia disorder; LL, left lateral view; RL, right lateral view; LM, left medial view; RM, right medial view.

**Table 2.**
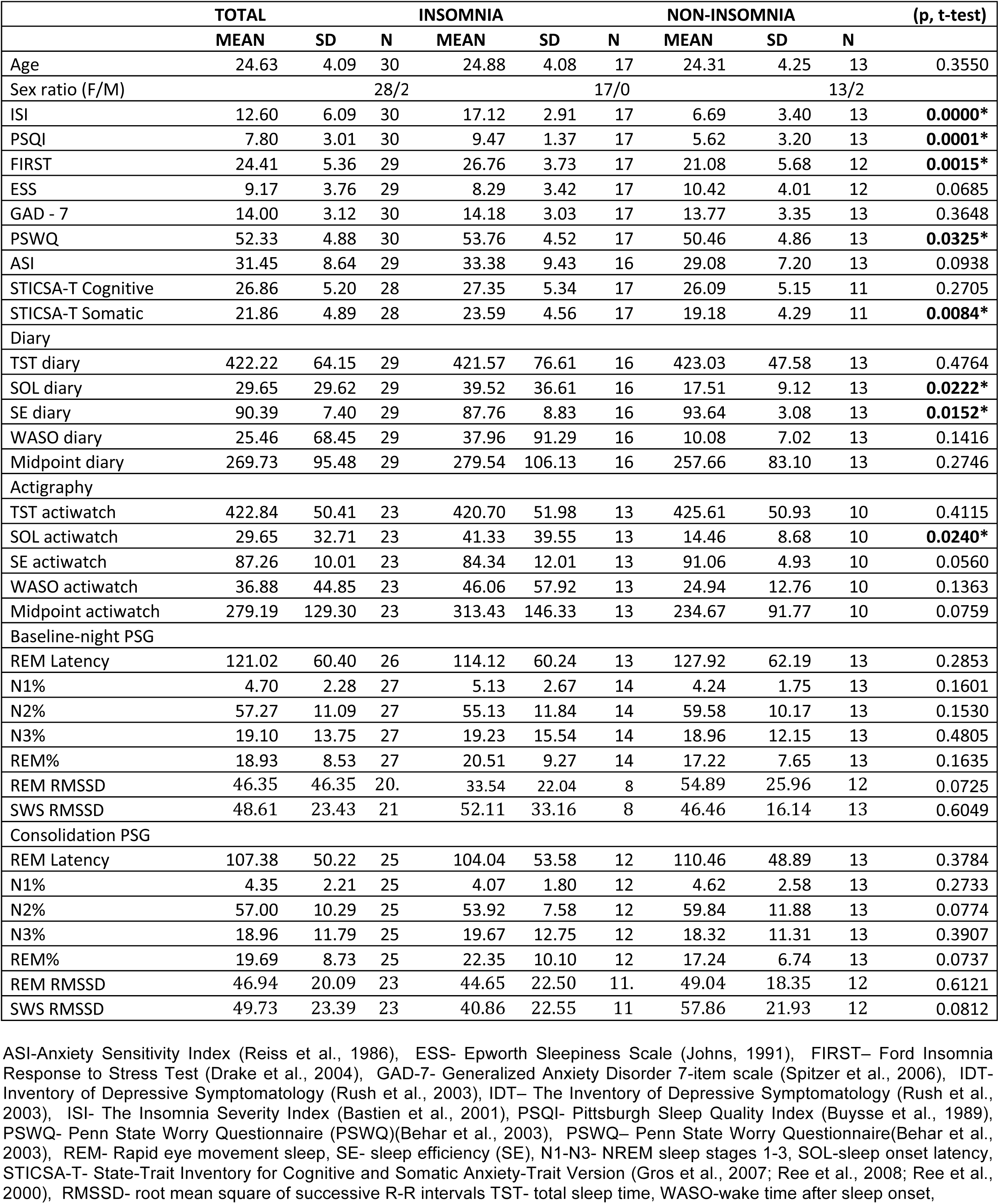
Demographic, psychometric and sleep parameters of individuals with GAD from whom fMRI data was available.

|  | TOTAL |  |  | INSOMNIA |  |  | NON-INSOMNIA |  |  | (p, t-test) |
| --- | --- | --- | --- | --- | --- | --- | --- | --- | --- | --- |
|  | MEAN | SD | N | MEAN | SD | N | MEAN | SD | N |  |
| Age | 24.63 | 4.09 | 30 | 24.88 | 4.08 | 17 | 24.31 | 4.25 | 13 | 0.3550 |
| Sex ratio (F/M) |  |  | 28/2 |  |  | 17/0 |  |  | 13/2 |  |
| ISI | 12.60 | 6.09 | 30 | 17.12 | 2.91 | 17 | 6.69 | 3.40 | 13 | **0.0000\*** |
| PSQI | 7.80 | 3.01 | 30 | 9.47 | 1.37 | 17 | 5.62 | 3.20 | 13 | **0.0001\*** |
| FIRST | 24.41 | 5.36 | 29 | 26.76 | 3.73 | 17 | 21.08 | 5.68 | 12 | **0.0015\*** |
| ESS | 9.17 | 3.76 | 29 | 8.29 | 3.42 | 17 | 10.42 | 4.01 | 12 | 0.0685 |
| GAD - 7 | 14.00 | 3.12 | 30 | 14.18 | 3.03 | 17 | 13.77 | 3.35 | 13 | 0.3648 |
| PSWQ | 52.33 | 4.88 | 30 | 53.76 | 4.52 | 17 | 50.46 | 4.86 | 13 | **0.0325\*** |
| ASI | 31.45 | 8.64 | 29 | 33.38 | 9.43 | 16 | 29.08 | 7.20 | 13 | 0.0938 |
| STICSA-T Cognitive | 26.86 | 5.20 | 28 | 27.35 | 5.34 | 17 | 26.09 | 5.15 | 11 | 0.2705 |
| STICSA-T Somatic | 21.86 | 4.89 | 28 | 23.59 | 4.56 | 17 | 19.18 | 4.29 | 11 | **0.0084\*** |
| Diary |  |  |  |  |  |  |  |  |  |  |
| TST diary | 422.22 | 64.15 | 29 | 421.57 | 76.61 | 16 | 423.03 | 47.58 | 13 | 0.4764 |
| SOL diary | 29.65 | 29.62 | 29 | 39.52 | 36.61 | 16 | 17.51 | 9.12 | 13 | **0.0222\*** |
| SE diary | 90.39 | 7.40 | 29 | 87.76 | 8.83 | 16 | 93.64 | 3.08 | 13 | **0.0152\*** |
| WASO diary | 25.46 | 68.45 | 29 | 37.96 | 91.29 | 16 | 10.08 | 7.02 | 13 | 0.1416 |
| Midpoint diary | 269.73 | 95.48 | 29 | 279.54 | 106.13 | 16 | 257.66 | 83.10 | 13 | 0.2746 |
| Actigraphy |  |  |  |  |  |  |  |  |  |  |
| TST actiwatch | 422.84 | 50.41 | 23 | 420.70 | 51.98 | 13 | 425.61 | 50.93 | 10 | 0.4115 |
| SOL actiwatch | 29.65 | 32.71 | 23 | 41.33 | 39.55 | 13 | 14.46 | 8.68 | 10 | **0.0240\*** |
| SE actiwatch | 87.26 | 10.01 | 23 | 84.34 | 12.01 | 13 | 91.06 | 4.93 | 10 | 0.0560 |
| WASO actiwatch | 36.88 | 44.85 | 23 | 46.06 | 57.92 | 13 | 24.94 | 12.76 | 10 | 0.1363 |
| Midpoint actiwatch | 279.19 | 129.30 | 23 | 313.43 | 146.33 | 13 | 234.67 | 91.77 | 10 | 0.0759 |
| Baseline-night PSG |  |  |  |  |  |  |  |  |  |  |
| REM Latency | 121.02 | 60.40 | 26 | 114.12 | 60.24 | 13 | 127.92 | 62.19 | 13 | 0.2853 |
| N1% | 4.70 | 2.28 | 27 | 5.13 | 2.67 | 14 | 4.24 | 1.75 | 13 | 0.1601 |
| N2% | 57.27 | 11.09 | 27 | 55.13 | 11.84 | 14 | 59.58 | 10.17 | 13 | 0.1530 |
| N3% | 19.10 | 13.75 | 27 | 19.23 | 15.54 | 14 | 18.96 | 12.15 | 13 | 0.4805 |
| REM% | 18.93 | 8.53 | 27 | 20.51 | 9.27 | 14 | 17.22 | 7.65 | 13 | 0.1635 |
| REM RMSSD | 46.35 | 46.35 | 20. | 33.54 | 22.04 | 8 | 54.89 | 25.96 | 12 | 0.0725 |
| SWS RMSSD | 48.61 | 23.43 | 21 | 52.11 | 33.16 | 8 | 46.46 | 16.14 | 13 | 0.6049 |
| Consolidation PSG |  |  |  |  |  |  |  |  |  |  |
| REM Latency | 107.38 | 50.22 | 25 | 104.04 | 53.58 | 12 | 110.46 | 48.89 | 13 | 0.3784 |
| N1% | 4.35 | 2.21 | 25 | 4.07 | 1.80 | 12 | 4.62 | 2.58 | 13 | 0.2733 |
| N2% | 57.00 | 10.29 | 25 | 53.92 | 7.58 | 12 | 59.84 | 11.88 | 13 | 0.0774 |
| N3% | 18.96 | 11.79 | 25 | 19.67 | 12.75 | 12 | 18.32 | 11.31 | 13 | 0.3907 |
| REM% | 19.69 | 8.73 | 25 | 22.35 | 10.10 | 12 | 17.24 | 6.74 | 13 | 0.0737 |
| REM RMSSD | 46.94 | 20.09 | 23 | 44.65 | 22.50 | 11. | 49.04 | 18.35 | 12 | 0.6121 |
| SWS RMSSD | 49.73 | 23.39 | 23 | 40.86 | 22.55 | 11 | 57.86 | 21.93 | 12 | 0.0812 |

ASI-Anxiety Sensitivity Index (Reiss et al., 1986), ESS- Epworth Sleepiness Scale (Johns, 1991), FIRST– Ford Insomnia Response to Stress Test (Drake et al., 2004), GAD-7- Generalized Anxiety Disorder 7-item scale (Spitzer et al., 2006), IDT- Inventory of Depressive Symptomatology (Rush et al., 2003), IDT– The Inventory of Depressive Symptomatology (Rush et al., 2003), ISI- The Insomnia Severity Index (Bastien et al., 2001), PSQI- Pittsburgh Sleep Quality Index (Buysse et al., 1989), PSWQ- Penn State Worry Questionnaire (PSWQ)(Behar et al., 2003), PSWQ– Penn State Worry Questionnaire(Behar et al., 2003), REM- Rapid eye movement sleep, SE- sleep efficiency (SE), N1-N3- NREM sleep stages 1-3, SOL-sleep onset latency, STICSA-T- State-Trait Inventory for Cognitive and Somatic Anxiety-Trait Version (Gros et al., 2007; Ree et al., 2008; Ree et al., 2000), RMSSD- root mean square of successive R-R intervals TST- total sleep time, WASO-wake time after sleep onset,

**Table 3.**
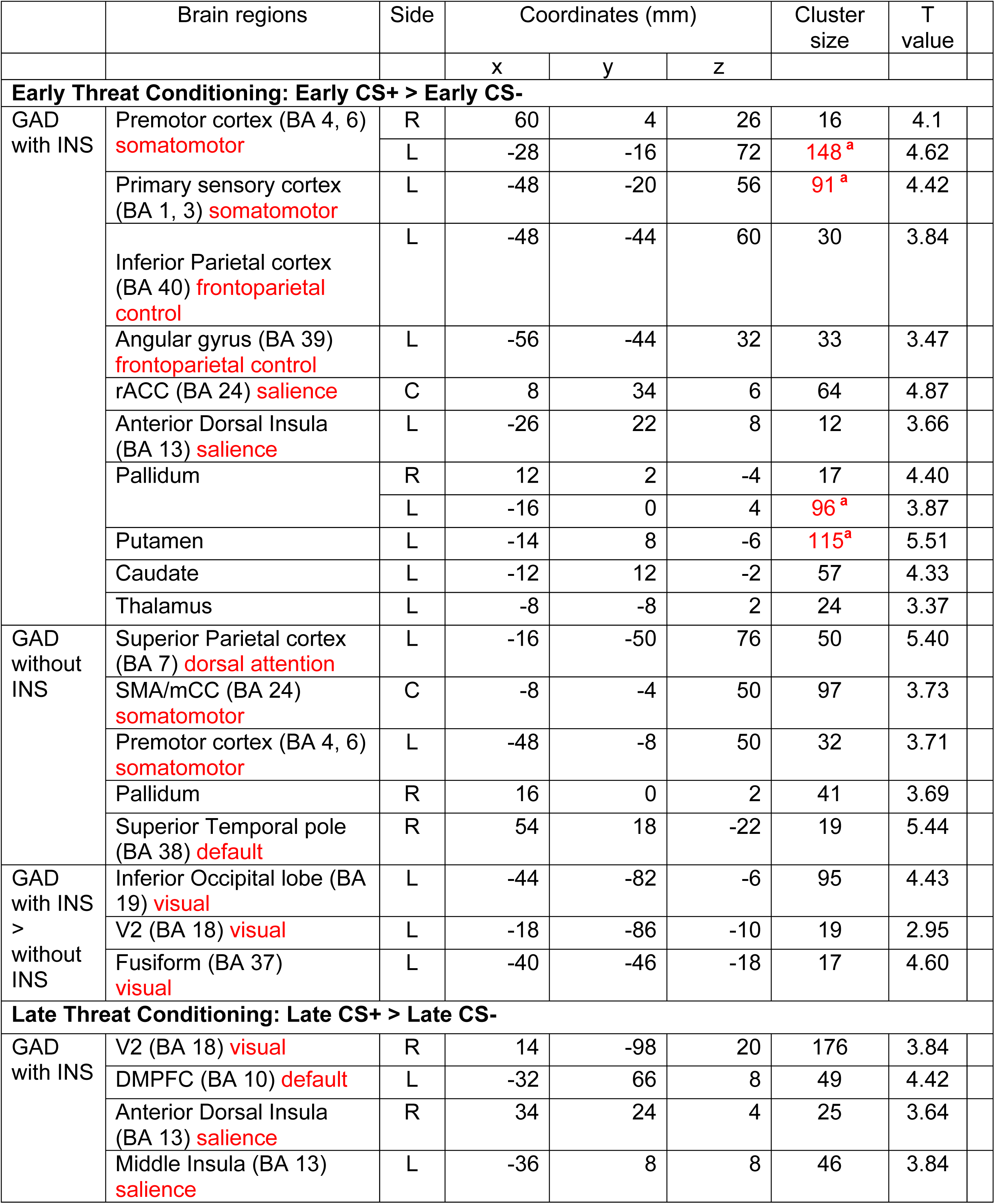
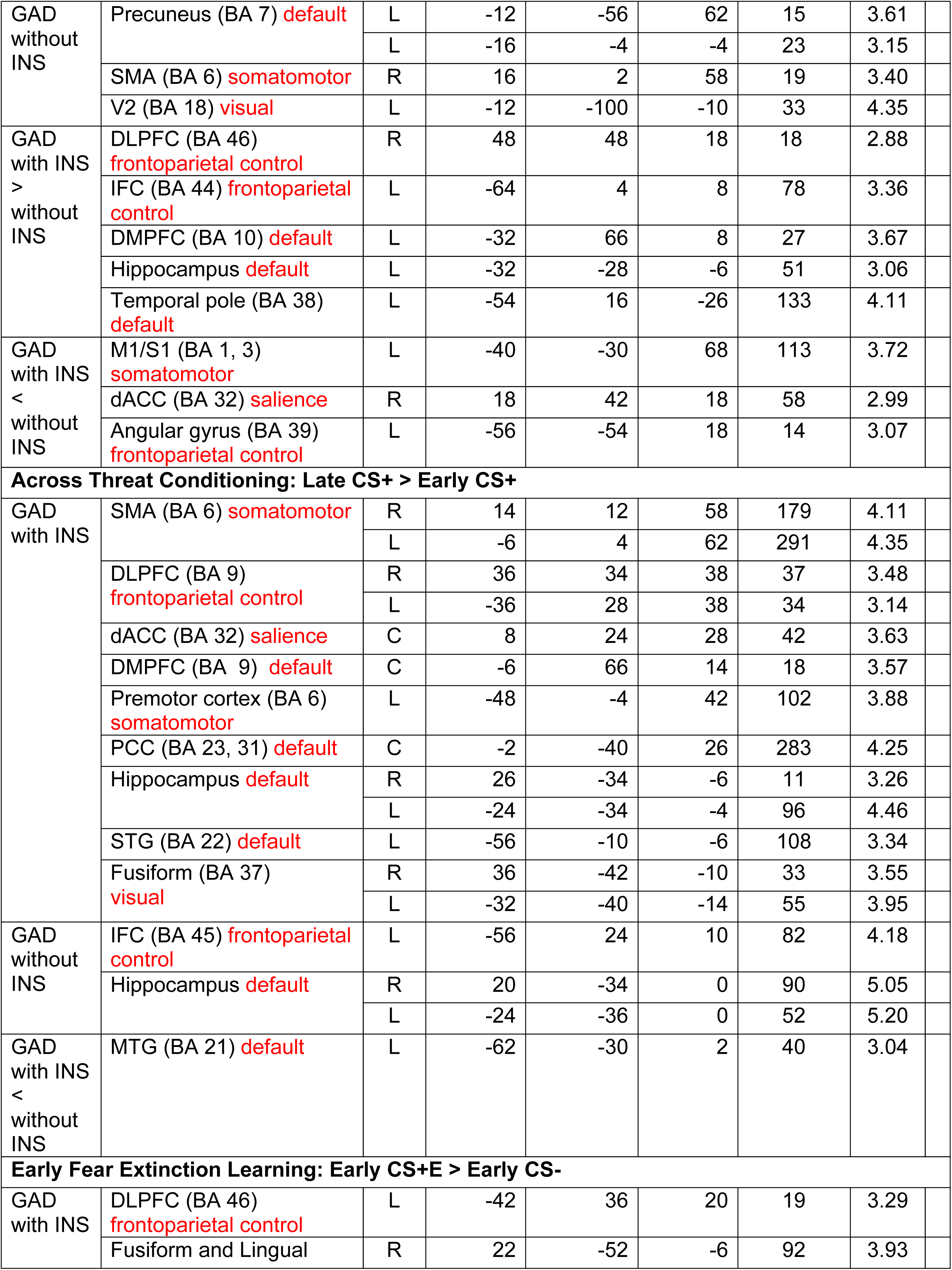
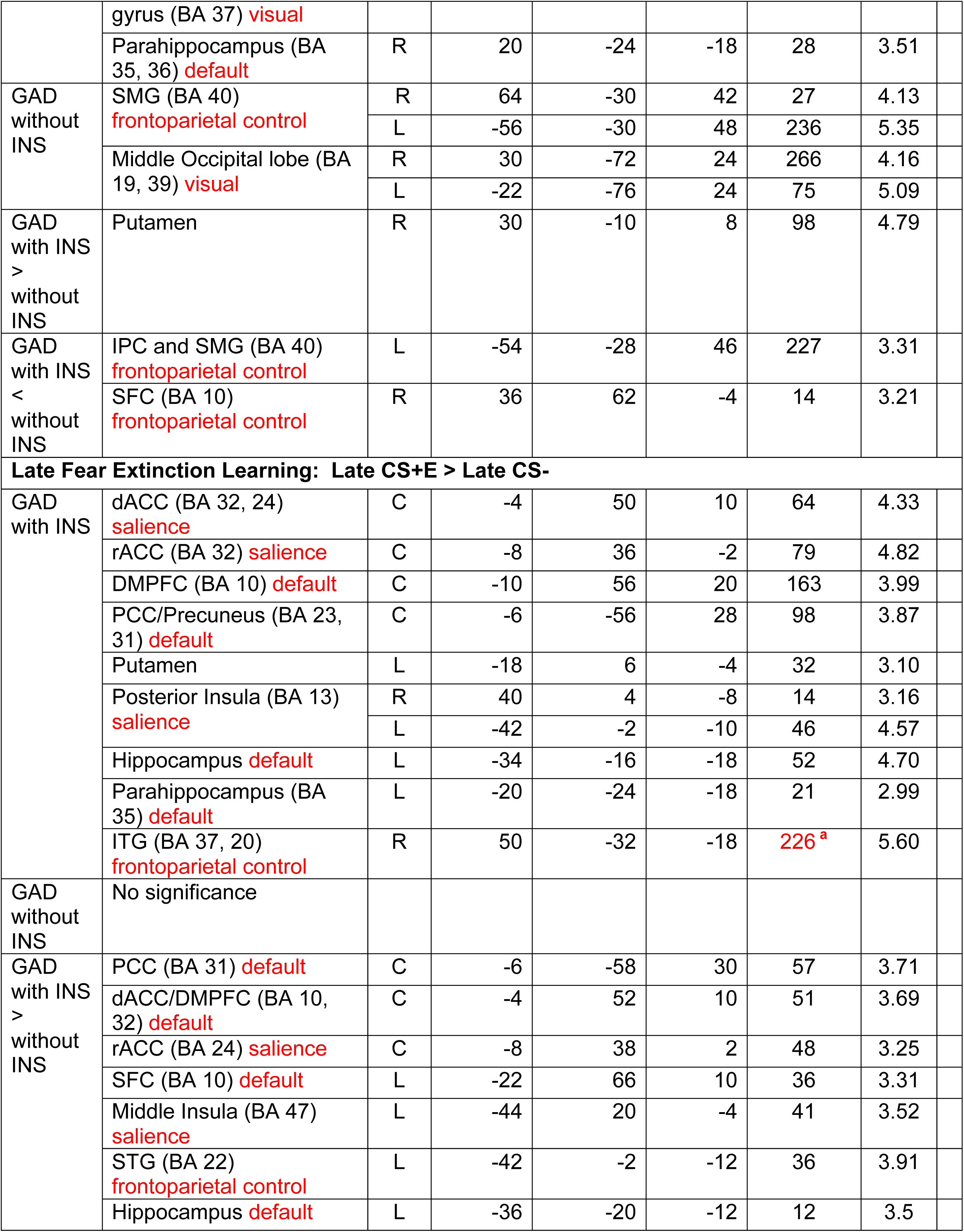
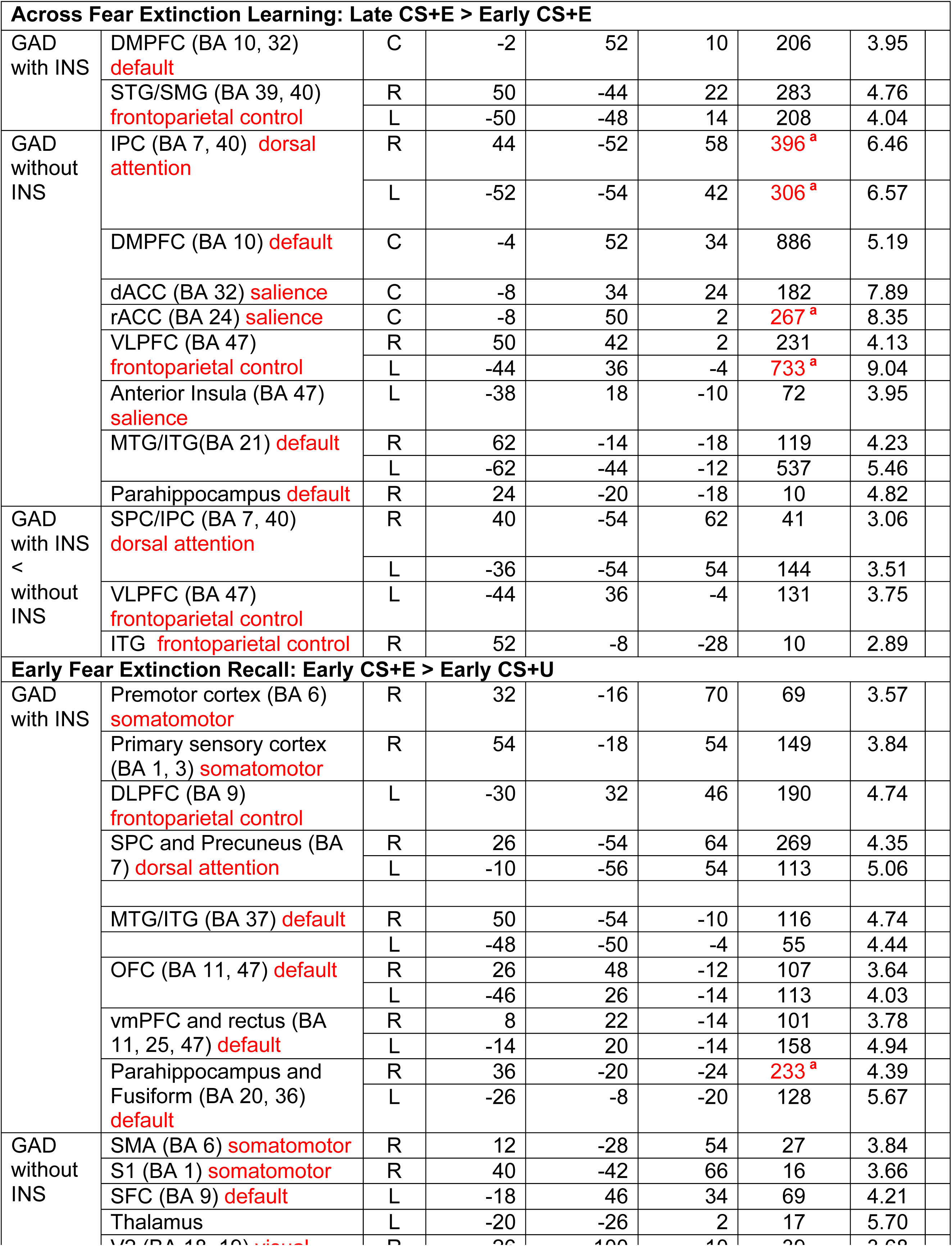
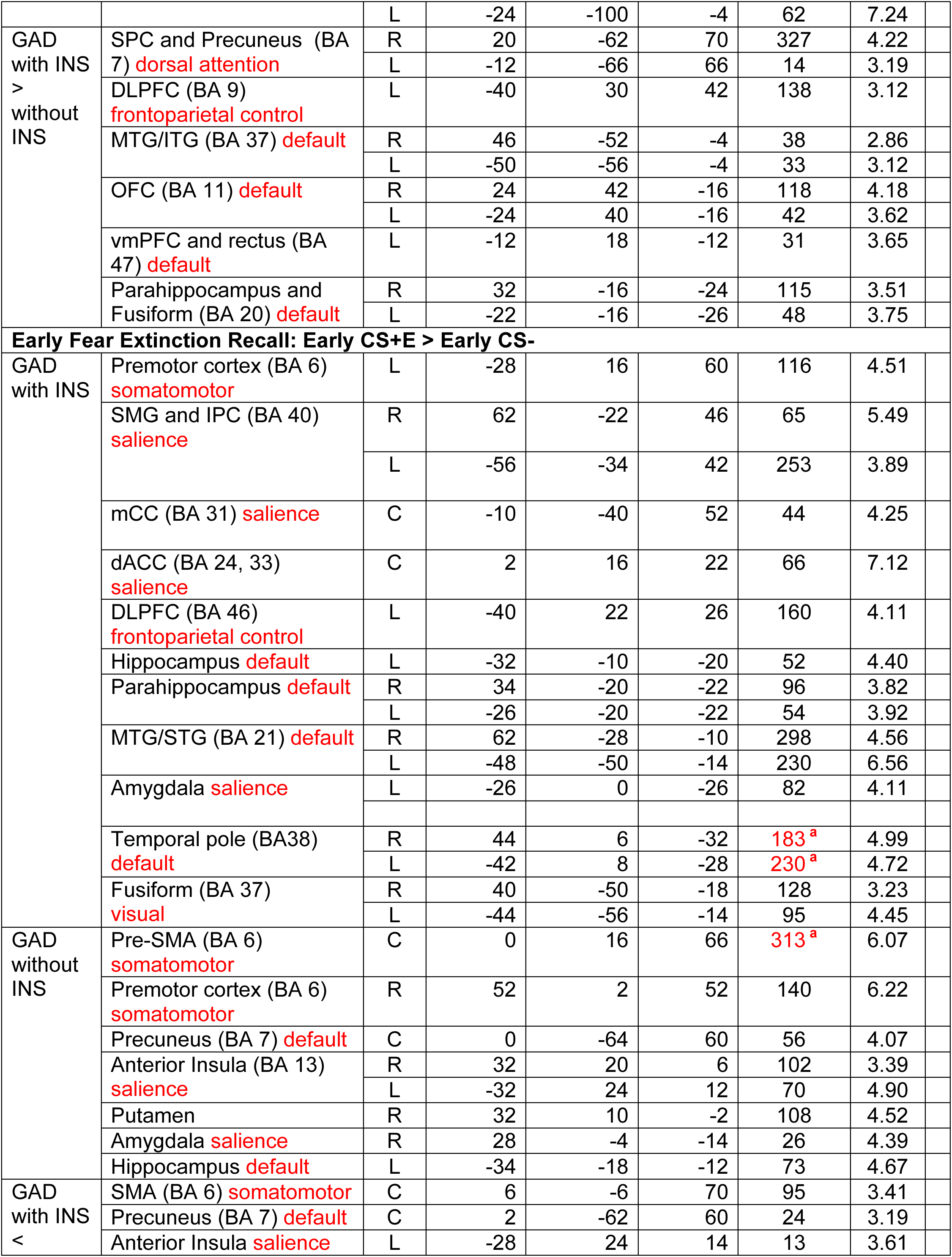
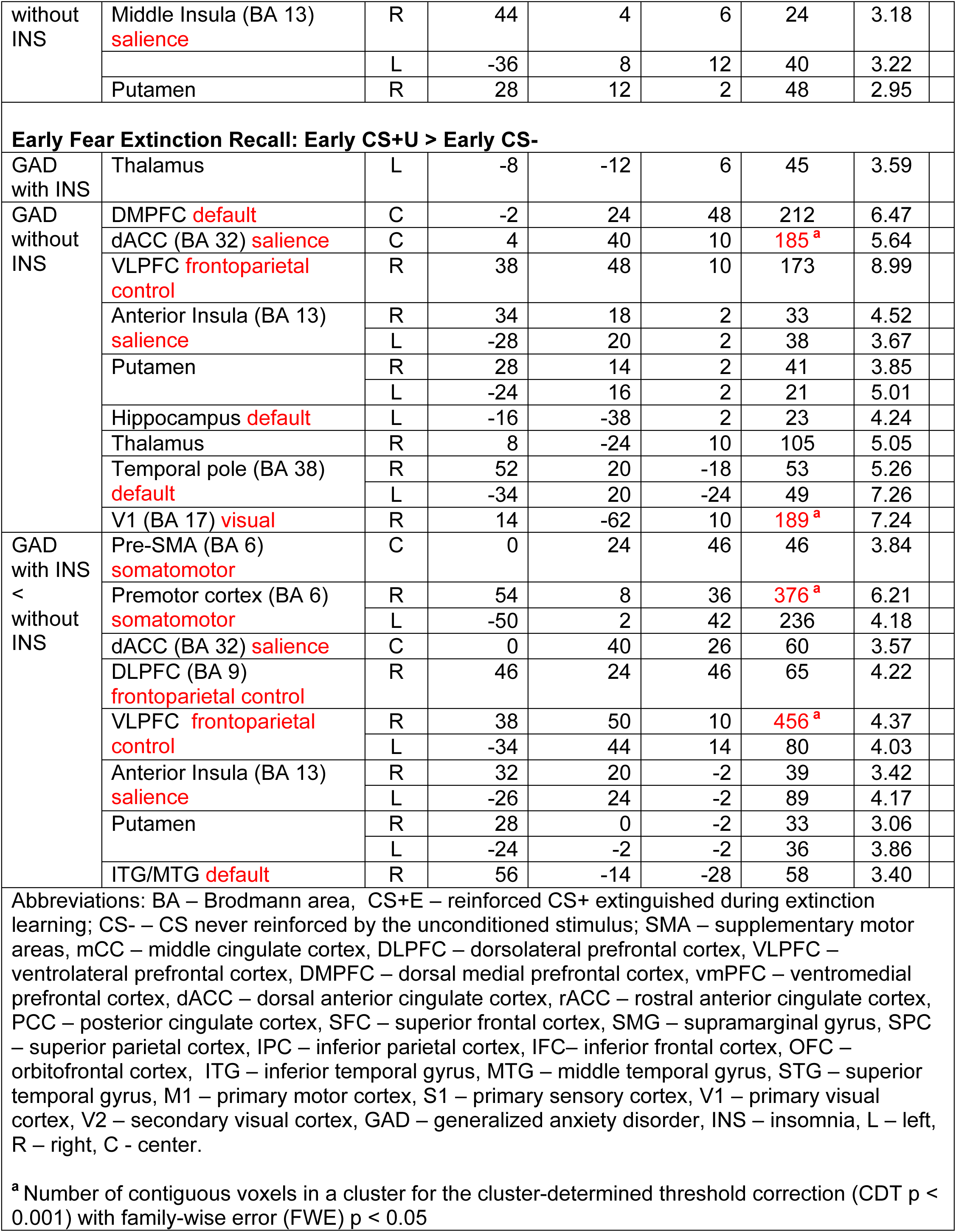
Neural activations to 9 contrasts, 3 per phase, within GAD groups and between GAD groups (GAD with INS and GAD without INS). Whole brain, uncorrected p < 0.005 with a contiguous voxel of 10. Peak Montreal Neurological Institute (MNI) coordinates, cluster size, and T value are reported for each cluster.

**Table 4.**
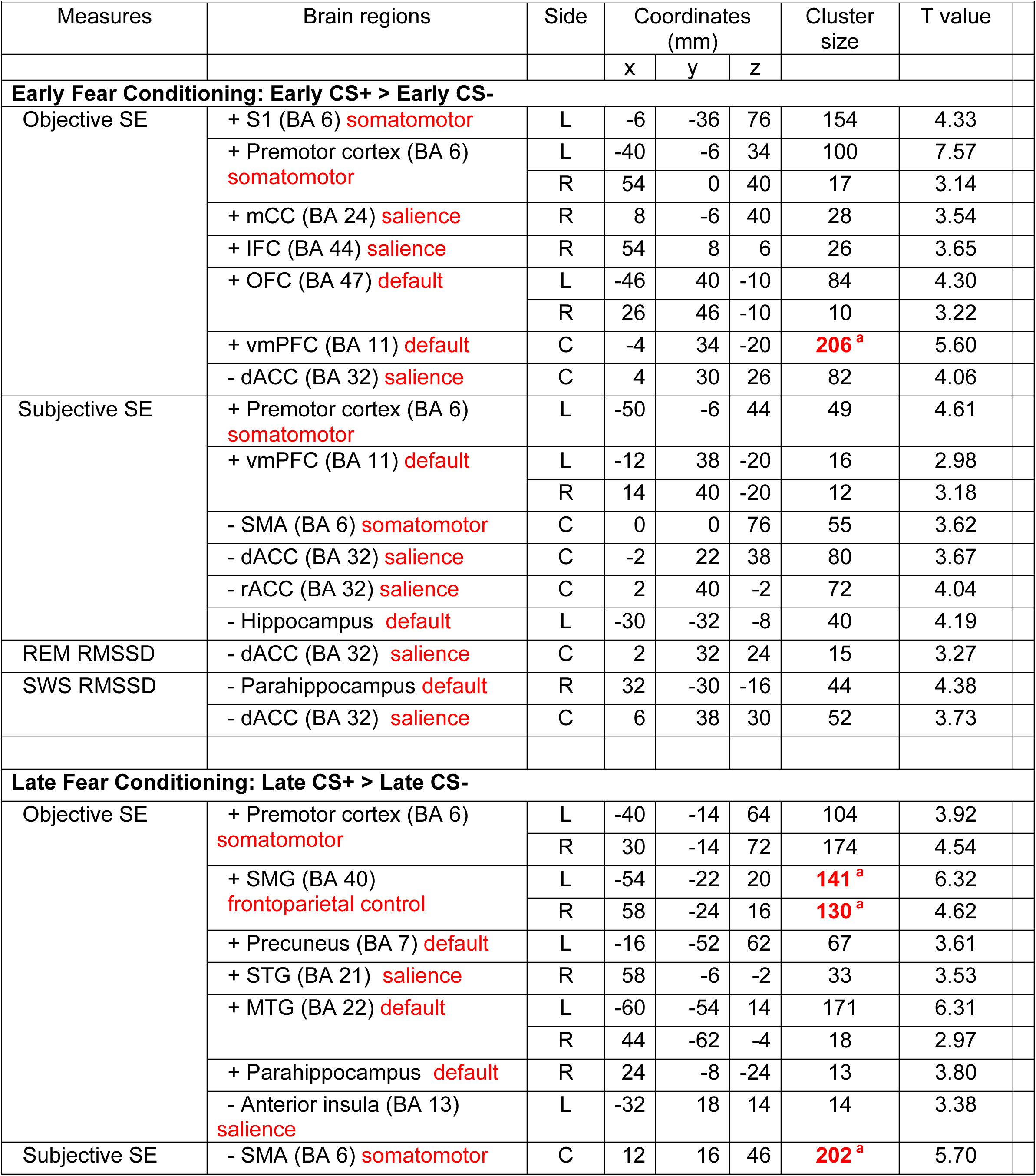
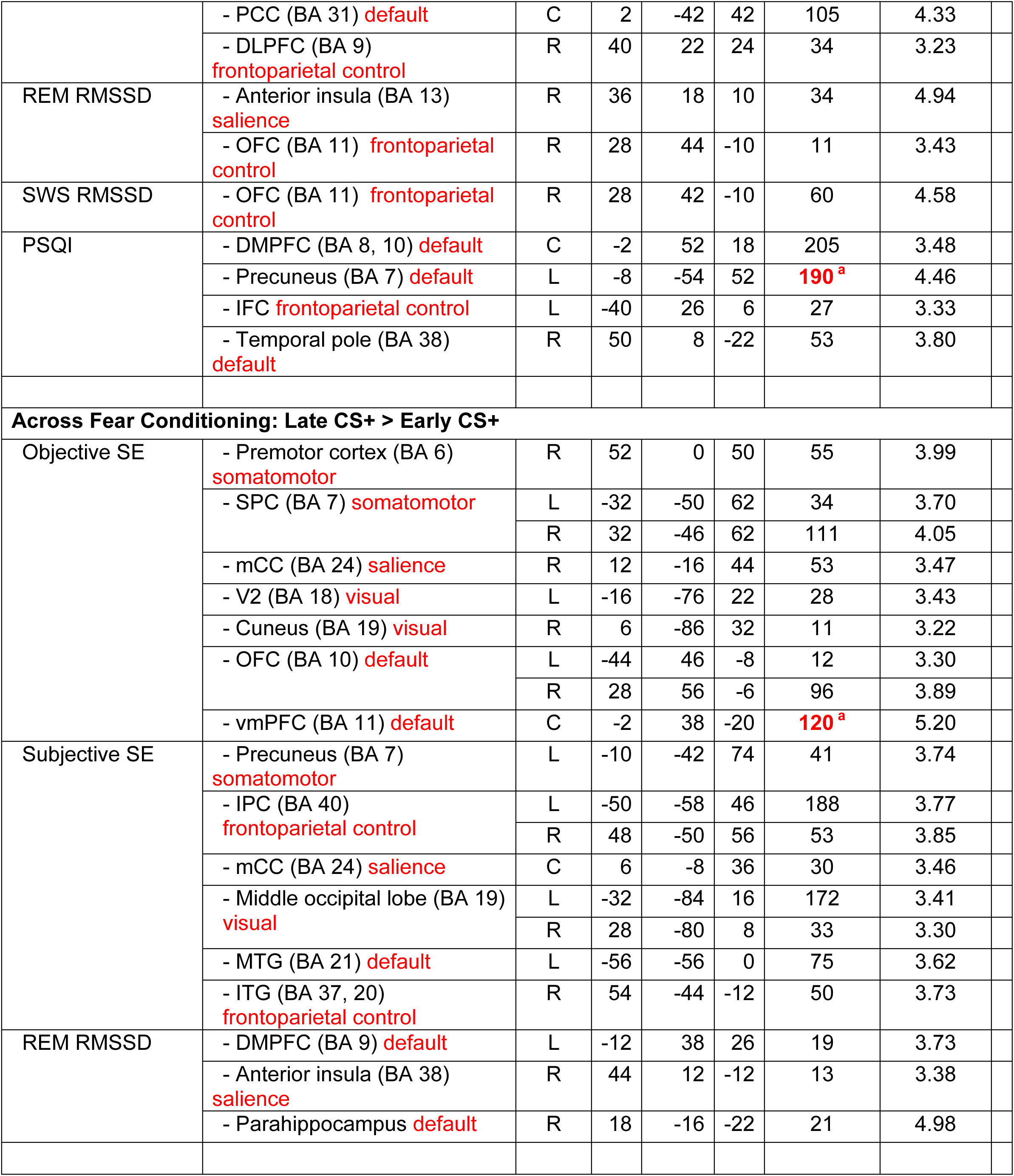
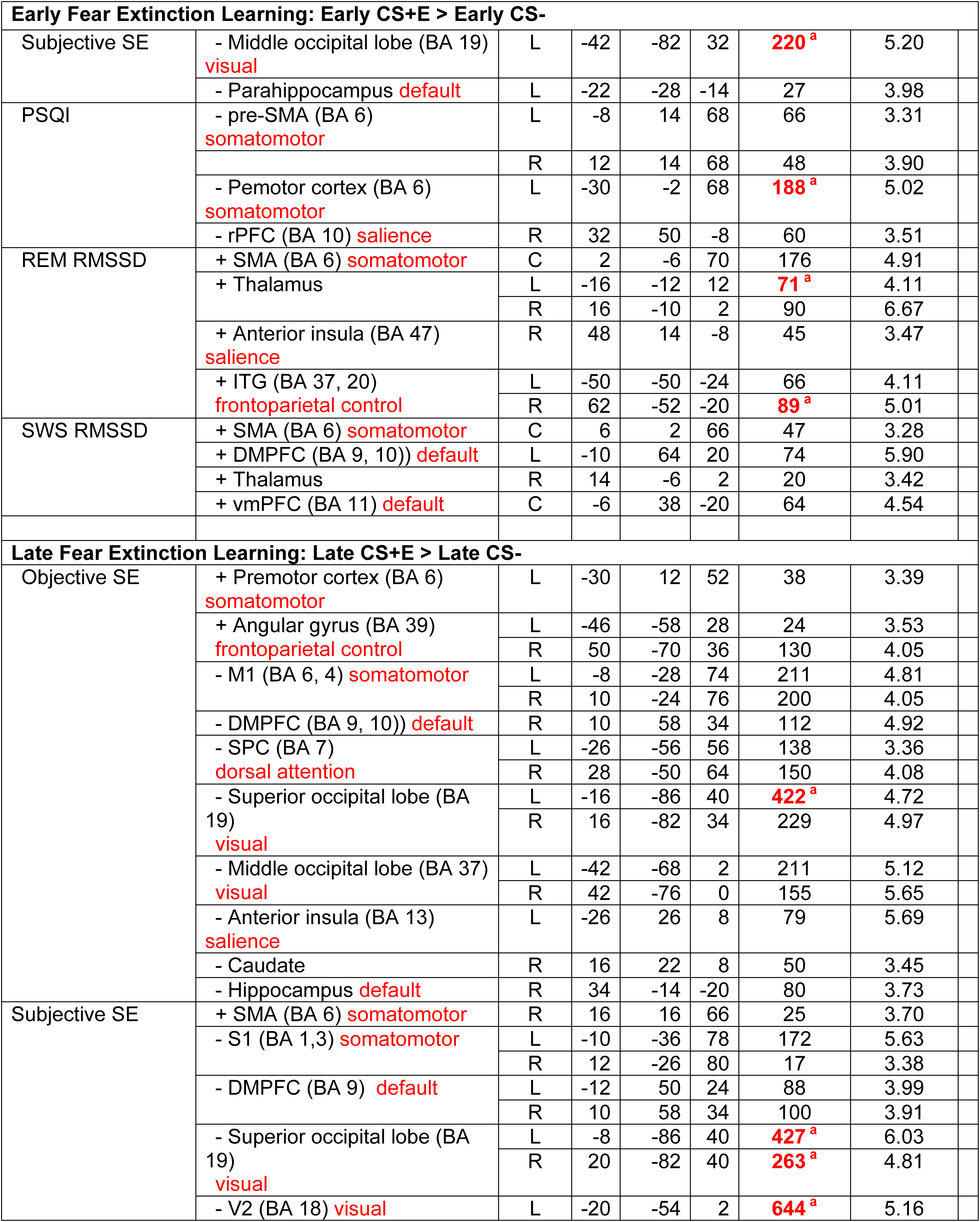
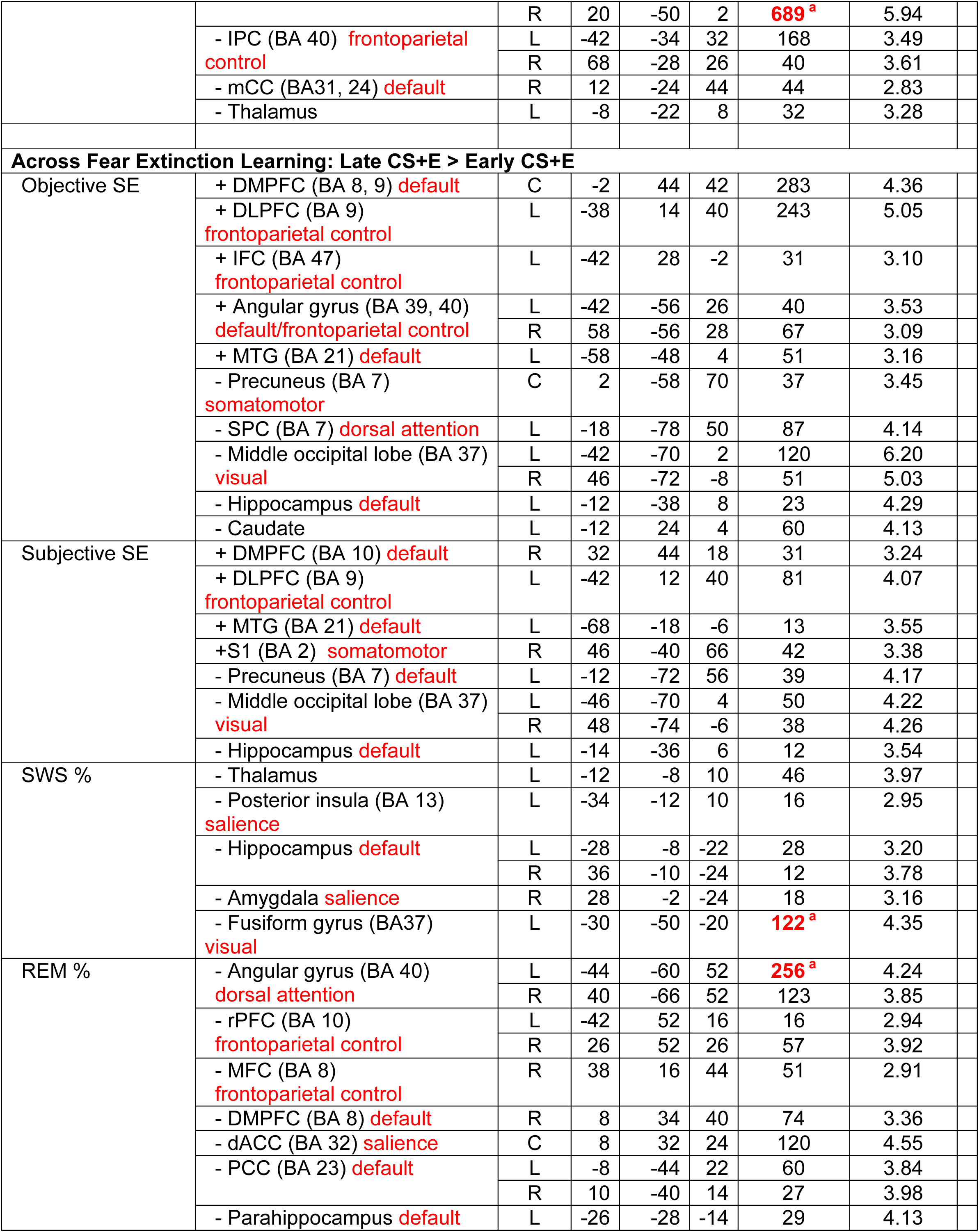
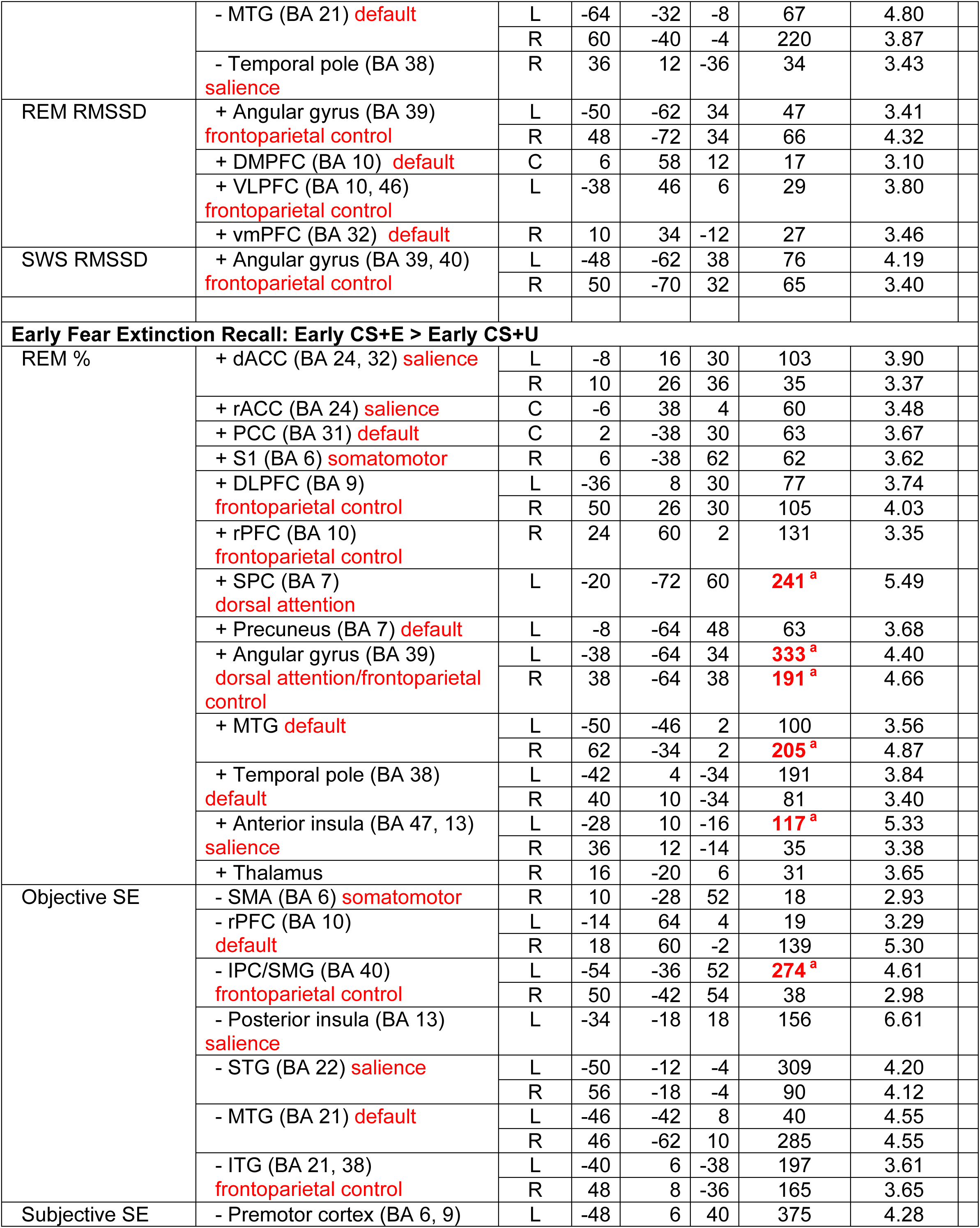
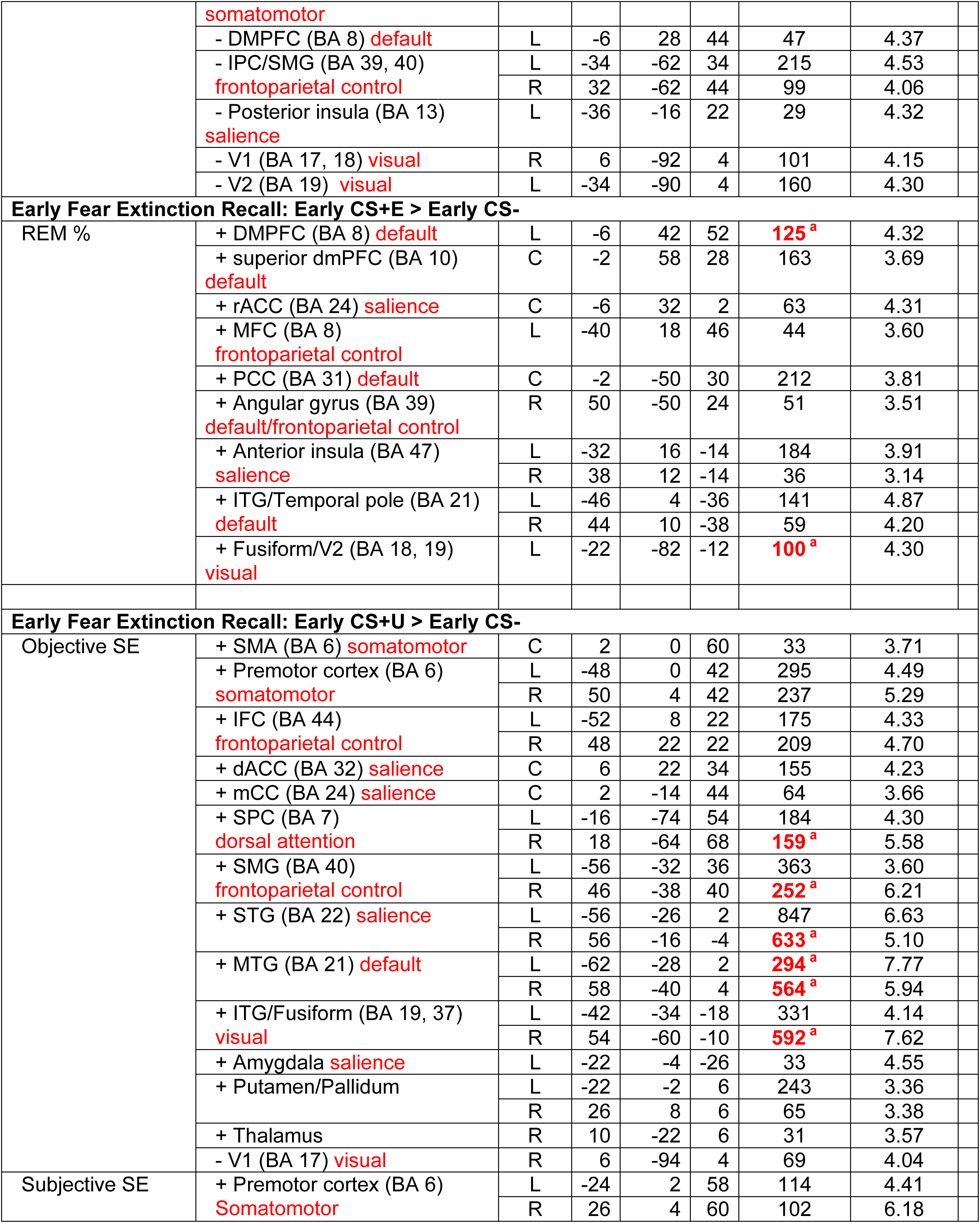
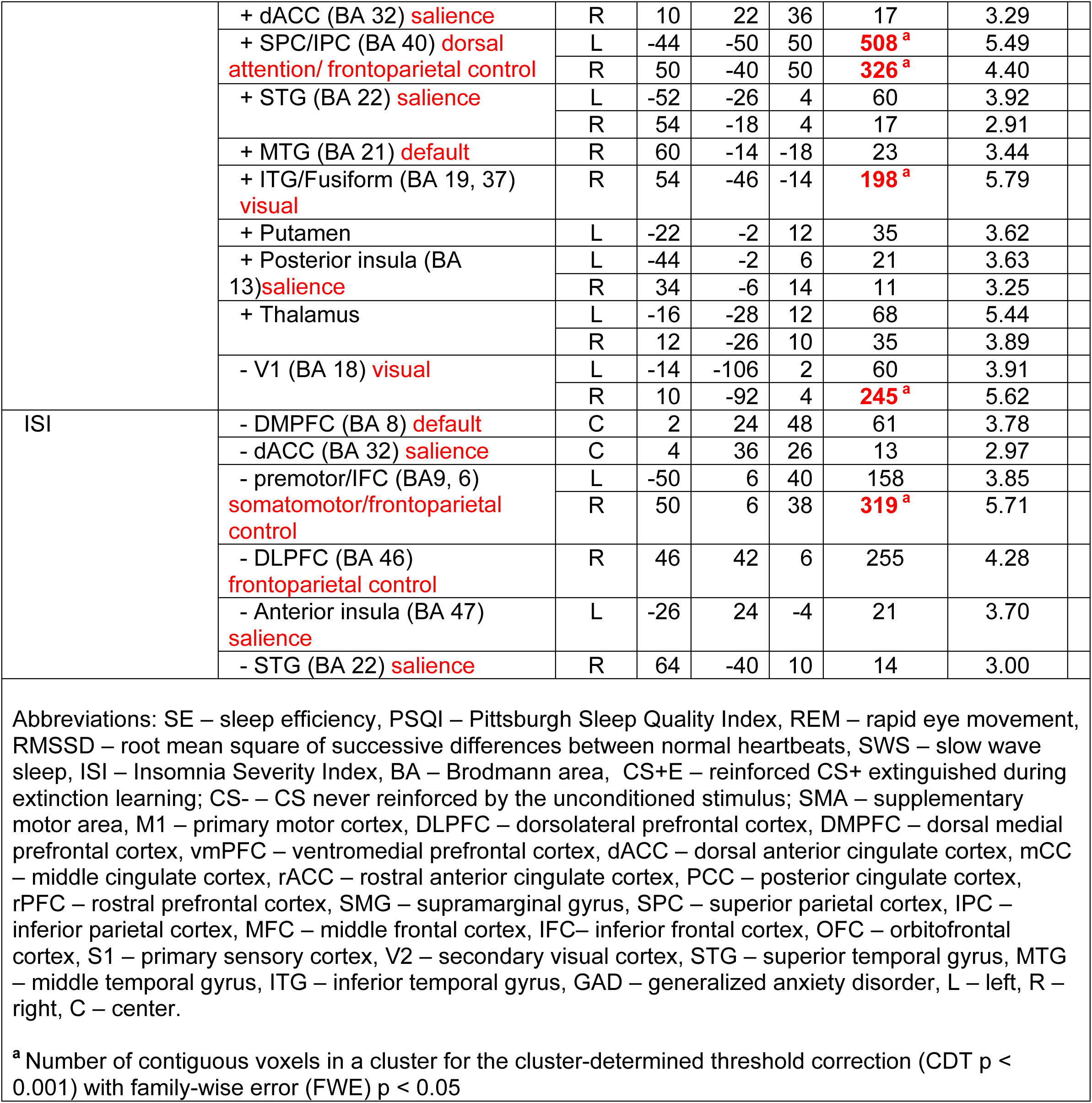
Multiple regression analyses to 9 contrasts, 3 per phase, within whole GAD. Whole brain, uncorrected p < 0.001 with a cluster-determining threshold FWE of p < 0.05. Peak Montreal Neurological Institute (MNI) coordinates, cluster size, and T value are reported for each cluster.

#### 2.4.3 Exclusion of specific scans based on image quality

Prior to second-level analyses, raw Echo Planar Imaging (EPI) data for all scans were examined using SPM’s “Check Registration” function (Di and Biswal, 2022), and specific scans were excluded when excessive signal intensity was observed. Given the limited sample size, all participants with good scanning data in an experimental phase were included in the analyses of that phase (i.e., pairwise deletion). In total, 6 Threat Conditioning scans and 3 scans each from Extinction Learning and Extinction Recall were excluded from second-level analyses. Scans from all 3 phases were obtained from 12 participants in the GAD+ID group and 10 from the GAD-ID group. Among the partial data sets for specific participants, Threat Conditioning and Extinction Learning scans only were obtained from 1 GAD+ID and 2 GAD-ID participants, Extinction Learning and Extinction Recall only from 2 GAD+ID and 1 GAD-ID participants, and Extinction Recall only from 1 GAD+ID participant.

### 2.5 Statistical analyses

Group comparisons and multiple regression analyses were carried out with Statistical Parametric Mapping (SPM12; Wellcome Trust Centre for Neuroimaging, www.fil.ion.ucl.ac.uk) implemented in MATLAB v2022a (The Mathworks Inc., Natick, Massachusetts, USA) using age and sex as covariates. For both group comparisons and multiple regression analyses, *low-threshold activations* used a cluster determining threshold (CDT) of *p* <0.005 with at least 10 contiguous voxels, whereas *high-threshold activations* used a CDT of *p* <0.001 and included only clusters surviving SPM12 family-wise error (FWE) correction at *p* <0.05. PSG sleep parameters used for multiple regressions with neural activations were derived from the Baseline night for activations measured during Conditioning and Extinction learning and from the Consolidation night for activations during Extinction Recall. Sleep, clinical, and demographic variables were compared between groups using independent sample t-tests (Table 2).

## 3. Results

### 3.1 Threat conditioning and extinction learning

As detailed in Supplemental Methods and Results, analyses of both psychophysiological response (skin conductance) and subjective ratings (shock expectancy) showed that both threat conditioning and its extinction were successfully achieved. A forthcoming report will detail objective and subjective ratings of threat conditioning, extinction learning and extinction recall in relation to sleep physiology.

### 3.2 Group characteristics

Of the 35 individuals who provided psychometric and sleep data, we obtained usable fMRI data from 30 (Table 2). Threat-Conditioning data were obtained from 26 (14 GAD+ID, 12 GAD-ID), Extinction Learning data were obtained from 29 (16 GAD+ID, 13 GAD-ID) and Extinction Recall data were obtained from 26 (15 GAD+ID, 11 GAD-ID). Table 2 provides mean group characteristics including only those individuals for whom fMRI data were available. Self-report and objective sleep data in Table 2 include measures to be analyzed in relation to neuronal activations (e.g., ISI, REM%) as well as additional measures (e.g., Ford Insomnia Response to Stress Test, REM Latency) provided to further characterize sleep in this sample. Group comparisons used independent-sample t-tests at an uncorrected alpha of p ≤ 0.05. For sleep quality measures, as expected, compared to GAD-ID, GAD+ID showed scores indicative of higher severity on the ISI, Pittsburgh Sleep Quality Index (PSQI), Ford Insomnia Response to Stress Test (FIRST), diary SE, diary SOL, actigraph SOL and a trend (p=0.056) for actigraph SE. Among psychometric measures, compared to GAD-ID, GAD+ID showed higher (greater severity) scores on the PSWQ and on the somatic subscale of the State-Trait Inventory of Cognitive and Somatic Anxiety-Trait Version (STICSA-T). There were no significant group differences among sleep-stage percentages.

### 3.3 Brain activation within and between groups

#### 3.3.1 *Across Threat Conditioning* (late>early CS+ contrast)

In GAD+ID, there was increased low-threshold activation in regions of the salience *(neuroanatomic region[s] =* dACC; *total number of voxels in network bilaterally* = 42 *network abbreviation* = SN), default (DMPFC, PCC, hippocampus, STG; 516 DMN), frontoparietal control (DLPFC; 71 FPCN), and somatomotor (SMA, PMC; 572 SM) networks (Fig. 1, 2, Table 3). However, no clusters reached high-threshold significance. In contrast, low-threshold activity in GAD-ID across this contrast showed activation in only the default (hippocampus; 142 DMN) and frontoparietal control (IFC; 82 FPCN) networks (Fig. 1, 2, Table 3). As in GAD+ID, no clusters reached high-threshold significance for this contrast. At the low threshold, GAD-ID was greater than GAD+ID in the default (MTG; 40 DMN) network (Table 3). No group differences were seen at the high threshold.

#### 3.3.2 Across *Extinction Learning* (late>early CS+E contrast)

In GAD+ID, there was increased low-threshold activation in the default (DMPFC; 206 DMN) and frontoparietal control (STG/SMG; 491 FPCN) networks, no clusters of which reached high-threshold significance (Fig. 1, 2, Table 3). However, in GAD-ID there was much more extensive low-threshold activation in salience (dACC/rACC, anterior insula; 521 SN), default (DMPFC, MTG, parahippocampus; 1552 DMN), frontoparietal control (VLPFC; 964 FPCN), and dorsal attention (IPC; 702 DAN) networks (Fig. 1, 2, Table 3). Clusters reached significant high-threshold activation for this contrast in default (bilateral IPC, left VLPFC) and salience (rACC) networks (Fig. 1, 2, Table 3). GAD-ID showed greater low-threshold activation than GAD+ID in two areas of the frontoparietal control network (VLPFC, ITG; 141 FPCN) and the dorsal attention network (SPC/IPC; 185 DAN), however, these differences did not reach high-threshold significance (Table 3).

#### 3.3.3 During early Threat Extinction Recall

##### 3.3.3.1 Early CS+E>CS+U contrast

In GAD+ID, there was increased low-threshold activation in default (MTG, OFC, vmPFC, parahipocampus; 883 DMN), frontoparietal control (DLPFC; 190 FPCN), dorsal attention (SPC/Precuneus; 382 DAN), and somatomotor (PMC, S1; 218 SM) networks (Fig. 1, 2, Table 3). Also, in GAD+ID, activation was significant at the high-threshold in the right parahippocampus (DMN) (Fig. 2, Table 3). For this contrast, GAD-ID showed low-threshold activation in default (SFC; 69 DMN), and somatomotor (SMA, S1; 43 SM) networks, and no areas reached high-threshold significance (Fig. 1, 2, Table 3). GAD+ID showed greater low-threshold activation than GAD-ID in default (vmPFC, MTG, OFC, and parahippocampus; 766 DMN), dorsal attention (SPC/precuneus; 341 DAN), and frontoparietal control (DLPFC; 138 FPCN) networks, however, none of these areas reached high-threshold significance (Fig. 2, Table 3).

##### 3.3.3.2 Early CS+E>CS- contrast

In GAD+ID there was increased low-threshold activation in salience (SMG, mCC, dACC, amygdala; 510 SN), default (hippocampus, parahippocampus, MTG, temporal pole; 1143 DMN), frontoparietal control (DLPFC; 160 FPCN), somatomotor (PMC; 116 SM) and visual (fusiform; 223 VN) networks (Fig. 1, 2, Table 3). In GAD+ID, there was also high-threshold activation in the bilateral temporal pole (DMN) (Fig. 2, Table 3). In contrast, low-threshold activity in GAD-ID for this contrast showed activation in salience (anterior insula, amygdala; 198 SN), default (precuneus, hippocampus; 129 DMN), and somatomotor (pre-SMA, PMC; 453 SM) networks (Fig. 1, 2, Table 3). In GAD-ID, high-threshold activation was reached in the pre-SMA (SM) (Fig. 2, Table 3). Low-threshold activation for this contrast was greater in GAD-ID than GAD+ID in salience (left anterior and bilateral middle insula; 77 SN), default (precuneus; 24 DMN) and somatomotor (SMA; 95 SM) networks, and subcortically in the right putamen, however, none of these areas reached high-threshold significance (Table 3).

##### 3.3.3.3 Early CS+U>CS- contrast

In GAD+ID, increased low-threshold activation was seen only subcortically in the left thalamus (Table 3). However, in GAD-ID, there was much more extensive activation in salience (dACC, anterior insula; 256 SN), default (DMPFC, hippocampus, temporal pole; 337 DMN), and frontoparietal control (VLPFC; 173 FPCN) networks (Fig. 1, 2, Table 3). In GAD-ID there was also significant high-threshold activation in dACC and right primary visual cortex (Fig. 2, Table 3). GAD-ID showed greater low-threshold activation than GAD+ID in salience (dACC, anterior insula; 188 SN), frontoparietal control (DLPFC, VLPFC; 601 FPCN) and somatomotor (pre-SMA, PMC; 658 SM) networks and subcortically in the putamen (Table 3). For this contrast, GAD-ID activation was greater than GAD+ID at high-threshold significance in the right VLPFC (FPCN) and right PMC (SM) (Table 3).

### 3.4 Association of sleep measures with neural activations

Among the entire sample (GAD+ID and GAD-ID combined) sleep measures that were examined for associations with neural activations to the 9 contrasts, included ISI, Pittsburgh Sleep Quality Index (PSQI), mean Objective SE (actiwatch), mean Subjective SE (diary), REM and SWS as percent of TST, and heart-rate variability in REM and SWS (Table 4, Figs. 3–5, Supplemental Materials). Among the measures shown in Table 2, these specific ones were selected, pre-hoc, as they best addressed hypotheses 2 and 3.

**Figure 4.**
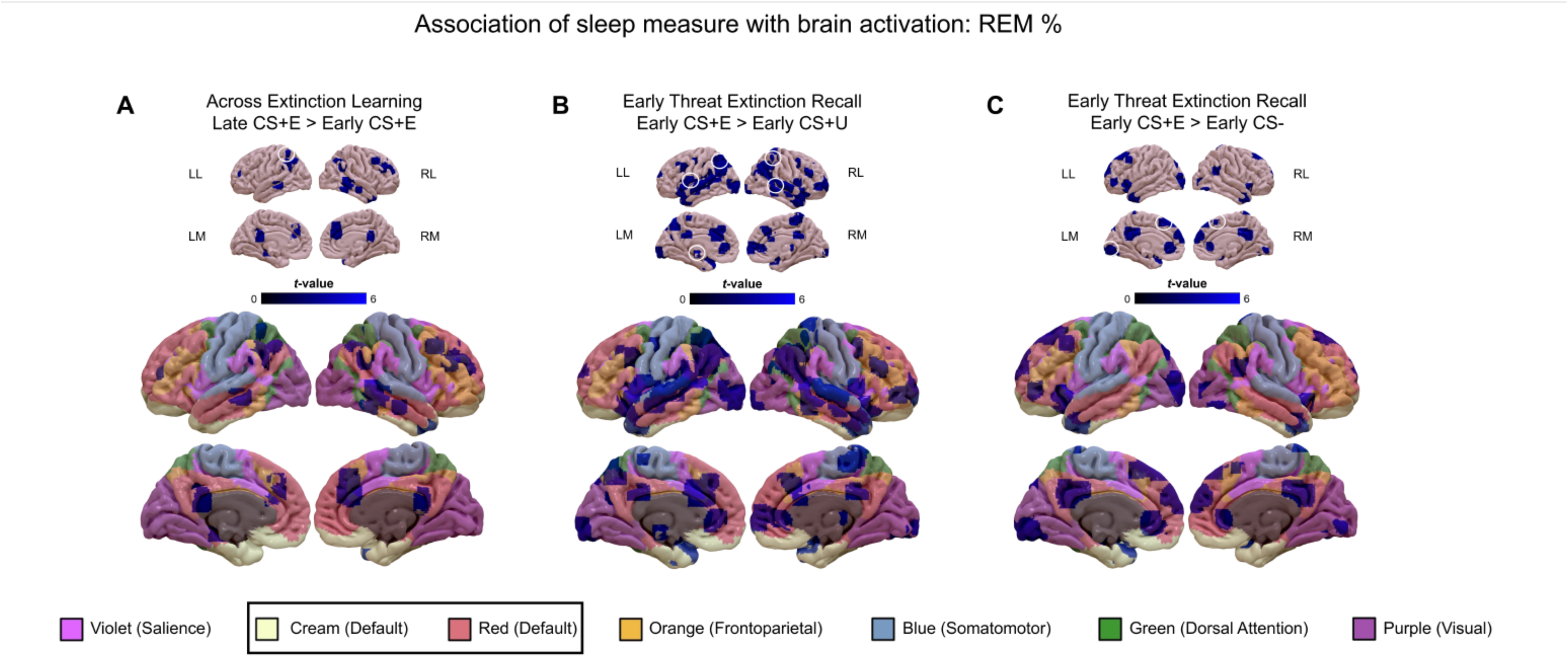
Associations of activations in the whole-sample (GAD+ID and GAD-ID) with **REM %** across the Extinction Learning phase (panel A), and the early Threat Extinction Recall phase (panels B and C). In each of panels A-E, *low-threshold activations* (cluster-determining threshold [CDT] of *p* <0.005 and at least 10 contiguous) are indicated in graded blues on both the upper sets of 8 smaller images and on the lower sets of 8 larger images which also display the 7 large-scale networks reported by Yeo et al. (Uddin et al., 2019; Yeo et al., 2011) color coded as indicated by the bottom legend. Clusters circled in white on the upper (smaller) images indicate *high-threshold activations* that survived family-wise error correction at *p* <0.05 with a CDT of *p* <0.001. Abbreviations: CS+, conditioned stimulus reinforced by the unconditioned (shock) stimulus; CS-, nonreinforced conditioned stimulus; CS+E, CS+ extinguished during Extinction Learning; CS+U, un-extinguished CS+; GAD, generalized anxiety disorder; ID, insomnia disorder; LL, left lateral view; RL, right lateral view; LM, left medial view; RM, right medial view.

**Figure 5.**
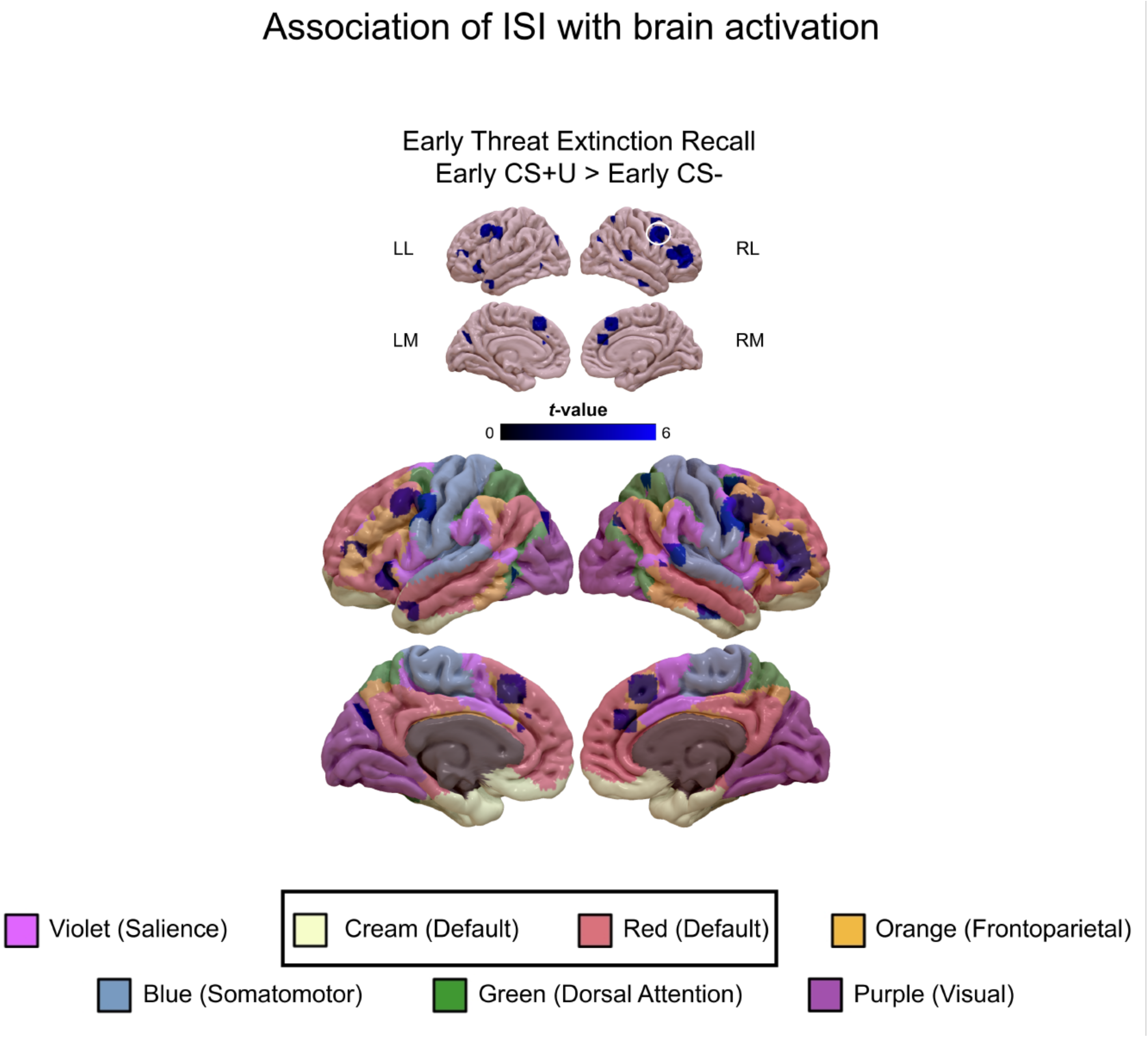
Associations of activations in the whole-sample (GAD+ID and GAD-ID) with the **Insomnia Severity Index (ISI)** during the early Threat Extinction Recall phase (early CS+U>early CS-). *low-threshold activations* (cluster-determining threshold [CDT] of *p* <0.005 and at least 10 contiguous) are indicated in graded blues on both the upper sets of 8 smaller images and on the lower sets of 8 larger images which also display the 7 large-scale networks reported by Yeo et al. (Uddin et al., 2019; Yeo et al., 2011) color coded as indicated by the bottom legend. Clusters circled in white on the upper (smaller) images indicate *high-threshold activations* that survived family-wise error correction at *p* <0.05 with a CDT of *p* <0.001. Abbreviations: CS+, conditioned stimulus reinforced by the unconditioned (shock) stimulus; CS-, nonreinforced conditioned stimulus; CS+E, CS+ extinguished during Extinction Learning; CS+U, un-extinguished CS+; GAD, generalized anxiety disorder; ID, insomnia disorder; LL, left lateral view; RL, right lateral view; LM, left medial view; RM, right medial view.

#### 3.4.1 Objective and subjective sleep efficiency (SE)

##### 3.4.1.1 Threat conditioning

Across Threat Conditioning (late CS+>early CS+ contrast), as **objective SE** increased, low-threshold activation decreased in salience (mCC; 53 SN), default (OFC, vmPFC; 228 DMN), somatomotor (PMC, SPC; 200 SM), and visual (V2, cuneus; 39 VN) networks. Only activation in vmPFC (DMN) significantly decreased at high-threshold with increased objective SE (Fig. 3, Table 4.). As, **subjective SE** increased, low-threshold activation decreased in salience (mCC; 30 SN), default (MTG; 75 DMN), frontoparietal control (ITG, IPC; 291 FPCN), somatomotor (precuneus; 41 SM), and visual (BA19; 205 VN) networks. However, no activations were associated with subjective SE at high-threshold significance (Table 4).

##### 3.4.1.2 Extinction Learning

Across Extinction Learning (late CS+E>early CS+E contrast), as **objective SE** increased, there was increased low-threshold activation in default (DMPFC, AG, MTG; 441 DMN) and frontoparietal control (DLPFC, IFC; 274 FPCN) networks, but decreased activation in somatomotor (Precuneus; 37 SM), dorsal attention (SPC; 87 DAN), and visual (MOL; 171 VN) networks. However, no activations were associated with objective SE at high-threshold significance (Fig. 3, Table 4). As, **subjective SE** increased, there was increased low-threshold activation in default (DMPFC, MTG; 44 DMN), frontoparietal <u>control</u> (DLPFC; 81 FPCN) and somatomotor (S1; 42 SM) networks, but decreased activation in default (hippocampus, precuneus; 51 DMN) and visual (MOL; 88 VN) networks. However, no activations were associated with subjective SE at high-threshold significance (Table 4)

##### 3.4.1.3 Extinction Recall

###### 3.4.1.3.1 Early CS+E>early CS+U contrast

During early Extinction Recall, for the early CS+E>early CS+U contrast, as **objective SE** increased, there was decreased low-threshold activation in default (PFC, MTG; 483 DMN), frontoparietal control (IPC and ITG; 674 FPCN), salience (PIC, STG; 555 SN) and somatomotor (SMA; 18 SM) networks. Decreased activation with increased objective SE reached high-threshold significance in the left IPC (FPCN) (Fig. 3, Table 4). As **subjective SE** increased, there was decreased low-threshold activation in salience (PIC; 29 SN), default (DMPFC; 47 DMN), frontoparietal control (IPC, SMG; 314 FPCN), and visual (V1, V2; 261 VN) networks. However, no activations were associated with subjective SE at high-threshold significance (Table 4).

###### 3.4.1.3.2 Early CS+U>early CS- contrast

For the early CS+U>early CS- contrast, as **objective SE** increased, there was increased low-threshold activation in salience (dACC, mCC, STG, amygdala; 1732 SN), frontoparietal control (IFC, SMG; 999 FPCN), dorsal attention (SPC; 343 DAN), somatomotor (SMA, PMC; 565 SM) and visual (ITG/fusiform; 923 VN) networks, but decreased activation in visual network (V1; 69 VN). Increased activations with increased objective SE reached high-threshold significance in right SPC (DAN), right SMG (FPCN), right STG (SN), bilateral MTG (DMN), and right ITG/fusiform(VN) (Fig. 3, Table 4). Similarly, as **subjective SE** increased, there was increased low-threshold activation in salience (dACC, STG, PIC;126 SN), default (MTG; 23 DMN), frontoparietal control/dorsal attention (IPC/SPC; 834 FPCN/DAN), somatomotor (PMC; 216 SM) and visual (ITG/fusiform; 198 VN) networks as well as the subcortex (putamen, thalamus; 138), but decreased activation in visual network (V1; 305 VN). Increased activations with increased SE reached high-threshold significance in bilateral IPC/SPC (FPCN/DAN) and right fusiform (VN). Decreased SE reached high-threshold significance in bilateral V1 (VN) (Table 4).

#### 3.4.2 REM as a percent of total sleep time

##### 3.4.2.1 Extinction Learning

Across Extinction Learning, As **REM %** increased, there was decreased low-threshold activation in default (DMPFC, PCC, parahippocampus, MTG; 477 DMN), frontoparietal control (rPFC, MFC; 124 FPCN), dorsal attention (AG; 379 DAN), and salience (dACC, temporal pole;ISCU 154 SN) networks. Decreased activation with increased REM% reached high-threshold significance in the left AG (DAN) (Fig. 4, Table 4).

##### 3.4.2.3 Extinction Recall

###### 3.4.2.1.1 Early CS+E>early CS+U contrast

During Extinction Recall, for the early CS+E>early CS+U contrast, as REM % increased, there was increased low-threshold activation in salience (dACC, rACC, AIC; 350 SN), default (PCC, precuneus, MTG, temporal pole; 703 DMN), frontoparietal control (DLPFC, rPFC; 313 FPCN), dorsal attention (SPC; 241 DAN), and frontoparietal control/dorsal attention (AG; 524 FPCN/DAN) networks. Increased activations with increased REM% reached high-threshold significance in left SPC (DAN), bilateral AG (FPCN), right MTG (DMN), and left AIC (SN) (Fig. 4, Table 4.).

###### 3.4.2.1.2 Early CS+E>early CS- contrast

At Extinction Recall, for the early CS+E>early CS- contrast, as **REM %** increased, there was increased low-threshold activation in, default (DMPFC, PCC, AG, ITG/temporal pole; 751 DMN), salience (rACC, AIC; 283 SN), frontoparietal control (MFC; 44 FPCN), and visual (fusiform, 100 VN) networks. Increased activations with increased REM% reached high-threshold significance in left DMPFC (DMN) and left fusiform gyrus (VN) (Fig. 4, Table 4).

#### 3.4.3 SWS as percent of total sleep time

Across Extinction Learning, as **SWS %** increased, there was decreased low-threshold activation in salience (PIC, amygdala; 34 SN), default (bilateral hippocampus; 40 DMN) and visual (fusiform gyrus; 122 VN) networks. Decreased activation with increased SWS % reached high-threshold significance in the left fusiform gyrus (VN) (Table 4.)

#### 3.4.4 HRV (RMSSD) in REM and SWS

##### 3.4.4.1 HRV in REM

###### 3.4.4.1.1 HRV and Threat Conditioning

Across Threat Conditioning, as **REM RMSSD** increased, there was decreased low-threshold activation in the salience (AIC; 13 SN) and default (DMPFC, parahippocampus; 40 DMN) networks. However, no activations were associated with subjective REM RMSSD at high-threshold significance (Table 4).

###### 3.4.4.1.2 HRV and Extinction Learning

Across Extinction Learning, as **REM RMSSD** increased, there was increased low-threshold activation in the default (DMPFC, vmPFC; 44 DMN) and frontoparietal control (AG, VLPFC; 142 FPCN) networks. Similarly, as SWS RMSSD increased, there was increased activation in the frontoparietal control network (AG; 141 FPCN). However, no activations were associated with subjective SE at high-threshold significance (Table 4).

#### 3.4.5 Insomnia Severity Index

At Extinction Recall, for the early CS+U>early CS- contrast, as **ISI** severity increased, there was decreased low-threshold activation in default (DMPFC; 61 DMN), salience (dACC, anterior insula, STG; 48 SN), somatomotor (PMC/IFC; 477 SM), and frontoparietal control (DLPFC; 255 FPCN) networks. Decreased activation with increased ISI reached high-threshold significance in the right PMC/IFC (SM/FPCN) (Fig. 5, Table 4).

## 4. Discussion

### 4.1 Summary of results

Neural activations to threat conditioning, extinction learning, and extinction recall were compared between persons diagnosed with GAD who had moderate to severe ID (GAD+ID) and those without ID or with mild ID (GAD-ID). Relationships of these activations with specific sleep measures known to be abnormal in anxiety-related disorders were also examined. Both group comparisons and whole-sample sleep-parameter relationships with neural activations were first detected at a low-threshold that showed the extent to which contrasts engaged large-scale brain networks but presumably had high Type-1 (false positive) error. Second, they were examined with FWE correction (high-threshold) that minimized Type-1 but risked high Type-2 (false negative) errors. Some contrasts but not others supported our 3 following hypotheses that: 1. GAD+ID vs. GAD-ID would show more activation in SN and less in regulatory networks; 2. regulatory regions would activate more, and SN areas less with better sleep quality, as well as with greater REM and SWS percent; and 3. regulatory regions would activate more and SN areas less with greater parasympathetic tone (HRV) in REM and SWS,. As expected, GAD+ID showed poorer self-reported sleep quality and greater worry and somatic anxiety (Table 2).

### 4.2 Comparison of activations in GAD+ID vs. GAD-ID

#### 4.2.1 Summary

Across the acquisition of threat conditioning, low threshold activations to the reinforced stimulus (CS+) increased more within the GAD+ID than within the GAD-ID group. In contrast, increased low-threshold activations to the CS+ across Extinction Learning were greater within the GAD-ID group reaching high-threshold (FWE) significance in the DMN and SN. At Extinction Recall, greater low-threshold activation became more extensive within the GAD+ID than within the GAD-ID group when contrasting the extinguished (CS+E) stimulus with either the un-extinguished (CS+U) or the non-reinforced (CS-) stimuli, an association reaching high-threshold significance in the DMN. In contrast, greater activation in the CS+U > CS- contrast was much more extensive within the GAD-ID group, reaching high-threshold significance in the SN and exceeding GAD+ID at high-threshold significance in the FPCN and SMN.

#### 4.2.2 Brain activations across Threat Conditioning

Across Threat Conditioning (late>early CS+ contrast), several observations suggested greater threat-based low-threshold activation in GAD+ID vs. GAD-ID. First, although within both groups there were activations in the FPCN and DMN, within GAD+ID, there were additional activations in the SN and SM (Fig. 1). However, there were no significant high-threshold activations within either group across this phase. Second, the CS+>CS- contrast at early Threat Conditioning resulted in a greater number of low-threshold clusters and networks (including the SN) within GAD+ID than within GAD-ID, and several of these clusters in the SM and subcortex reached high-threshold significance (Table 3, Supplemental Materials).

#### 4.2.3 Brain activations across Extinction Learning

Across Extinction Learning (late>early CS+E contrast), strikingly different low-threshold activation patterns were seen within GAD-ID vs. GAD+ID groups. First, across-phase low-threshold activations to the CS+ in the FPCN, DM, and SN were more extensive in GAD-ID than in GAD+ID (Fig. 1, 2, Table 3). Second, across-phase activations in the SN and, especially, the FPCN reached high-threshold significance in GAD-ID, whereas in GAD+ID, there were no significant high-threshold activations (Fig. 2, Table 3). Third, across Extinction Learning, low-threshold activations were greater in GAD-ID than GAD+ID in the FPCN (Fig. 2, Table 3). Fourth, there were extensive low-threshold activations to the late CS+E>CS- contrast in GAD+ID, whereas no late CS+E>CS- activations were seen in GAD-ID (Table 3, Supplemental Materials). This contrasts with greater activations to the CS+E *across* Extinction Learning in GAD-ID. These findings suggest that, across Extinction Learning, GAD-ID, unlike GAD+ID, increased activity of both top-down regulatory (FPCN) and, to a lesser extent, threat (SN) regions to the CS+E whereas, by the end of this phase, these regions were no longer more reactive to the CS+E than to the CS-, i.e., the threat had been extinguished (Table 3, Supplemental Materials). In contrast, at the end of Extinction Learning, GAD+ID remained highly reactive to this contrast (i.e., remained more fearful of CS+ than CS-). Extinction Learning involves a competition between the memory for the previously learned threat and acquisition of new safety learning, thus requiring activation of *both* threat (SN) and regulatory (FPCN) regions as took place across this phase in GAD-ID but not GAD+ID.

#### 4.2.4 Brain activations during early Extinction Recall

##### 4.2.4.1 Early CS+E>early CS+U contrast

At Extinction Recall, unlike across Extinction Learning, for the early CS+E>early CS+U contrast, low-threshold activations were now more extensive within GAD+ID than GAD-ID (Fig. 1, 2, Table 3). This was especially the case in the DMN in which one cluster in the parahippocampus and fusiform gyri reached high-threshold significance within GAD+ID (Fig. 2, Table 3). Moreover, greater low-threshold activations, again especially in the DMN, were seen when comparing CS+ID with CS-ID, although these did not reach high-threshold significance (Table 3). This striking reversal of the group showing the greater activation following a delay replicates our previous findings in individuals with ID versus good-sleeping controls (Seo et al., 2018), in sleep-deprived versus fully rested healthy young adults (Seo et al., 2020), and individuals recently exposed to trauma who developed PTSD versus trauma-exposed individuals who did not (Seo et al., 2022) (reviewed in Pace-Schott et al., 2023). This can be interpreted as a delay in the engagement of the neural substrates of extinction learning in individuals whose sleep is compromised (Pace-Schott et al., 2023). Such a delay is possibly analogous to the immediate extinction deficit seen in rodent studies, which has been attributed to a carry-over of stress responses elicited during threat conditioning (Giustino et al., 2017; Maren, 2014, 2022). This possibility is suggested by significant high-threshold activation in CS+ID to the early CS+E>early CS+U contrast in the parahippocampus, a DMN region strongly associated with memory encoding.

##### 4.2.4.2 Early CS+E>early CS- contrast

For the early CS+E>early CS- contrast at Extinction Recall, a greater extent of low-threshold activations within the GAD+ID vs. within the GAD-ID group also occurred (Fig. 1, 2, Table 3). This can be seen in more extensive low-threshold activations of the SN, SM, and DMN in GAD+ID vs. GAD-ID (Fig 1, Table 3). In addition, only within GAD+ID was there low-threshold activation of the FPCN (Fig 1, Table 3). Although within both groups there were regions reaching high-threshold significance, this was seen in a DMN region (temporal pole) for GAD+ID but in the SMN (pre-SMA) for GAD-ID (Fig 1, Table 3). Interestingly, when the two groups were directly compared, low-threshold activation in specific SN and DMN areas was greater in GAD-ID vs. GAD+ID despite more extensive low-threshold activation within GAD+ID (Table 3). The persistence of low threshold activations in the SN of both groups suggests that some degree of threat memory had been maintained in GAD-ID despite their greater neural activity across Extinction Learning.

##### 4.2.4.3 Early CS+U>early CS- contrast

The pattern seen in the contrasts of CS+E with CS+U and CS- was then strikingly reversed for the early CS+U>early CS- contrast in which GAD-ID now showed extensive low-threshold activation in the SN, DMN, and FPCN (Fig. 1), which reached high-threshold significance in an SN region (dACC), whereas, for this contrast in GAD+ID there was only a low-threshold activation in the thalamus (Fig. 1, 2, Table 3). Moreover, low-threshold activation to this contrast was greater in GAD-ID than GAD+ID in SN, DMN, FPCN, and SM networks (Fig. 1, Table 3), reaching high-threshold significance in a region of the FPCN (right VLPFC) (Fig. 2, Table 3). Activation to the reinforced but un-extinguished CS+ in GAD-ID might be interpreted as an memory-based adaptive response to a threat that remains dangerous (unextinguished) whereas the lack of activation in GAD+ID may represent a delay in initiating new extinction learning to the currently non-reinforced CS+U.

### 4.3 Association of sleep measures with neural activations

Across the entire sample (GAD+ID and GAD-ID combined), the interpretation of positive and negative associations between neural activations and sleep variables requires operational definitions of the neurocognitive processes represented by contrasts. Table 1 lists such putative processes associated with each of the 9 contrasts. Only associations reaching low and/or high-threshold significance are reported. Although sleep associations with activations to contrasts are interpreted in light of these processes, it must be remembered that any such sleep effects are indirect.

#### 4.3.1 Sleep quality (sleep efficiency)

To limit the number of analyses, overall sleep quality was operationalized as sleep efficiency (SE). Both objective and subjective SE were analyzed since, in individuals with insomnia, there are characteristic differences between objective sleep and subjective estimates. Among sleep parameters examined, mean objective (actigraphic) and subjective (diary-based) sleep efficiency (SE) showed the greatest number of low-threshold associations with neural activations among the different contrasts (Table 4, Fig. 3).

##### 4.3.1.1 Sleep efficiency and Threat Conditioning

Across Threat Conditioning (late CS+>early CS+), as objective SE increased, threat learning was associated with lesser low-threshold activation in SN, DMN, SM and VN networks, a decrease that was significant at high threshold in vmPFC, a DMN region (Table 4, Fig. 3). Similarly, as subjective SE increased, threat conditioning was associated with reduced activation in areas of the SN, DMN, FPCN, SM and VN (Table 4). These might be interpreted as better sleep quality buffering against anxious (over)reactivity to mild threats, represented here by an annoying shock.

##### 4.3.1.2 Sleep efficiency and Extinction Learning

Across Extinction Learning (late CS+E>early CS+E), as both objective and subjective SE increased, acquisition of extinction (with a concomitant decrease in threat perception) was associated with increased low-threshold activation in the DMN and FPCN networks, albeit without reaching high-threshold significance (Table 4).

##### 4.3.1.3 Sleep efficiency and Extinction Recall

At early Extinction Recall, as both objective and subjective SE increased, discrimination of an extinguished from an un-extinguished threat (early CS+E>early CS+U) was associated with decreased low threshold activation in the SN, DMN, FPCN and SM networks, with increased objective SE associated with high-threshold decrease in the SMG, an FPCN region (Table 4, Fig. 4). At early Extinction Recall, neither objective nor subjective SE were related to the remaining differential threat response to an extinguished CS+ (early CS+E>early CS-). However, at early Extinction Recall, as both objective and subjective SE increased, the differential response to a conditioned but unextinguished threat (early CS+U>early CS-) was associated with increased low-threshold activation in multiple networks (SN, DMN, FPCN, SM, DAN, VN and subcortex and significantly so at high threshold in SPC, SMG and STG (DAN, FPCN and DMN networks respectively).

##### 4.3.1.4 Sleep efficiency relationships with threat conditioning and extinction

Thus greater sleep quality (SE) was associated with brain activations reflecting lesser acquisition of threat, greater acquisition of extinction, lesser differentiation of an extinguished from unextinguished threat, but increased activation to an unextinguished threat. Greater neural response to the recall of the unextinguished stimulus might suggest the association of greater SE with better memory for a continuing threat. Thus, one might speculate that greater sleep quality diminishes initial response to a threat and enhances learning of the extinction of that threat. The seeming paradox of finding an association of greater SE with increased activations across Extinction Learning but decreased activation to the contrast CS+E > CS+U at Extinction Recall may indicate that better sleep quality favors the ability to extinguish soon after conditioning whereas such activation is delayed in those with poorer sleep quality. In contrast, better sleep quality favors increased responding to a “real” (unextinguished) threat represented by the CS+U>CS- at Extinction Recall. Such a generally adaptive, threat-modulating effect of greater SE is further suggested by significant, high-threshold associations seen with activations of emotion-regulatory areas in the FPCN and DMN.

#### 4.3.2 REM and SWS percent

##### 4.3.2.1 Extinction Learning

Across Extinction Learning, as the conditioned threat was being extinguished, increased REM% was associated with decreased low-threshold activations in SN, DMN, and FPCN networks and significantly so at high-threshold in the left AG, a region of the FPCN.

##### 4.3.2.2 Extinction Recall

During early Extinction Recall, when distinguishing extinguished from unextinguished threat conditioning (early CS+E>early CS+U), increased REM% was associated with increased low-threshold activation in SN, DMN, FPCN, and DAN networks and significantly so at high threshold in bilateral AG (FPCN), right MTL (DMN) and left SPC (DAN). When responding to conditioned threat memory remaining after prior extinction (early CS+E>early CS-), increased REM% continued to be associated with greater low-threshold activation in SN, DMN, FPCN as well as VN networks and significantly so at high threshold in the DMPFC (DMN) and fusiform gyrus (VN). However, REM% was unrelated to neural activations during threat conditioning or when the memory of unextinguished threat (early CS+U>early CS-) was being recalled.

##### 4.3.2.3 REM and SWS percent relationships with threat conditioning and extinction

Thus, since unassociated with threat acquisition and unextinguished threat memory, REM% may specifically support processes related to extinction learning and memory. For example, its association with a decrease in activity across Extinction Learning may represent down-regulation of reactivity as extinction is learned (threat is diminished), while the positive association of REM% with activations during recall of an extinguished stimulus might suggest that a greater prior night’s REM% supports differentiation of extinguished from unextinguished threat (CS+E>CS+U) as well as from safety cues (CS+E>CS-). As with REM%, across Extinction Learning, greater SWS% was associated with decreased low-threshold activation in SN and DMN as well as significantly at high threshold in the fusiform gyrus (VN). Thus, like REM%, SWS% may support down-regulation of reactivity as extinction is learned.

#### 4.3.3 REM and SWS HRV relationships with threat conditioning and extinction

REM and SWS HRV (RMSSD) are indicators of parasympathetic outflow during sleep. It has been hypothesized that the role of the parasympathetic nervous system in maintaining physiological homeostasis extends to the domain of emotional regulation (Porges, 2011; Thayer and Lane, 2000). Therefore, the positive association of parasympathetic activity during sleep with activation of regions of the FPCN and DMN across Extinction Learning might suggest that greater HRV during sleep supports the mobilization of emotional and cognitive control during wakefulness. Similarly, the negative associations of sleep HRV with these same networks at the beginning, end, and across Threat Conditioning (Table 4, Supplemental Materials) may indicate that sleep HRV also supports adaptive withdrawal of top-down regulation when threat is being learned. Our recent findings suggest that parasympathetic outflow during REM sleep supports the consolidation of extinction memory in this same protocol (Yuksel et al., 2024b). In addition, parasympathetic activity (during wake) has been shown to be associated with extinction learning (Jenness et al., 2019; Pappens et al., 2014; Wendt et al., 2015), and enhancing parasympathetic activity via vagus nerve stimulation boosts extinction learning and retention of extinction memory in both rodents (e.g., (Souza et al., 2022)) and humans (Burger et al., 2019; Burger et al., 2016; Szeska et al., 2020).

#### 4.3.4 Self-report sleep-quality measures and threat conditioning and extinction

Retrospective measures of sleep quality and insomnia (PSQI, ISI) showed few associations with primary contrasts. While recalling unextinguished differential threat remaining following 24-h delay (CS+U>CS-), greater ISI was associated with lesser low-threshold activations in the SN, DMN, FPCN and SM networks and significantly so at high threshold in IFC/PMC (FPCN/SM networks). For this same contrast, as noted, SE was positively associated with activations in all networks. Thus, greater ID severity may interfere with the ability to recall or respond to threat stimuli that may continue to be threatening. Greater PSQI was negatively associated with somatomotor activation at the beginning of Extinction Learning (early CS+E>CS-) (Table 4, Supplemental Material), which may indicate that poorer sleep quality interferes with initial brain activations associated with extinguishing threat (acting oppositely to RMSSD).

### 4.4 Limitations

The primary limitation of the current study is its small sample size, which resulted from limited funds and restrictions on research during the pandemic. In addition, there were only 2 male participants, both in the GAD-ID group, therefore results may not generalize to men with GAD. Thus, statistical power is clearly limited and results should be interpreted cautiously and replicated with analyses of fully powered samples. The small sample size also precluded the use of connectivity-based measures such as dynamic functional connectivity (Wen et al., 2022) to define large-scale networks. Although the majority of group differences and correlations were seen at the low threshold, the high threshold results either supported or did not contradict low-threshold findings. For example, high-threshold activations during early Threat Conditioning were seen in GAD+ID but not GAD-ID, high-threshold activations in both salience and frontoparietal control networks across Extinction Learning were seen in GAD-ID but not GAD+ID, delayed activation to the CS+E>CS+U and CS+E>CS- contrasts at Extinction Recall was seen in GAD+ID but not GAD-ID, and renewed activity to a previously unextinguished reinforced stimulus (CS+U>CS-) at Extinction Recall was seen only in GAD-ID. Similarly, in multiple regressions, the most common correlate of high-threshold activations was Objective SE which showed positive correlation with activations that were also seen in GAD-ID but not GAD+ID, especially for the CS+U>CS-contrast at Extinction Recall. Several additional limitations existed. First, the menstrual phase and use of contraceptives, both of which can affect extinction (Hwang et al., 2015), were not standardized. Second, in GAD+ID compared to GAD-ID groups, it is possible that both their differential patterns of neural activation to extinction and their insomnia severity might have resulted from more severe GAD (as suggested by PSWQ and STCSA-S being greater in GAD+ID). In this case, differences in neural activations might not have been directly related to sleep quality but rather to GAD severity. Third, sleep associates of threat conditioning and extinction learning and those of extinction recall were measured on two different nights. This might have introduced additional confounding factors such as PSG-measured sleep quality on the consolidation (third) night having benefitted from the additional acclimation to PSG during the baseline (second) night. Fourth, to date, behavioral (psychophysiological and self-report) measurements of fear conditioning and extinction have not shown the delays in timing of extinction learning in sleep-compromised individuals that are suggested by imaging findings (Bottary et al., 2020; Seo et al., 2018).

### 4.5 Conclusions

Group comparisons between individuals with GAD differing in insomnia severity are strikingly similar to our fMRI findings that compared other more to less sleep-compromised individuals, *viz,* the more compromised group fails to engage brain areas supporting extinction-learning immediately following threat conditioning but does so following a delay (Pace-Schott et al., 2023; Seo et al., 2018; Seo et al., 2022; Seo et al., 2020). Such findings may be related to the compromised group being more susceptible to an immediate extinction deficit reported in rodents (Maren, 2022). Nonetheless, studies that separate threat conditioning from extinction learning by varying durations would be needed to assess whether other deficits in the initiation of extinction are also associated with poor sleep. In individuals with GAD, most of whom had some degree of insomnia symptoms, better sleep quality (SE) and increased parasympathetic outflow during REM and SWS (RMSSD) were associated with increased activation of regions that support top-down emotional regulation (FPCN, DMN), while extinction is learned. However, greater REM% was associated with greater activation of emotion regulatory regions when extinction was recalled, suggesting that REM may specifically support consolidation of extinction learning. More definitive conclusions, however, must await better-powered studies contrasting individuals with GAD and other anxiety-related disorders to healthy controls with high-quality sleep.

## 5. Data Availability Statement

The data underlying this article is available in the NIMH Data Archive (NDA) at https://nda.nih.gov, and can be accessed following instructions at https://nda.nih.gov/get/access-data.html.

## 6. Acknowledgments

The authors would like to thank Karen Gannon, RPSGT for scoring sleep recordings, Mary O’Hara and Larry White, Senior MR Technicians at the MGH Martinos Center, for scanning support. Research was carried out at the Athinoula A. Martinos Center for Biomedical Imaging, Charlestown MA and the Massachusetts General Hospital, Department of Psychiatry, Psychiatric Neuroimaging Division.

## 7. Financial disclosures

This project was supported by NIH/NIMH grant R21MH115279 to E.P-S. The authors report no financial conflicts of interest. From 6/2021 to 6/2022 Praxis Precision Medicines, Inc. provided partial salary support to E.P-S.

## Non-financial disclosure

The authors report no non-financial conflicts of interest.

## Abbreviations

AASM: American Acad. Sleep Med
ACC: anterior cingulate cortex
AG: angular gyrus
AIC: anterior insular cortex
BOLD: blood-oxygen-level-dependent
CDT: cluster-determining threshold
CS: conditioned stimulus
CS-: non-reinforced CS
CS+: reinforced CS
CS+E: reinforced and extinguished CS
CS+U: reinforced, not extinguished CS
dACC: dorsal anterior cingulate cortex
dAI: dorsal anterior insula
DAN: dorsal attention network
DLPFC: dorsolateral prefrontal cortex
DMN: default mode network
DMPFC: dorsomedial prefrontal cortex
DSM: APA Diag.Statistical Manual
ECG: electrocardiogram
EEG: electroencephalogram
EMG: electromyogram
EMSQ: eve.-morning sleep diary
EOG: electrooculogram
FIRST: Ford Insomnia Response
fMRI: functional magnetic resonance
FPCN: frontoparietal control network
FWE: family-wise error
GAD: Generalized Anxiety Disorder
GAD+ID: GAD with moderate to severe ID
GAD-ID: GAD without or with mild ID
HC: healthy controls
HRV: heart-rate variability
ID: Insomnia Disorder
IFC: inferior lateral frontal cortex
IPC: inferior parietal cortex
OSA: obstructive sleep apnea
PCC: posterior cingulate cortex
PFC: prefrontal cortex
PIC: posterior insular cortex
PLMP: periodic limb movement disorder
PMC: pre-motor cortex
pre-SMA: pre-supplementary motor area
PSG: polysomnography
PSQI: Pittsburgh Sleep Quality Index
PSWQ: Penn State Worry Questionnaire
PTSD: Posttraumatic Stress Disorder
rACC: rostral anterior cingulate cortex
REM: Rapid Eye Movement sleep
RMSSD: root mean square of R-R intervals
rPFC: rostral prefrontal cortex
RPSGT: registered polysomnographic technician
rsFC: resting state functional connectivity
RV: research version
S1: primary sensory cortex
S2: secondary sensory cortex
SCID: structured clinical interview for DSM-5
SCIDSLD: Pittsburgh interview for sleep disorders
SCL: skin conductance level
SCR: skin conductance response
SE: sleep efficiency
SFC: superior lateral frontal cortex
SM: somatomotor network
SMA: supplementary motor area
SMG: supramarginal gyrus
SN: salience network
SOL: sleep onset latency
SPC: superior parietal cortex
SPM: statistical parametric mapping
STG: superior temporal gyrus
STICSA: State-Trait Invent. Cog. & Somatic Anxiety
ISI: insomnia severity index
ITG: inferior temporal gyrus
M1: primary motor cortex
mCC: middle anterior cingulate cortex
MFC: middle frontal cortex
MIC: middle insular cortex
MOL: middle occipital lobe
MTG: middle temporal gyrus
NREM: Non rapid eye movement
OCD: Obsessive Compulsive Disorder
OFC: orbitofrontal cortex
SWS: slow wave sleep
TIB: time in bed
TST: total sleep time
V1: primary visual cortex
V2: secondary visual cortex
VLPFC: ventrolateral prefrontal cortex
vmPFC: ventromedial prefrontal cortex
VN: visual network
WASO: wake time after sleep onset

## References

Alvaro, P.K., Roberts, R.M., Harris, J.K., 2013. A Systematic Review Assessing Bidirectionality between Sleep Disturbances, Anxiety, and Depression. Sleep 36, 1059–1068.

APA, 2013a. Diagnostic and Statistical Manual of Mental Disorders (Fifth ed.).

American Psychiatric Publishing, Arlington, VA.

APA, 2013b. Diagnostic and Statistical Manual of Mental Disorders, Fifth Edition.

American Psychiatric Publishing, Arlington, VA.

Baglioni, C., Battagliese, G., Feige, B., Spiegelhalder, K., Nissen, C., Voderholzer, U., Lombardo, C., Riemann, D., 2011a. Insomnia as a predictor of depression: a meta-analytic evaluation of longitudinal epidemiological studies. J. Affect. Disord. 135, 10–19.

Baglioni, C., Nanovska, S., Regen, W., Spiegelhalder, K., Feige, B., Nissen, C., Reynolds, C.F., Riemann, D., 2016. Sleep and mental disorders: A meta-analysis of polysomnographic research. Psychol. Bull. 142, 969–990.

Baglioni, C., Riemann, D., 2012. Is chronic insomnia a precursor to major depression? Epidemiological and biological findings. Current psychiatry reports 14, 511–518.

Baglioni, C., Spiegelhalder, K., Nissen, C., Riemann, D., 2011b. Clinical implications of the causal relationship between insomnia and depression: how individually tailored treatment of sleeping difficulties could prevent the onset of depression. The EPMA journal 2, 287–293.

Bastien, C.H., Vallieres, A., Morin, C.M., 2001. Validation of the Insomnia Severity Index as an outcome measure for insomnia research. Sleep medicine 2, 297–307.

Behar, E., Alcaine, O., Zuellig, A.R., Borkovec, T.D., 2003. Screening for generalized anxiety disorder using the Penn State Worry Questionnaire: a receiver operating characteristic analysis. J. Behav. Ther. Exp. Psychiatry 34, 25–43.

Belanger, L., Morin, C.M., Langlois, F., Ladouceur, R., 2004. Insomnia and generalized anxiety disorder: effects of cognitive behavior therapy for gad on insomnia symptoms. J. Anxiety Disord. 18, 561–571.

Bonnet, M.H., Arand, D.L., 2010. Hyperarousal and insomnia: state of the science. Sleep medicine reviews 14, 9–15.

Bottary, R., Seo, J., Daffre, C., Gazecki, S., Moore, K.N., Kopotiyenko, K., Dominguez, J.P., Gannon, K., Lasko, N.B., Roth, B., Milad, M.R., Pace-Schott, E.F., 2020. Fear extinction memory is negatively associated with REM sleep in insomnia disorder. Sleep 43.

Breslau, N., Roth, T., Rosenthal, L., Andreski, P., 1996. Sleep disturbance and psychiatric disorders: a longitudinal epidemiological study of young adults. Biol. Psychiatry 39, 411–418.

Bressler, S.L., Menon, V., 2010. Large-scale brain networks in cognition: emerging methods and principles. Trends in cognitive sciences 14, 277–290.

Burger, A.M., Van Diest, I., Van der Does, W., Korbee, J.N., Waziri, N., Brosschot, J.F., Verkuil, B., 2019. The effect of transcutaneous vagus nerve stimulation on fear generalization and subsequent fear extinction. Neurobiol. Learn. Mem. 161, 192–201.

Burger, A.M., Verkuil, B., Van Diest, I., Van der Does, W., Thayer, J.F., Brosschot, J.F., 2016. The effects of transcutaneous vagus nerve stimulation on conditioned fear extinction in humans. Neurobiol. Learn. Mem. 132, 49–56.

Buysse, D.J., Reynolds, C.F., 3rd, Monk, T.H., Berman, S.R., Kupfer, D.J., 1989. The Pittsburgh Sleep Quality Index: a new instrument for psychiatric practice and research. Psychiatry Res. 28, 193–213.

Cha, J., Carlson, J.M., Dedora, D.J., Greenberg, T., Proudfit, G.H., Mujica-Parodi, L.R., 2014. Hyper-reactive human ventral tegmental area and aberrant mesocorticolimbic connectivity in overgeneralization of fear in generalized anxiety disorder. J. Neurosci. 34, 5855–5860.

Cooper, S.E., Grillon, C., Lissek, S., 2018. Impaired discriminative fear conditioning during later training trials differentiates generalized anxiety disorder, but not panic disorder, from healthy control participants. Compr. Psychiatry 85, 84–93.

Cox, R.C., Olatunji, B.O., 2016. A systematic review of sleep disturbance in anxiety and related disorders. J. Anxiety Disord. 37, 104–129.

Daffre, C., Oliver, K.I., Nazareno, J.R.S., Mader, T., Seo, J., Dominguez, J.P., Gannon, K., Lasko, N.B., Orr, S.P., Pace-Schott, E.F., 2022. Rapid eye movement sleep parasympathetic activity predicts wake hyperarousal symptoms following a traumatic event. J. Sleep Res., e13685.

Di, X., Biswal, B.B., 2022. A functional MRI pre-processing and quality control protocol based on statistical parametric mapping (SPM) and MATLAB. Front Neuroimaging 1, 1070151.

Diekelmann, S., Born, J., 2010. The memory function of sleep. Nature reviews 11, 114–126.

Dodds, K.L., Miller, C.B., Kyle, S.D., Marshall, N.S., Gordon, C.J., 2017. Heart rate variability in insomnia patients: A critical review of the literature. Sleep medicine reviews 33, 88–100.

Dodhia, S., Hosanagar, A., Fitzgerald, D.A., Labuschagne, I., Wood, A.G., Nathan, P.J., Phan, K.L., 2014. Modulation of resting-state amygdala-frontal functional connectivity by oxytocin in generalized social anxiety disorder. Neuropsychopharmacology 39, 2061–2069.

Drake, C., Richardson, G., Roehrs, T., Scofield, H., Roth, T., 2004. Vulnerability to stress-related sleep disturbance and hyperarousal. Sleep 27, 285–291.

Eklund, A., Nichols, T.E., Knutsson, H., 2016. Cluster failure: Why fMRI inferences for spatial extent have inflated false-positive rates. Proc. Natl. Acad. Sci. U. S. A. 113, 7900–7905.

Etkin, A., Prater, K.E., Schatzberg, A.F., Menon, V., Greicius, M.D., 2009. Disrupted amygdalar subregion functional connectivity and evidence of a compensatory network in generalized anxiety disorder. Arch. Gen. Psychiatry 66, 1361–1372.

First, M.B., Williams, J.B.W., Karg, R.S., Spitzer, R.L., 2015. Structured clinical interview for DSM-5—Research version (SCID-5 for DSM-5, research version; SCID-5-RV). American Psychiatric Association, Arlington, VA.

Fonzo, G.A., Etkin, A., 2017. Affective neuroimaging in generalized anxiety disorder: an integrated review. Dialogues in clinical neuroscience 19, 169–179.

Ford, D.E., Cooper-Patrick, L., 2001. Sleep disturbances and mood disorders: an epidemiologic perspective. Depress. Anxiety 14, 3–6.

Ford, D.E., Kamerow, D.B., 1989. Epidemiologic study of sleep disturbances and psychiatric disorders. An opportunity for prevention? JAMA 262, 1479–1484.

Fullana, M.A., Albajes-Eizagirre, A., Soriano-Mas, C., Vervliet, B., Cardoner, N., Benet, O., Radua, J., Harrison, B.J., 2018. Fear extinction in the human brain: A meta-analysis of fMRI studies in healthy participants. Neurosci. Biobehav. Rev. 88, 16–25.

Fullana, M.A., Harrison, B.J., Soriano-Mas, C., Vervliet, B., Cardoner, N., Avila-Parcet, A., Radua, J., 2016. Neural signatures of human fear conditioning: an updated and extended meta-analysis of fMRI studies. Mol. Psychiatry 21, 500–508.

Giustino, T.F., Seemann, J.R., Acca, G.M., Goode, T.D., Fitzgerald, P.J., Maren, S., 2017. beta-Adrenoceptor Blockade in the Basolateral Amygdala, But Not the Medial Prefrontal Cortex, Rescues the Immediate Extinction Deficit. Neuropsychopharmacology.

Glidewell, R.N., McPherson Botts, E., Orr, W.C., 2015. Insomnia and Anxiety: Diagnostic and Management Implications of Complex Interactions. Sleep Med Clin 10, 93–99.

Graham, B.M., Milad, M.R., 2011. The study of fear extinction: implications for anxiety disorders. Am. J. Psychiatry 168, 1255–1265.

Gros, D.F., Antony, M.M., Simms, L.J., McCabe, R.E., 2007. Psychometric properties of the State-Trait Inventory for Cognitive and Somatic Anxiety (STICSA): comparison to the State-Trait Anxiety Inventory (STAI). Psychol Assess 19, 369–381.

Hertenstein, E., Feige, B., Gmeiner, T., Kienzler, C., Spiegelhalder, K., Johann, A., Jansson-Frojmark, M., Palagini, L., Rucker, G., Riemann, D., Baglioni, C., 2018. Insomnia as a predictor of mental disorders: A systematic review and meta-analysis. Sleep medicine reviews 43, 96–105.

Hilbert, K., Lueken, U., Beesdo-Baum, K., 2014. Neural structures, functioning and connectivity in Generalized Anxiety Disorder and interaction with neuroendocrine systems: a systematic review. J. Affect. Disord. 158, 114–126.

Huang, Z., Liang, P., Jia, X., Zhan, S., Li, N., Ding, Y., Lu, J., Wang, Y., Li, K., 2012. Abnormal amygdala connectivity in patients with primary insomnia: evidence from resting state fMRI. Eur. J. Radiol. 81, 1288–1295.

Hunt, C., Park, J., Bomyea, J., Colvonen, P.J., 2023. Sleep efficiency predicts improvements in fear extinction and PTSD symptoms during prolonged exposure for veterans with comorbid insomnia. Psychiatry Res. 324, 115216.

Hwang, M.J., Zsido, R.G., Song, H., Pace-Schott, E.F., Miller, K.K., Lebron-Milad, K., Marin, M.-F., Milad, M.R., 2015. Contribution of estradiol levels and hormonal contraceptives to sex differences within the fear network during fear conditioning and extinction. BMC psychiatry 15, 295.

Iber, C., al., e., 2007. The AASM Manual for the Scoring of Sleep and Associated Events: Rules, Terminology and Technical Specification. American Academy of Sleep Medicine, Westchester, IL.

Insana, S.P., Hall, M., Buysse, D.J., Germain, A., 2013. Validation of the Pittsburgh Sleep Quality Index Addendum for posttraumatic stress disorder (PSQI-A) in U.S. male military veterans. J. Trauma. Stress 26, 192–200.

Jansson-Frojmark, M., Lindblom, K., 2008. A bidirectional relationship between anxiety and depression, and insomnia? A prospective study in the general population. J. Psychosom. Res. 64, 443–449.

Jenness, J.L., Miller, A.B., Rosen, M.L., McLaughlin, K.A., 2019. Extinction Learning as a Potential Mechanism Linking High Vagal Tone with Lower PTSD Symptoms among Abused Youth. J. Abnorm. Child Psychol. 47, 659–670.

Johns, M.W., 1991. A new method for measuring daytime sleepiness: the Epworth sleepiness scale. Sleep 14, 540–545.

Johnson, E.O., Roth, T., Breslau, N., 2006. The association of insomnia with anxiety disorders and depression: exploration of the direction of risk. J. Psychiatr. Res. 40, 700–708.

Kay, D.B., Buysse, D.J., 2017. Hyperarousal and Beyond: New Insights to the Pathophysiology of Insomnia Disorder through Functional Neuroimaging Studies. Brain sciences 7.

Kessler, R.C., Chiu, W.T., Demler, O., Merikangas, K.R., Walters, E.E., 2005. Prevalence, severity, and comorbidity of 12-month DSM-IV disorders in the National Comorbidity Survey Replication. Arch. Gen. Psychiatry 62, 617–627.

Kessler, R.C., Ruscio, A.M., Shear, K., Wittchen, H.U., 2010. Epidemiology of anxiety disorders. Current topics in behavioral neurosciences 2, 21–35.

Kolesar, T.A., Bilevicius, E., Wilson, A.D., Kornelsen, J., 2019. Systematic review and meta-analyses of neural structural and functional differences in generalized anxiety disorder and healthy controls using magnetic resonance imaging. NeuroImage. Clinical 24, 102016.

Kong, R.Q., Spreng, R.N., Xue, A., Betzel, R., Cohen, J.R., Damoiseaux, J., De Brigard, F., Eickhoff, S.B., Fornito, A., Gratton, C., Gordon, E.M., Holmes, A.J., Laird, A.R., Larson-Prior, L., Nickerson, L.D., Pinho, A.L., Razi, A., Sadaghiani, S., Shine, J., Yendiki, A., Yeo, B.T.T., Uddin, L.Q., 2024. A network correspondence toolbox for quantitative evaluation of novel neuroimaging results. bioRxiv.

Krystal, A.D., Stein, M.B., Szabo, S.T., 2017. Anxiety Disorders and Posttraumatic Stress Disorder, in: Kryger, M.H., Roth, T., Dement, W.C. (Eds.), Principles and Practice of Sleep Medicine 6th Edition. Elsevier Health Sciences, Philadelphia, pp. 1341–1351.

Lei, Y., Shao, Y., Wang, L., Ye, E., Jin, X., Zou, F., Zhai, T., Li, W., Yang, Z., 2015. Altered superficial amygdala-cortical functional link in resting state after 36 hours of total sleep deprivation. J. Neurosci. Res. 93, 1795–1803.

Li, J., Zhong, Y., Ma, Z., Wu, Y., Pang, M., Wang, C., Liu, N., Wang, C., Zhang, N., 2020. Emotion reactivity-related brain network analysis in generalized anxiety disorder: a task fMRI study. BMC psychiatry 20, 429.

Li, R., Shen, F., Sun, X., Zou, T., Li, L., Wang, X., Deng, C., Duan, X., He, Z., Yang, M., Li, Z., Chen, H., 2023a. Dissociable salience and default mode network modulation in generalized anxiety disorder: a connectome-wide association study. Cereb. Cortex 33, 6354–6365.

Li, W., Cui, H., Li, H., Colcombe, S., Smith, R.C., Cao, X., Pang, J., Hu, Q., Zhang, L., Yang, Z., Wang, J., Li, C., 2023b. Specific and common functional connectivity deficits in drug-free generalized anxiety disorder and panic disorder: A data-driven analysis. Psychiatry Res. 319, 114971.

Li, W., Cui, H., Zhu, Z., Kong, L., Guo, Q., Zhu, Y., Hu, Q., Zhang, L., Li, H., Li, Q., Jiang, J., Meyers, J., Li, J., Wang, J., Yang, Z., Li, C., 2016. Aberrant Functional Connectivity between the Amygdala and the Temporal Pole in Drug-Free Generalized Anxiety Disorder. Frontiers in human neuroscience 10, 549.

Liu, W.J., Yin, D.Z., Cheng, W.H., Fan, M.X., You, M.N., Men, W.W., Zang, L.L., Shi, D.H., Zhang, F., 2015. Abnormal functional connectivity of the amygdala-based network in resting-state FMRI in adolescents with generalized anxiety disorder. Medical science monitor : international medical journal of experimental and clinical research 21, 459–467.

Ma, Z., Wang, C., Hines, C.S., Lu, X., Wu, Y., Xu, H., Li, J., Wang, Q., Pang, M., Zhong, Y., Zhang, N., 2019. Frontoparietal network abnormalities of gray matter volume and functional connectivity in patients with generalized anxiety disorder. Psychiatry Res Neuroimaging 286, 24–30.

Maren, S., 2014. Nature and causes of the immediate extinction deficit: a brief review. Neurobiol. Learn. Mem. 113, 19–24.

Maren, S., 2022. Unrelenting Fear Under Stress: Neural Circuits and Mechanisms for the Immediate Extinction Deficit. Frontiers in systems neuroscience 16, 888461.

Mellman, T.A., 2008. Sleep and Anxiety Disorders. Sleep Medicine Clinics 3, 261–268.

Milad, M.R., Furtak, S.C., Greenberg, J.L., Keshaviah, A., Im, J.J., Falkenstein, M.J., Jenike, M., Rauch, S.L., Wilhelm, S., 2013. Deficits in conditioned fear extinction in obsessive-compulsive disorder and neurobiological changes in the fear circuit. JAMA Psychiatry 70, 608–618; quiz 554.

Milad, M.R., Quirk, G.J., 2012. Fear extinction as a model for translational neuroscience: ten years of progress. Annu. Rev. Psychol. 63, 129–151.

Milad, M.R., Wright, C.I., Orr, S.P., Pitman, R.K., Quirk, G.J., Rauch, S.L., 2007. Recall of fear extinction in humans activates the ventromedial prefrontal cortex and hippocampus in concert. Biol. Psychiatry 62, 446–454.

Mochcovitch, M.D., da Rocha Freire, R.C., Garcia, R.F., Nardi, A.E., 2014. A systematic review of fMRI studies in generalized anxiety disorder: evaluating its neural and cognitive basis. J. Affect. Disord. 167, 336–342.

Monti, J.M., Monti, D., 2000. Sleep disturbance in generalized anxiety disorder and its treatment. Sleep medicine reviews 4, 263–276.

Motomura, Y., Katsunuma, R., Yoshimura, M., Mishima, K., 2017. Two Days’ Sleep Debt Causes Mood Decline During Resting State Via Diminished Amygdala-Prefrontal Connectivity. Sleep 40.

Motomura, Y., Kitamura, S., Oba, K., Terasawa, Y., Enomoto, M., Katayose, Y., Hida, A., Moriguchi, Y., Higuchi, S., Mishima, K., 2013. Sleep Debt Elicits Negative Emotional Reaction through Diminished Amygdala-Anterior Cingulate Functional Connectivity. PLoS ONE 8, e56578.

Neckelmann, D., Mykletun, A., Dahl, A.A., 2007. Chronic insomnia as a risk factor for developing anxiety and depression. Sleep 30, 873–880.

Noble, S., Scheinost, D., Constable, R.T., 2020. Cluster failure or power failure? Evaluating sensitivity in cluster-level inference. Neuroimage 209, 116468.

Oathes, D.J., Patenaude, B., Schatzberg, A.F., Etkin, A., 2015. Neurobiological signatures of anxiety and depression in resting-state functional magnetic resonance imaging. Biol. Psychiatry 77, 385–393.

Ohayon, M.M., Carskadon, M.A., Guilleminault, C., Vitiello, M.V., 2004. Meta-analysis of quantitative sleep parameters from childhood to old age in healthy individuals: developing normative sleep values across the human lifespan. Sleep 27, 1255–1273.

Ohayon, M.M., Caulet, M., Lemoine, P., 1998. Comorbidity of mental and insomnia disorders in the general population. Compr. Psychiatry 39, 185–197.

Ohayon, M.M., Roth, T., 2003. Place of chronic insomnia in the course of depressive and anxiety disorders. J. Psychiatr. Res. 37, 9–15.

Ou, C.H., Cheng, C.S., Lin, P.L., Lee, C.L., 2024. Grey matter alterations in generalized anxiety disorder: A voxel-wise meta-analysis of voxel-based morphometry studies. Int. J. Dev. Neurosci. 84, 281–292.

Pace-Schott, E.F., Germain, A., Milad, M.R., 2015. Sleep and REM sleep disturbance in the pathophysiology of PTSD: the role of extinction memory. Biology of mood & anxiety disorders 5, 3.

Pace-Schott, E.F., Kaji, J., Stickgold, R., Hobson, J.A., 1994. Nightcap measurement of sleep quality in self-described good and poor sleepers. Sleep 17, 688–692.

Pace-Schott, E.F., Seo, J., Bottary, R., 2023. The influence of sleep on fear extinction in trauma-related disorders. Neurobiology of stress 22, 100500.

Pace-Schott, E.F., Zimmerman, J.P., Bottary, R.M., Lee, E.G., Milad, M.R., Camprodon, J.A., 2017. Resting state functional connectivity in primary insomnia, generalized anxiety disorder and controls. Psychiatry Res. 265, 26–34.

Pappens, M., Schroijen, M., Sutterlin, S., Smets, E., Van den Bergh, O., Thayer, J.F., Van Diest, I., 2014. Resting heart rate variability predicts safety learning and fear extinction in an interoceptive fear conditioning paradigm. PLoS ONE 9, e105054.

Perogamvros, L., Castelnovo, A., Samson, D., Dang-Vu, T.T., 2020. Failure of fear extinction in insomnia: An evolutionary perspective. Sleep medicine reviews 51, 101277.

Pessoa, L., 2018. Understanding emotion with brain networks. Curr Opin Behav Sci 19, 19–25.

Pessoa, L., McMenamin, B., 2017. Dynamic Networks in the Emotional Brain. Neuroscientist 23, 383–396.

Peterson, A., Thome, J., Frewen, P., Lanius, R.A., 2014. Resting-state neuroimaging studies: a new way of identifying differences and similarities among the anxiety disorders? Can. J. Psychiatry. 59, 294–300.

Pizzagalli, D.A., 2011. Frontocingulate dysfunction in depression: toward biomarkers of treatment response. Neuropsychopharmacology 36, 183–206.

Porges, S.W., 2011. The polyvagal theory. Neurophysiological Foundations of Emotions, Attachment, Communication, and Self-regulation. Norton, New York.

Qiao, J., Li, A., Cao, C., Wang, Z., Sun, J., Xu, G., 2017. Aberrant Functional Network Connectivity as a Biomarker of Generalized Anxiety Disorder. Frontiers in human neuroscience 11, 626.

Rasch, B., Born, J., 2013. About sleep’s role in memory. Physiol. Rev. 93, 681–766.

Ree, M.J., French, D., MacLeod, C., Locke, V., 2008. Distinguishing Cognitive and Somatic Dimensions of State and Trait Anxiety: Development and Validation of the State-Trait Inventory for Cognitive and Somatic Anxiety (STICSA). Behavioural and Cognitive Psychotherapy 36, 313–332.

Ree, M.J., MacLeod, C., French, D., Locke, V., 2000. State–Trait Inventory for Cognitive and Somatic Anxiety (STICSA)—Trait Version. The University of Western Australia, Perth, Australia.

Reiss, S., Peterson, R.A., Gursky, D.M., McNally, R.J., 1986. Anxiety sensitivity, anxiety frequency and the prediction of fearfulness. Behav. Res. Ther. 24, 1–8.

Riemann, D., 2007. Insomnia and comorbid psychiatric disorders. Sleep medicine 8 Suppl 4, S15–20.

Riemann, D., Spiegelhalder, K., Feige, B., Voderholzer, U., Berger, M., Perlis, M., Nissen, C., 2010. The hyperarousal model of insomnia: a review of the concept and its evidence. Sleep medicine reviews 14, 19–31.

Roy, A.K., Fudge, J.L., Kelly, C., Perry, J.S., Daniele, T., Carlisi, C., Benson, B., Castellanos, F.X., Milham, M.P., Pine, D.S., Ernst, M., 2013. Intrinsic functional connectivity of amygdala-based networks in adolescent generalized anxiety disorder. J. Am. Acad. Child Adolesc. Psychiatry 52, 290–299 e292.

Rush, A.J., Trivedi, M.H., Ibrahim, H.M., Carmody, T.J., Arnow, B., Klein, D.N., Markowitz, J.C., Ninan, P.T., Kornstein, S., Manber, R., Thase, M.E., Kocsis, J.H., Keller, M.B., 2003. The 16-Item Quick Inventory of Depressive Symptomatology (QIDS), clinician rating (QIDS-C), and self-report (QIDS-SR): a psychometric evaluation in patients with chronic major depression. Biol. Psychiatry 54, 573–583.

Seo, J., Moore, K.N., Gazecki, S., Bottary, R.M., Milad, M.R., Song, H., Pace-Schott, E.F., 2018. Delayed fear extinction in individuals with insomnia disorder. Sleep 41.

Seo, J., Oliver, K.I., Daffre, C., Moore, K.N., Gazecki, S., Lasko, N.B., Milad, M.R., Pace-Schott, E.F., 2022. Associations of sleep measures with neural activations accompanying fear conditioning and extinction learning and memory in trauma-exposed individuals. Sleep 45.

Seo, J., Pace-Schott, E.F., Milad, M.R., Song, H., Germain, A., 2020. Partial and Total Sleep Deprivation Interferes With Neural Correlates of Consolidation of Fear Extinction Memory. Biol Psychiatry Cogn Neurosci Neuroimaging.

Shao, Y., Lei, Y., Wang, L., Zhai, T., Jin, X., Ni, W., Yang, Y., Tan, S., Wen, B., Ye, E., Yang, Z., 2014. Altered resting-state amygdala functional connectivity after 36 hours of total sleep deprivation. PLoS ONE 9, e112222.

Souza, R.R., Powers, M.B., Rennaker, R.L., McIntyre, C.K., Hays, S.A., Kilgard, M.P., 2022. Timing of vagus nerve stimulation during fear extinction determines efficacy in a rat model of PTSD. Scientific reports 12, 16526.

Spitzer, R.L., Kroenke, K., Williams, J.B., Lowe, B., 2006. A brief measure for assessing generalized anxiety disorder: the GAD-7. Arch. Intern. Med. 166, 1092–1097.

Stein, M.B., Sareen, J., 2015. CLINICAL PRACTICE. Generalized Anxiety Disorder. N. Engl. J. Med. 373, 2059–2068.

Stocker, R.P.J., Khan, H., Henry, L., Germain, A., 2017. Effects of Sleep Loss on Subjective Complaints and Objective Neurocognitive Performance as Measured by the Immediate Post-Concussion Assessment and Cognitive Testing. Arch Clin Neuropsychol 32, 349–368.

Sun, J., Zhang, B., Xu, W., Li, P., Zhang, D., Zhao, B., Wang, Z., Wang, B., 2024. Effectiveness of repetitive transcranial magnetic stimulation for insomnia disorder on fear memory extinction: study protocol for a randomised controlled trial. Trials 25, 396.

Szeska, C., Richter, J., Wendt, J., Weymar, M., Hamm, A.O., 2020. Promoting long-term inhibition of human fear responses by non-invasive transcutaneous vagus nerve stimulation during extinction training. Scientific reports 10, 1529.

Thayer, J.F., Lane, R.D., 2000. A model of neurovisceral integration in emotion regulation and dysregulation. J. Affect. Disord. 61, 201–216.

Tsypes, A., Aldao, A., Mennin, D.S., 2013. Emotion dysregulation and sleep difficulties in generalized anxiety disorder. J. Anxiety Disord. 27, 197–203.

Uddin, L.Q., 2023. A Brain Network by Any Other Name. J. Cogn. Neurosci. 35, 363–364.

Uddin, L.Q., Betzel, R.F., Cohen, J.R., Damoiseaux, J.S., De Brigard, F., Eickhoff, S.B., Fornito, A., Gratton, C., Gordon, E.M., Laird, A.R., Larson-Prior, L., McIntosh, A.R., Nickerson, L.D., Pessoa, L., Pinho, A.L., Poldrack, R.A., Razi, A., Sadaghiani, S., Shine, J.M., Yendiki, A., Yeo, B.T.T., Spreng, R.N., 2023. Controversies and progress on standardization of large-scale brain network nomenclature. Netw Neurosci 7, 864–905.

Uddin, L.Q., Yeo, B.T.T., Spreng, R.N., 2019. Towards a Universal Taxonomy of Macro-scale Functional Human Brain Networks. Brain Topogr. 32, 926–942.

Wang, H., Mou, S., Pei, X., Zhang, X., Shen, S., Zhang, J., Shen, X., Shen, Z., 2025. The power spectrum and functional connectivity characteristics of resting-state EEG in patients with generalized anxiety disorder. Scientific reports 15, 5991.

Wang, M., Cao, L., Li, H., Xiao, H., Ma, Y., Liu, S., Zhu, H., Yuan, M., Qiu, C., Huang, X., 2021. Dysfunction of Resting-State Functional Connectivity of Amygdala Subregions in Drug-Naive Patients With Generalized Anxiety Disorder. Frontiers in psychiatry 12, 758978.

Wang, T., Yan, J., Li, S., Zhan, W., Ma, X., Xia, L., Li, M., Lin, C., Tian, J., Li, C., Jiang, G., 2017. Increased insular connectivity with emotional regions in primary insomnia patients: a resting-state fMRI study. Eur. Radiol.

Wang, W., Hou, J., Qian, S., Liu, K., Li, B., Li, M., Peng, Z., Xin, K., Sun, G., 2016. Aberrant regional neural fluctuations and functional connectivity in generalized anxiety disorder revealed by resting-state functional magnetic resonance imaging. Neurosci. Lett. 624, 78–84.

Wang, Z., Luo, Y., Zhang, Y., Chen, L., Zou, Y., Xiao, J., Min, W., Yuan, C., Ye, Y., Li, M., Tu, M., Hu, J., Zou, Z., 2023. Heart rate variability in generalized anxiety disorder, major depressive disorder and panic disorder: A network meta-analysis and systematic review. J. Affect. Disord. 330, 259–266.

Wen, Z., Pace-Schott, E.F., Lazar, S.W., Rosen, J., Ahs, F., Phelps, E.A., LeDoux, J.E., Milad, M.R., 2024. Distributed neural representations of conditioned threat in the human brain. Nature communications 15, 2231.

Wen, Z., Seo, J., Pace-Schott, E.F., Milad, M.R., 2022. Abnormal dynamic functional connectivity during fear extinction learning in PTSD and anxiety disorders. Mol. Psychiatry 27, 2216–2224.

Wendt, J., Neubert, J., Koenig, J., Thayer, J.F., Hamm, A.O., 2015. Resting heart rate variability is associated with inhibition of conditioned fear. Psychophysiology 52, 1161–1166.

Westlin, C., Theriault, J.E., Katsumi, Y., Nieto-Castanon, A., Kucyi, A., Ruf, S.F., Brown, S.M., Pavel, M., Erdogmus, D., Brooks, D.H., Quigley, K.S., Whitfield-Gabrieli, S., Barrett, L.F., 2023. Improving the study of brain-behavior relationships by revisiting basic assumptions. Trends in cognitive sciences 27, 246–257.

Xiong, H., Guo, R.J., Shi, H.W., 2020. Altered Default Mode Network and Salience Network Functional Connectivity in Patients with Generalized Anxiety Disorders: An ICA-Based Resting-State fMRI Study. Evidence-based complementary and alternative medicine : eCAM 2020, 4048916.

Yang, F., Zhang, J., Fan, L., Liao, M., Wang, Y., Chen, C., Zhai, T., Zhang, Y., Li, L., Su, L., Dai, Z., 2020. White matter structural network disturbances in first-episode, drug-naive adolescents with generalized anxiety disorder. J. Psychiatr. Res. 130, 394–404.

Yeo, B.T., Krienen, F.M., Sepulcre, J., Sabuncu, M.R., Lashkari, D., Hollinshead, M., Roffman, J.L., Smoller, J.W., Zollei, L., Polimeni, J.R., Fischl, B., Liu, H., Buckner, R.L., 2011. The organization of the human cerebral cortex estimated by intrinsic functional connectivity. J. Neurophysiol. 106, 1125–1165.

Yoo, S.S., Gujar, N., Hu, P., Jolesz, F.A., Walker, M.P., 2007. The human emotional brain without sleep--a prefrontal amygdala disconnect. Curr. Biol. 17, R877–878.

Yuan, M., Liu, B., Yang, B., Dang, W., Xie, H., Lui, S., Qiu, C., Zhu, H., Zhang, W., 2023. Dysfunction of default mode network characterizes generalized anxiety disorder relative to social anxiety disorder and post-traumatic stress disorder. J. Affect. Disord. 334, 35–42.

Yuksel, C., Watford, L., Muranaka, M., Daffre, C., McCoy, E., Lax, H., Mendelsohn, A.K., Oliver, K.I., Acosta, A., Vidrin, A., Martinez, U., Lasko, N., Orr, S., Pace-Schott, E.F., 2024a. REM disruption and REM vagal activity predict extinction recall in trauma-exposed individuals. Psychol. Med. 54, 1–12.

