## Supplemental Materials for "Local and network neural activations and their associations with sleep parameters during threat conditioning and extinction in persons with Generalized Anxiety Disorder with and without Insomnia Disorder"

### S1. Supplemental Methods

#### S1.1 Group cutoff rationale

The cut-off score was chosen based upon a median split of the entire sample (ISI=13.5) before individuals were excluded due to fMRI artifact. Although the standard ISI cutoff on the ISI is between sub-threshold insomnia (maximum 14) and moderate severity insomnia (minimum 15), we wanted the GAD-ID group to be as asymptomatic as possible to enhance the separation between groups as most individuals diagnosed with GAD show some degree of sleep disturbance.

#### S1.2 Exclusion criteria

1. Lifetime history of psychosis, bipolar disorder, autism spectrum or other neurodevelopmental disorder, suicide attempt or chronic suicidal ideation
2. Current major depressive episode or suicidal ideation
3. Lifetime history of a sleep disorder except Insomnia Disorder (however Insomnia Disorder with medical comorbidity excluded)

4. On the PSG acclimation/screening night, no OSA or PLMD of a severity sufficient to warrant referral for treatment.

5. Neurologic conditions including past neurosurgical procedures, seizure, neurodegenerative disease

6. History of significant head injury.

7. Medical conditions that could confound outcome variables such as severe cardiovascular or other systemic disease.

9. Use of specific psychotropic medications such as hypnotics and antipsychotics

10. Current drug or alcohol abuse or dependence or positive urinalysis result for any major class of abused drugs.

11. Expressed unwillingness, when asked, to remain entirely alcohol- and drug-free throughout the study

12, Daily caffeine intake of >4 cups or weekly alcohol >10 drinks

13. MRI contraindications (e.g., metal in body or eyes, pacemaker, pump, stimulator, shunt, claustrophobia, pregnancy, weight >250 lbs.).

15. Supervisees of study investigators

#### S1.3 Procedural flow-chart

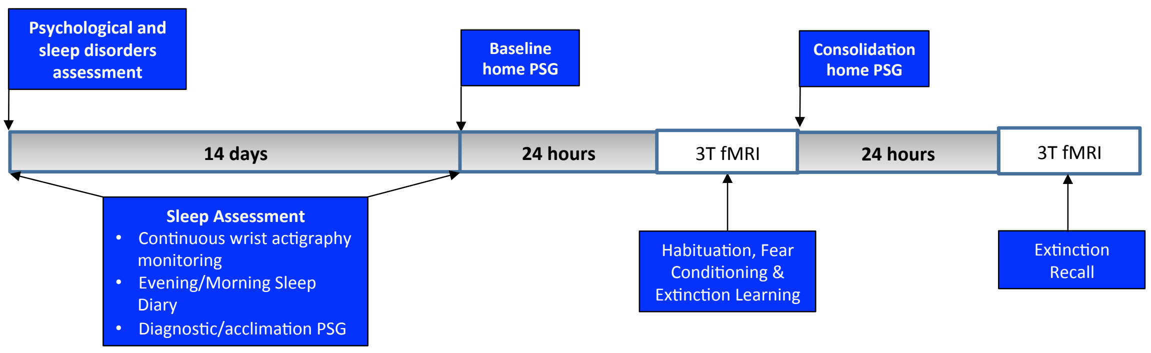

**S1.4 Actigraphy**

Participants wore a wrist actigraph (Actiwatch 2, Philips Respironics, Bend, OR) on the non-dominant wrist 24 hours per day throughout the study. The Actiwatch recorded arm movements in 1-min epochs. Participants pressed an event marker button on the watch at the beginning of each sleep attempt and as soon as they awoke each morning. These paired markings demarcated nightly Actiwatch time in bed, which was used by the default Actiware software algorithm (Actiware 5.61) to determine total sleep time (TST), sleep onset latency (SOL), sleep efficiency (SE), and sleep midpoint (calculated as minutes past midnight). When participants failed to press the event marker at either time point, diary entries for bed and/or wake times from the corresponding day were substituted. Due to probable artifact or equipment malfunction, nights showing SE < 50% and/or SE more than 20% below the corresponding diary SE were excluded from analyses.

**S1.5 Timing of threat conditioning and extinction**

The threat and extinction sessions with fMRI were performed in the evenings in order to: 1) Avoid potential inter-individual differences in circadian effects on SCR(Pace-Schott et al., 2013), 2) to maximize the magnitude of SC responses(Pace-Schott et al., 2013), and 3) to minimize wake time prior sleep following extinction learning given the possibility of a limited post-encoding interval during which sleep-dependent memory can take place.

**S1.6 Polysomnographic methods**

Electrodes were attached in the laboratory during the evening prior to each night of ambulatory PSG. The Somte-PSG recorder was contained in a customized cloth pack, worn on the chest, which encloses all loose wires. It was set to begin recording before the participant's earliest anticipated bedtime. The montage included 6 EEG channels

(F3, F4, C3, C4, O1, O2) referenced to contralateral mastoids (A1, A2), as well as 2 EOG channels (both outer canthi, one above and one below the eye), 2 channels of submental EMG, and 2 channels for ECG (right clavicle, left 5th intercostal space). On the diagnostic night, channels for pulse-oximeter, respiration transducer belts, nasal cannula and tibialis periodic limb movement (PLM) sensors were added to screen for sleep disordered breathing and periodic limb movement disorder. A research-experienced Registered Polysomnographic Technician (RPSGT), blind to participant group, scored sleep following standard AASM criteria(Iber and al., 2007; Rechtschaffen and Kales, 1968) using Compumedics Profusion 4.0 software. Resulting hypnograms were be used to calculate REM%, SWS% and their heart-rate variability (HRV). Electrocardiogram (ECG) data were analyzed for HRV using Kubios HRV Premium software (Kubios Oy, Kuopio, Finland). Measures of parasympathetic activity in all REM and SWS periods exceeding 5 min were computed in the time domain (root-mean square differences of successive R-R intervals, RMSSD).(Laborde et al., 2017; Shaffer and Ginsberg, 2017)

**S1.7 Sleep diary: Evening-Morning Sleep Questionnaire (EMSQ)** (Pace-Schott et al., 1994; Pace-Schott et al., 2005), The evening portion of the EMSQ queried prior daytime activities and the time at which the participant began to attempt sleep. Morning portions queried time waking for the day, subjective SOL, subjective TST, and number and duration of nocturnal awakenings (summed as subjective wake time after sleep onset or "WASO"). Subjective sleep efficiency was computed from diaries as proportion of time in bed (TIB; duration between the time at which subject began to attempt sleep and time of waking for the day) occupied by sleep [TIB – (SOL + WASO)].

**S1.8 Self-report assessments:**

The Generalized Anxiety Disorder 7-item scale (GAD-7) (Spitzer et al., 2006), is a 7-item scale with a GAD threshold of $\geq$ 10. The Pittsburgh Sleep Quality Index (PSQI) (Buysse et al., 1989), is the most widely used standardized sleep quality assessment. The Insomnia Severity Index (ISI)(Bastien et al., 2001), measures insomnia severity with ID threshold of 15 (Bastien et al., 2001). Ford Insomnia Response to Stress Test (FIRST) (Drake et al., 2004) is an assessment of individual vulnerability to acute-stress induced insomnia. The Morningness-Eveningness questionnaire (MEQ) (Horne and Ostberg, 1976), is a standard assessment of chronotype. The Epworth Sleepiness Scale (ESS) (Johns, 1991), is the standard assessment of habitual daytime sleepiness. The Penn State Worry Questionnaire (PSWQ)(Behar et al., 2003), is a standard assessment of worry. Spielberger State-Trait Anxiety Inventory (STAI-T) (Spielberger et al., 1990) , is a standard measure of trait anxiety.(Spielberger et al., 1990). The Inventory of Depressive Symptomatology (IDT) (Rush et al., 2003), measures current depressive symptoms. The Revised NEO Personality Inventory (NEO-PI-R) (Costa and McCrae, 1992), yields scores for “big-five” personality factors. The State-Trait Inventory for Cognitive and Somatic Anxiety-Trait Version (STICSA-T) (Gros et al., 2007; Ree et al., 2008; Ree et al., 2000). The Anxiety Sensitivity Index (ASI) (Reiss et al., 1986), The Multidimensional Assessment of Interoceptive Awareness (MAIA) (Mehling et al., 2012), subjective interoceptive sensitivity.

**S1.9 fMRI data acquisition and preprocessing:** Scanning took place in a 3-Tesla Siemens Magnetom Prisma scanner magnet with a 32-channel head coil. The fMRI protocol will be an event-related design used extensively by our team(Milad et al., 2013; Milad et al., 2009; Milad et al., 2007a; Milad et al., 2007b; Seo et al., 2018; Seo et al., 2022a) with MRI acquisition parameters identical to those previously used. Briefly, after an automated scout image was acquired and shimming procedures performed to optimize field homogeneity, high-resolution 3D MEMPRAGE sequences (TR/TE1,2,3,4/flip angle=2530ms/1.64,3.5,5,36,7.22ms/7°) with an in-plane resolution of 1.0 mm, and 1 mm slice thickness, was collected for spatial normalization, positioning the slice prescription to assist in registration of the functional data to the high-resolution anatomical scan. Functional MRI images were acquired using gradient echo T2*-weighted sequences (TR/TE/Flip angle = 2.56 sec/30 ms/ 90 degrees) consisting of 46 axial oblique slices parallel to the anterior-posterior commissure line with a slice thickness of 3 mm x 3 mm x 3mm. Pre-processing: Data were pre-processed using statistical parametric mapping software (SPM12; http://www.fil.ion.ucl.ac.uk/spm). Data for each participant was first corrected for differences in acquisition time between slices, realigned using the first slice as a reference and unwarped to correct for static inhomogeneity of the magnetic field and movement by inhomogeneity interactions. Realigned functional images were then co-registered with each participant's anatomical image, normalized with the anatomical gray matter parameters to the standard Montreal Neurological Institute template, and spatially smoothed with a Gaussian kernel of 8-mm full-width at half-maximum (FWHM).

**S1.10 Measurements of objective and subjective threat conditioning and extinction**

*Objective measure (skin conductance response, SCR):* As detailed in numerous past publications,(Bottary et al., 2020; Marin et al., 2017; Seo et al., 2018; Seo et al., 2022b; Seo et al., 2020; Yuksel et al., 2024) skin conductance level (SCL) was continuously measured during fMRI scans throughout the threat conditioning and extinction paradigm using a Biopac Systems, Inc. (Goleta, CA) MP150 MRI-compatible system. Sampling was carried out at 200 Hz along with event markers indicating the onset of each context, CS and shock allowing precise synchronization of each stimulus onset with the ongoing

SCL recording. SCR was calculated by subtracting the mean SCL during the last 2 sec of context-alone presentation from the peak SCL during the 6-sec CS presentation and square-root transformed. If the untransformed SCR was negative, the negative sign was retained after calculating the square root of its absolute value.(Orr et al., 2000) Excluded non-conditioning subjects showed ≦ 1 non-transformed SCR to a CS+ that exceeded 0.05 μS(Orr et al., 1995) during Fear Conditioning.

*The subjective measure was shock expectancy*. Immediately following each phase, participants verbally rated their expectancy of being shocked by each CS color at its first and last presentation on a 5-point scale.

**S1.11 Statistical analysis confirming successful objective and subjective threat conditioning and extinction learning.**

To test whether physiological fear conditioning and extinction learning were achieved, one-way repeated measures ANOVAs were used to compare SCRs between CS+ and CS- at early and late Conditioning and Extinction Recall subphases. For Conditioning, the early subphase consisted of trials 2-4, while it included the first 4 trials for Extinction Learning. The first trial for Conditioning was excluded because fear learning had not yet occurred. The late subphase included the last 4 trials for both Conditioning and Extinction Learning phases. To test whether explicit fear conditioning and extinction learning were achieved, two-way repeated measures ANOVAs with CS type (CS+ and CS-) and order (first and last) as between-subject factors, were used to compare the change in subjective ratings between CS+ and CS- at Conditioning and Extinction Learning. Statistical analyses were performed using IBM SPSS Statistics Version 26.

### S2. Supplemental Results

#### S2.1 Confirmation of successful objective and subjective threat conditioning and extinction learning.

Analysis of both SCR and subjective ratings showed that fear conditioning and extinction were achieved. For SCR, there was a significantly higher response to CS+ compared to CS- in both early ($F(1,19)=11.999$, $p=0.003$, $\eta_p^2 = 0.387$) and late ($F(1,19)=13.840$, $p=0.001$, $\eta_p^2 = 0.421$) Conditioning. There was no difference in SCR between CS types (CS+ vs. CS-) at early ($F(1,19)=0.114$, $p=0.739$, $\eta_p^2 = 0.006$) or late ($F(1,19)=0.214$, $p=0.649$, $\eta_p^2 = 0.011$) Extinction Learning. For subjective ratings, there was a significant CS type (CS+ vs. CS-) × Order (first vs. last) interaction at both Conditioning ($F(1,27)=42.226$, $p<0.001$, $\eta_p^2 = 0.619$) and Extinction Learning ($F(1,27)= 7.619$, $p=0.010$, $\eta_p^2 = 0.227$).

#### S2.2 Within group brain activation results

*S2.2.1 Early Threat Conditioning (early CS+ > early CS- contrast)*

At early Threat Conditioning (early CS+ > early CS- contrast), in GAD+ID, there was increased low-threshold activation in regions of the somatomotor (PMC, S1; 255 SM voxels), frontoparietal control (IPC, AG: 63 FPCN voxels), salience (rACC, dAI; 76 SN voxels) as well as subcortical areas (pallidum, putamen, caudate, thalamus; 309 voxels) (Table 3). In GAD+ID, clusters reached significant high-threshold activation for this contrast in the somatomotor (left PMC and S1) network and subcortically in left Pallidum and Putamen (Table 3). In GAD-ID, there was increased low-threshold activation in regions of the dorsal attention (SPC; 50 DAN voxels), somatomotor (SMA, PMC; 129 SM voxels), default (STP; 19 DMN voxels) and subcortical areas (pallidum; 41 voxels) (Table 3). No clusters reached high-threshold significance for this contrast in In GAD-ID (Table 2.).

*S2.2.2 Late Threat Conditioning (late CS+ > late CS- contrast)*

At late Threat Conditioning (late CS+ > late CS- contrast), in GAD+ID, there was increased low-threshold activation in regions of the salience (dorsal anterior insula, middle insula; 71 SN voxels), default (DMPFC; 49 DMN voxels) and visual (V2; 176 VN voxels) networks (Table 3). However, no clusters reached high threshold significance (Table 3). In GAD-ID, there was increased low-threshold activation in regions of the default (precuneus; 38 DMN voxels), somatomotor (SMA; 19 SM) and visual (V2; 33 VN voxels), however, as in GAD+ID, no clusters reached high-threshold significance (Table 3). GAD+ID showed greater low-threshold activation than GAD-ID in the default (DMPFC, hippocampus, temporal pole; 211 DMN voxels) and frontoparietal control network (DLPFC, IFC; 96 FPCN voxels), while GAD-ID showed greater low-threshold activation than GAD+ID in salience (dACC; 58 SN voxels) and somatomotor (M1, S1; 113 SM voxels) however, neither group difference contained clusters that reached high-threshold significance (Table 3).

*S2.2.3 Early Extinction Learning (early CS+E > early CS-)*

At early Extinction Learning (early CS+E > early CS-) in GAD+ID, there was increased low-threshold activation in regions of the default (parahippocampus; 28 DMN voxels), frontoparietal control (DLPFC; 19 FPCN voxels) and visual (fusiform; 92 VN voxels) networks, no clusters in which reached high-threshold significance (Table 3). In GAD-ID, there was increased low-threshold activation in regions of the frontoparietal control (DLPFC, SMG; 263 FPCN voxels) and visual (MOL; 341 VN voxels) networks. GAD+ID showed greater low-threshold activation than GAD-ID in subcortex (putamen; 98 voxels), while GAD-ID showed greater low-threshold activation than GAD+ID in frontoparietal control network (IPC, SMG, SFC; 241 FPCN voxels), however, neither group difference contained clusters that reached high-threshold significance (Table3)

*S2.2.4 Late Extinction Learning (late CS+E > late CS-)*

At late Extinction Learning (late CS+E > late CS-) in GAD+ID, there was increased low-threshold activation in regions of the salience (dACC, rACC, PIC; 203 SN voxels), default (DMPFC, PCC/precuneus, hippocampus, 334 DMN voxels), frontoparietal control (ITG; 226 FPCN voxels) networks and subcortically in the putamen (32 voxels). no clusters in which reached high-threshold significance (Table 2.). In GAD+ID, a cluster in the right ITG (FPCN network) reached significant high-threshold activation. In contrast, at late Extinction Learning for this contrast, GAD-ID showed no low-threshold activations. GAD+ID showed greater low-threshold activation than GAD-ID in salience (rACC, MIC; 89 SN voxels), default (PCC, DMPFC, hippocampus, SFC; 156 DMN voxels) and frontoparietal control (STG; 36 FPCN voxels) networks, however no clusters showed high-threshold significance for these group differences.

**S2.3 Association of sleep measures with neural activations**

Note: Only associations reaching low and/or high-threshold significance are reported.

*S2.3.1 Objective and subjective sleep efficiency (SE)*

*S2.3.1.1 SE and early threat conditioning* (early CS+ > early CS- contrast)

During Fear Conditioning, for the early CS+ > early CS- contrast, as **objective SE** increased, low threshold activation increased in somatomotor (S1, premotor cortex, 217 SM voxels), salience (mCC, IFC, 54 SN voxels), and default (OFC, vmPFC, 300 DMN voxels) networks. Increased activation with increased objective SE reached high threshold significance in the vmPFC (Table 4). As **subjective SE** increased, low threshold activation increased in the default (vmPFC; 26 DMN voxels) and somatomotor (PMC; 49 SM voxels) networks but decreased in the salience (dACC, rACC; 152 SN voxels), default mode (hippocampus; 40 DMN voxels) and somatomotor (SMA; 55 SM voxels) networks. However, no activations were associated with subjective SE at high-threshold significance (Table 4).

*S2.3.1.2 SE and late threat conditioning* (late CS+ > late CS- contrast)

During Fear Conditioning, for the late CS+ > late CS- contrast, as **objective SE** increased, low-threshold activation increased in salience (STG; 33 SN voxels), default (precuneus, MTG, parahippocampus; 269 DMN voxels), frontoparietal control (SMG; 271 FPCN voxels) and somatomotor (PMC; 278 SM voxels) networks but decreased in the salience network (AIC; 14 SN voxels). Increased activation with increased objective SE reached high threshold significance in the bilateral SMG (Table 4.). As **subjective SE** increased, low-threshold activation decreased in default (PCC; 105 DMN voxels), frontoparietal control (DLPFC; 34 FPCN voxels) and somatomotor (SMA; 202 SM voxels) networks. Decreased activation with increased subjective SE reached high-threshold significance in the SMA (Table 4).

*S2.3.1.3 SE and early Extinction Learning* (early CS+ > early CS- contrast)

During Extinction Learning, for the early CS + E > early CS- contrast, as **subjective SE** increased, low-threshold activation decreased in the default (parahippocampus; 27 DMN voxels) and visual (middle occipital cortex; 220 VN voxels). Decreased activation with increased subjective SE reached high threshold significance in the left middle occipital cortex)

*S2.3.1.4 SE and late Extinction Learning* (late CS+ > late CS- contrast)

During Extinction Learning, for the late CS + E > late CS- contrast, as **objective SE** increased, low threshold activation increased in the frontoparietal control (AG; 154 FPCN voxels) and somatomotor (PMC; 38 SM voxels), but decreased in the salience (AIC; 79 SN voxels), default (DMPFC, hippocampus; 192 DMN voxels), visual (superior and middle occipital lobe; 1017 VN voxels), somatomotor (M1; 411 SM voxels) networks and subcortex (caudate; 50 voxels). Decreased activation with increased objective SE reached high-threshold significance in the left superior occipital cortex. As **subjective SE** increased, low-threshold activation increased in the somatomotor network (SMA; 25 SMA voxels) but decreased in default (DMPFC, mCC; 232 DMN voxels), frontoparietal control (IPC; 208 FPCN voxels) and visual (superior and middle occipital lobes; 2023 VN voxels). Decreased activation with increased subjective SE reached high-threshold significance in the bilateral superior and middle occipital lobes (Table 4).

*S2.3.2 Heart-rate variability (RMSSD) in REM and SWS*

S2.3.2.1 Early Threat Conditioning

During early Threat Conditioning, for the early CS + E > early CS- contrast, as REM RMSSD increased, low-threshold activation decreased in the salience network (dACC; 15 SN voxels). Similarly, as SWS RMSSD increased, low-threshold activation decreased in the salience (dACC; 52 SN voxels) and default (parahippocampus; 44 DMN voxels) networks.

S2.3.2.2 Late Threat Conditioning

During late Threat Conditioning, for the late CS + E > late CS- contrast, as REM RMSSD increased, low-threshold activation decreased in the salience (AIC; 34 SN voxels) and frotoparietal control (OFC; 11 FPCN voxels) networks. Similarly, as SWS RMSSD increased, there was decreased low-threshold activation in the frotoparietal control network (OFC; 60 FPCN voxels).

*S2.3.3 Pittsburgh Sleep Quality Index (PSQI)*

During early Extinction Learning, for the early CS + E > early CS- contrast as PSQI increased, low-threshold activation decreased in the somatomotor network (PMC, pre-SMA; 302 SM voxels). Decreased activation with increased PSQI reached high-threshold significance in the left PMC (Table 4).

### S.3 References


Bastien, C.H., Vallieres, A., Morin, C.M., 2001. Validation of the Insomnia Severity Index as an outcome measure for insomnia research. Sleep medicine 2, 297-307.

Behar, E., Alcaine, O., Zuellig, A.R., Borkovec, T.D., 2003. Screening for generalized anxiety disorder using the Penn State Worry Questionnaire: a receiver operating characteristic analysis. J. Behav. Ther. Exp. Psychiatry 34, 25-43.

Bottary, R., Seo, J., Daffre, C., Gazecki, S., Moore, K.N., Kopotiyenko, K., Dominguez, J.P., Gannon, K., Lasko, N.B., Roth, B., Milad, M.R., Pace-Schott, E.F., 2020. Fear extinction memory is negatively associated with REM sleep in insomnia disorder. Sleep 43.

Buysse, D.J., Reynolds, C.F., 3rd, Monk, T.H., Berman, S.R., Kupfer, D.J., 1989. The Pittsburgh Sleep Quality Index: a new instrument for psychiatric practice and research. Psychiatry Res. 28, 193-213.

Costa, P.T., McCrae, R.R., 1992. Revised NEO Personality Inventory (NEO-PI-R) and NEO five-factor inventory (NEO-FFI) Professional Manual. Psychological Assessment Resources, Inc, Odessa FL.

Drake, C., Richardson, G., Roehrs, T., Scofield, H., Roth, T., 2004. Vulnerability to stress-related sleep disturbance and hyperarousal. Sleep 27, 285-291.

Gros, D.F., Antony, M.M., Simms, L.J., McCabe, R.E., 2007. Psychometric properties of the State-Trait Inventory for Cognitive and Somatic Anxiety (STICSA): comparison to the State-Trait Anxiety Inventory (STAI). Psychol Assess 19, 369-381.

Horne, J.A., Ostberg, O., 1976. A self-assessment questionnaire to determine morningness-eveningness in human circadian rhythms. Int. J. Chronobiol. 4, 97-110.

Iber, C., al., e., 2007. The AASM Manual for the Scoring of Sleep and Associated Events: Rules, Terminology and Technical Specification. American Academy of Sleep Medicine, Westchester, IL.

Johns, M.W., 1991. A new method for measuring daytime sleepiness: the Epworth sleepiness scale. Sleep 14, 540-545.
Laborde, S., Mosley, E., Thayer, J.F., 2017. Heart Rate Variability and Cardiac Vagal Tone in Psychophysiological Research - Recommendations for Experiment Planning, Data Analysis, and Data Reporting. Frontiers in psychology 8, 213.
Marin, M.F., Zsido, R.G., Song, H., Lasko, N.B., Killgore, W.D.S., Rauch, S.L., Simon, N.M., Milad, M.R., 2017. Skin Conductance Responses and Neural Activations During Fear Conditioning and Extinction Recall Across Anxiety Disorders. JAMA Psychiatry 74, 622-631.
Mehling, W.E., Price, C., Daubenmier, J.J., Acree, M., Bartmess, E., Stewart, A., 2012. The Multidimensional Assessment of Interoceptive Awareness (MAIA). PLoS ONE 7, e48230.
Milad, M.R., Furtak, S.C., Greenberg, J.L., Keshaviah, A., Im, J.J., Falkenstein, M.J., Jenike, M., Rauch, S.L., Wilhelm, S., 2013. Deficits in conditioned fear extinction in obsessive-compulsive disorder and neurobiological changes in the fear circuit. JAMA Psychiatry 70, 608-618; quiz 554.
Milad, M.R., Pitman, R.K., Ellis, C.B., Gold, A.L., Shin, L.M., Lasko, N.B., Zeidan, M.A., Handwerger, K., Orr, S.P., Rauch, S.L., 2009. Neurobiological basis of failure to recall extinction memory in posttraumatic stress disorder. Biol. Psychiatry 66, 1075-1082.
Milad, M.R., Quirk, G.J., Pitman, R.K., Orr, S.P., Fischl, B., Rauch, S.L., 2007a. A role for the human dorsal anterior cingulate cortex in fear expression. Biol. Psychiatry 62, 1191-1194.
Milad, M.R., Wright, C.I., Orr, S.P., Pitman, R.K., Quirk, G.J., Rauch, S.L., 2007b. Recall of fear extinction in humans activates the ventromedial prefrontal cortex and hippocampus in concert. Biol. Psychiatry 62, 446-454.
Orr, S.P., Lasko, N.B., Shalev, A.Y., Pitman, R.K., 1995. Physiologic responses to loud tones in Vietnam veterans with posttraumatic stress disorder. J. Abnorm. Psychol. 104, 75-82.
Orr, S.P., Metzger, L.J., Lasko, N.B., Macklin, M.L., Peri, T., Pitman, R.K., 2000. De novo conditioning in trauma-exposed individuals with and without posttraumatic stress disorder. J. Abnorm. Psychol. 109, 290-298.
Pace-Schott, E.F., Kaji, J., Stickgold, R., Hobson, J.A., 1994. Nightcap measurement of sleep quality in self-described good and poor sleepers. Sleep 17, 688-692.
Pace-Schott, E.F., Spencer, R.M., Vijayakumar, S., Ahmed, N.A., Verga, P.W., Orr, S.P., Pitman, R.K., Milad, M.R., 2013. Extinction of conditioned fear is better learned and recalled in the morning than in the evening. J. Psychiatr. Res. 47, 1776-1784.
Pace-Schott, E.F., Stickgold, R., Muzur, A., Wigren, P.E., Ward, A.S., Hart, C.L., Clarke, D., Morgan, A., Hobson, J.A., 2005. Sleep quality deteriorates over a binge--abstinence cycle in chronic smoked cocaine users. Psychopharmacology (Berl). 179, 873-883.
Rechtschaffen, A., Kales, A., 1968. A Manual of Standardized Terminology Techniques and Scoring System for Sleep Stages of Human Subjects. University of California at Los Angeles, Brain Information Service/Brain Research Institute.

Ree, M.J., French, D., MacLeod, C., Locke, V., 2008. Distinguishing Cognitive and Somatic Dimensions of State and Trait Anxiety: Development and Validation of the State-Trait Inventory for Cognitive and Somatic Anxiety (STICSA). Behavioural and Cognitive Psychotherapy 36, 313–332.
Ree, M.J., MacLeod, C., French, D., Locke, V., 2000. State–Trait Inventory for Cognitive and Somatic Anxiety (STICSA)—Trait Version. The University of Western Australia, Perth, Australia.
Reiss, S., Peterson, R.A., Gursky, D.M., McNally, R.J., 1986. Anxiety sensitivity, anxiety frequency and the prediction of fearfulness. Behav. Res. Ther. 24, 1-8.
Rush, A.J., Trivedi, M.H., Ibrahim, H.M., Carmody, T.J., Arnow, B., Klein, D.N., Markowitz, J.C., Ninan, P.T., Kornstein, S., Manber, R., Thase, M.E., Kocsis, J.H., Keller, M.B., 2003. The 16-Item Quick Inventory of Depressive Symptomatology (QIDS), clinician rating (QIDS-C), and self-report (QIDS-SR): a psychometric evaluation in patients with chronic major depression. Biol. Psychiatry 54, 573-583.
Seo, J., Moore, K.N., Gazecki, S., Bottary, R.M., Milad, M.R., Song, H., Pace-Schott, E.F., 2018. Delayed fear extinction in individuals with insomnia disorder. Sleep 41.
Seo, J., Oliver, K.I., Daffre, C., Moore, K.N., Gazecki, S., Lasko, N.B., Milad, M.R., Pace-Schott, E.F., 2022a. Associations of sleep measures with neural activations accompanying fear conditioning and extinction learning and memory in trauma-exposed individuals. Sleep 45.
Seo, J., Oliver, K.I., Daffre, C., Moore, K.N., Gazecki, S., Lasko, N.B., Milad, M.R., Pace-Schott, E.F., 2022b. Associations of sleep measures with neural activations accompanying fear conditioning and extinction learning and memory in trauma-exposed individuals. Sleep in press.
Seo, J., Pace-Schott, E.F., Milad, M.R., Song, H., Germain, A., 2020. Partial and Total Sleep Deprivation Interferes with Neural Correlates of Consolidation of Fear Extinction Memory. Biological Psychiatry: Cognitive Neuroscience and Neuroimaging (in press).
Shaffer, F., Ginsberg, J.P., 2017. An Overview of Heart Rate Variability Metrics and Norms. Front Public Health 5, 258.
Spielberger, C.D., Gorsuch, R.L., Lushene, R.E., 1990. Manual for the State-Trait Anxiety Inventory (Self-Evaluation Questionnaire). Consulting Psychologists Press, Palo Alto.
Spitzer, R.L., Kroenke, K., Williams, J.B., Lowe, B., 2006. A brief measure for assessing generalized anxiety disorder: the GAD-7. Arch. Intern. Med. 166, 1092-1097.
Yuksel, C., Watford, L., Muranaka, M., Daffre, C., McCoy, E., Lax, H., Mendelsohn, A.K., Oliver, K.I., Acosta, A., Vidrin, A., Martinez, U., Lasko, N., Orr, S., Pace-Schott, E.F., 2024. REM disruption and REM vagal activity predict extinction recall in trauma-exposed individuals. Psychol. Med. 54, 1-12.
